# Modulation of FGF pathway signaling and vascular differentiation using designed oligomeric assemblies

**DOI:** 10.1101/2023.03.14.532666

**Authors:** Natasha I Edman, Rachel L Redler, Ashish Phal, Thomas Schlichthaerle, Sanjay R Srivatsan, Ali Etemadi, Seong J An, Andrew Favor, Devon Ehnes, Zhe Li, Florian Praetorius, Max Gordon, Wei Yang, Brian Coventry, Derrick R. Hicks, Longxing Cao, Neville Bethel, Piper Heine, Analisa Murray, Stacey Gerben, Lauren Carter, Marcos Miranda, Babak Negahdari, Sangwon Lee, Cole Trapnell, Lance Stewart, Damian C. Ekiert, Joseph Schlessinger, Jay Shendure, Gira Bhabha, Hannele Ruohola-Baker, David Baker

## Abstract

Growth factors and cytokines signal by binding to the extracellular domains of their receptors and drive association and transphosphorylation of the receptor intracellular tyrosine kinase domains, initiating downstream signaling cascades. To enable systematic exploration of how receptor valency and geometry affects signaling outcomes, we designed cyclic homo-oligomers with up to 8 subunits using repeat protein building blocks that can be modularly extended. By incorporating a *de novo* designed fibroblast growth-factor receptor (FGFR) binding module into these scaffolds, we generated a series of synthetic signaling ligands that exhibit potent valency- and geometry-dependent Ca2+ release and MAPK pathway activation. The high specificity of the designed agonists reveal distinct roles for two FGFR splice variants in driving endothelial and mesenchymal cell fates during early vascular development. The ability to incorporate receptor binding domains and repeat extensions in a modular fashion makes our designed scaffolds broadly useful for probing and manipulating cellular signaling pathways.

**Highlights:**

- De novo designed cyclic oligomers with tunable geometric properties
- Cyclic, homo-oligomeric FGFR binding modules induce geometry- and valency-dependent activity of isoform-specific FGF signaling
- Modulation of FGFR isoform activity controls bifurcation of endothelial and mesenchymal fate during vascular development
- C-isoform activation favors arterial endothelial cell formation while B-isoform induces pericyte differentiation

**Graphical Abstract:** 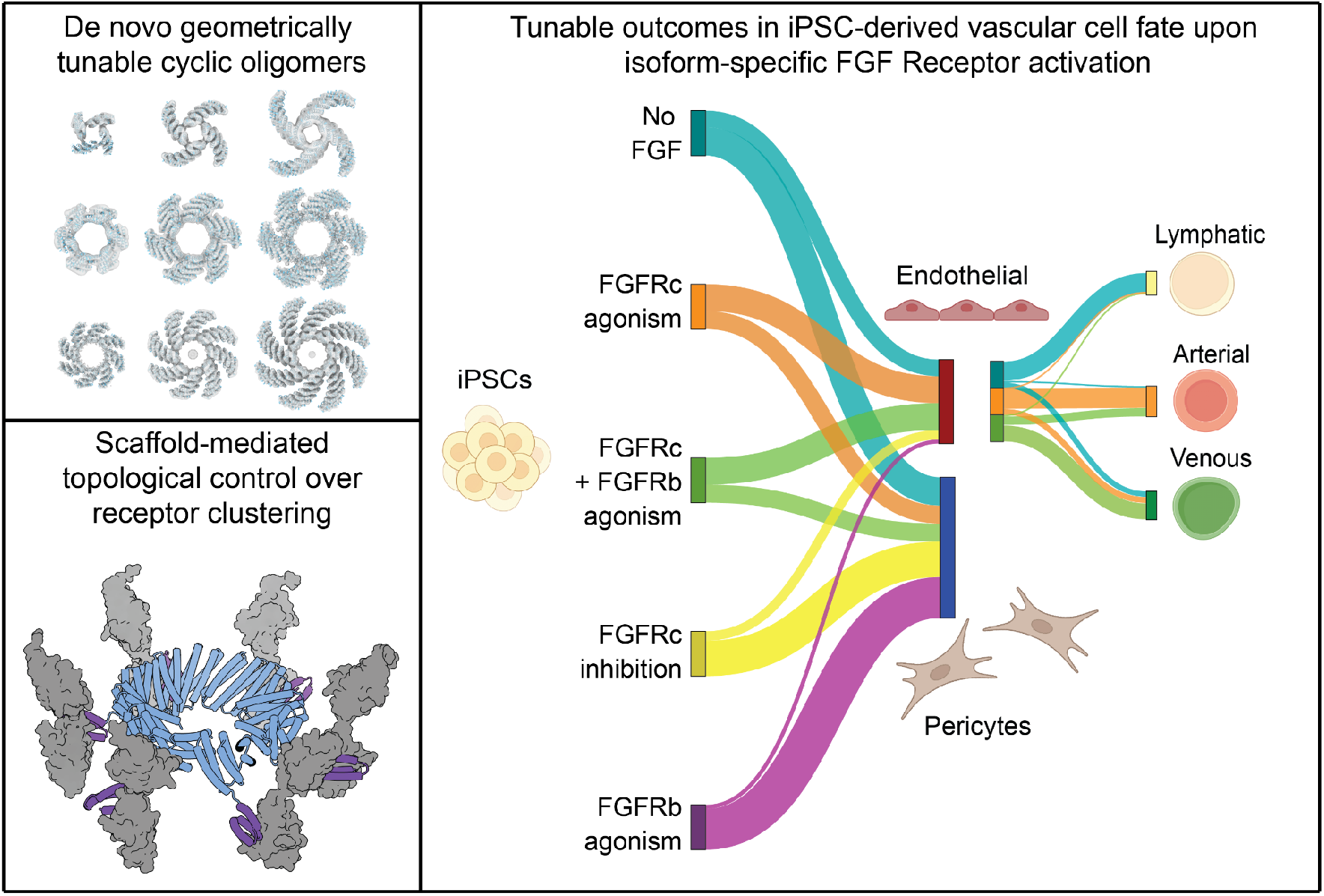

## Introduction

Clustering of cell surface receptors can enhance and sustain activation in response to an extracellular signal, and there is considerable interest in technologies to manipulate receptor clustering.^1–5^ Designed protein assemblies have been used to drive receptor clustering;^6–8^ particularly useful are oligomers made from idealized repeat proteins because their length can be systematically varied by addition of repeat units.^9^ Previous work has largely focused on designs that bring together two receptor subunits,^10–13^ but higher order receptor assemblies are thought to function in a number of signaling systems.^14–16^ Previous design efforts have not generated geometrically tunable assemblies higher than 5-fold cyclic (C5) symmetry, highlighting the need for methods to design synthetic ligands that can drive association of higher order receptor assemblies.^17^ To enable systematic probing of the physiological effects of clustering receptors at higher valencies and different spacings, we set out to design repeat protein homo-oligomers with 2- to 8-fold cyclic symmetry. We combined these oligomers with a *de novo* designed binder^18^ against the fibroblast growth factor receptor 2 (FGFR2).

FGF receptors are tyrosine kinases that play critical roles in vascular development and in cancer. The pathway is complex and highly regulated with four FGF receptor genes and two isoforms (Ig-like domain IIIb and IIIc; we refer to these as “b” and “c” throughout the remainder of the text) generated by alternative splicing (exon 8 vs exon 9) that produces the C-terminus of the third Ig-like domain (D3) which is part of the FGF binding region.^19–21^ How this complexity mediates proper tissue differentiation is not fully understood. FGFR amplification has been observed in many solid carcinomas; the c splice variant is predominantly enriched in tumors, indicating that this isoform may be a druggable target for cancer therapy.^22^ While FGF signaling is critical in endothelial and mesenchymal branches of vascular development, its contribution to the bifurcation process is not clear.

Here we describe the *de novo* design of geometrically tunable cyclic oligomers, and the use of these synthetic scaffolds with a FGFRIIIc isoform specific designed minibinder to probe and manipulate vascular differentiation.

## Results

### De novo oligomer design

Cyclic oligomers (Cx, with “x” denoting valency) were designed using a set of 18 designed helical repeat proteins (DHRs), each consisting of four identical repeats of a two helix module and for which high-resolution crystal structures or small-angle X-ray (SAXS)^23–25^ spectra showed close agreement with the respective design models (**Supplementary Table I, Supplementary Figure 1**). We docked each DHR into C4, C5, C6, C7 and C8 cyclic oligomeric assemblies and evaluated them using the protein backbone based residue-pair transform (RPX) metric, which assesses interface designability.^26^ For the top scoring docks, the residue identities and conformations at the homo-oligomeric interface were optimized using RosettaDesign to favor oligomer assembly. We filtered for designs with high solvent accessible surface area (SASA >700 Å^2^), favorable free energies of assembly (∆∆G between -35 and -70), high shape complementary (sc > 0.65), and interfaces with fewer than 2 unsatisfied hydrogen bonds.^27,28^ A total of 109 designs were selected for structural characterization: 15 tetramers, 16 pentamers, 24 hexamers, 24 heptamers, and 30 octamers. A second set of designs using a computational library of 1526 “*junior helical repeat proteins*” (JHRs; manuscript in preparation)^29^ were docked into C2 symmetry and from 3747 C2 oligomers, 14 designs were selected for further analysis (**Supplementary Figure 2**).

### Design characterization

Synthetic genes encoding the 109 designs of symmetry C4 or higher were synthesized, expressed as protein in *Escherichia coli*, and purified using immobilized metal affinity chromatography (IMAC). Of the 60 designs that were soluble, 28 had single monodisperse peaks on size exclusion chromatography (SEC). Of these, ten designs were found to have a single oligomeric state by both SAXS and SEC-MALS. Five of the successes were tetramers, four were hexamers, and one was an octamer. From the 14 C2 designs, three (C2-58, C2-CDX, C2-Y2D) had soluble expression, were confirmed as a monodisperse peak on SEC and had a correctly assembled oligomeric state verified by SAXS and SEC-MALS (**Supplementary Figure 3**).

The varied topology of the repeat protein building blocks enabled us to create oligomers with distinct arm orientations. The starting scaffold DHR71 generated 5 successful designs (C4-71, C4-717, C6-71, C6-714, and C8-71), with a variety of interface geometries that permitted this building block to assume 3 distinct valencies. C4-71 and C4-717, for example, contain changes in different sets of residues that result in distinct oligomer geometries. In contrast, the designs C4-71, C6-71, and C8-71 employ a similar backbone region as the oligomeric interface, yet adopt different oligomeric states (**Supplementary Figure 4-5**). C4-181 utilizes DHR18 as the single chain building block and is docked together at the C-terminal helices yielding an inner cavity diameter of 45.6 Å (C-terminal distance of opposing chains, **Figure 1A**). C4-717 is tightly docked together at the C-terminal helices creating a purely hydrophobic core between all four chains (**Figure 1B**). C6-714 has an inner cavity diameter of 43.2 Å and its N-terminus can be extended to achieve larger distance spacing, whereas the structure is again docked together at the C-terminus (**Figure 1C**). C6-46 involves the carboxyl- (C) and amino- (N) terminal helices at the interfaces to adjacent chains, where the N-terminus points towards the central cavity and the C-terminus towards the outside (**Figure 1D**). Six designs were further selected for characterization by cryo-electron microscopy (cryo-EM) (**Supplementary Table II and III**). The cryo-EM map for windmill-shaped C4-131 was limited to >10 Å global resolution due to preferred orientation bias, but shows that the “blades” are arranged as designed and that the four core C-terminal helices are tightly packed (**Figure 1E**). The global resolution of design C4-814 allows individual helices to be clearly distinguished, and rigid-body fitting using ChimeraX of the design model to the cryo-EM map shows good agreement (**Figure 1F**). C6-79 again involved N- and C-terminal helices for docking to adjacent chains; however, it preferentially formed a hexamer instead of the designed octamer. Accordingly, a C6 predicted SAXS trace matched the experimental data more closely than the original C8 predicted trace. Using the same lowest energy predicted C6 dock of C6-79 as was employed in SAXS analysis, we found that the cryo-EM map for C6-79 closely matched the C6 dock, and the 2D classes clearly indicate that it is a hexamer under our cryo-EM conditions (**Figure 1G**).

**Figure 1:**
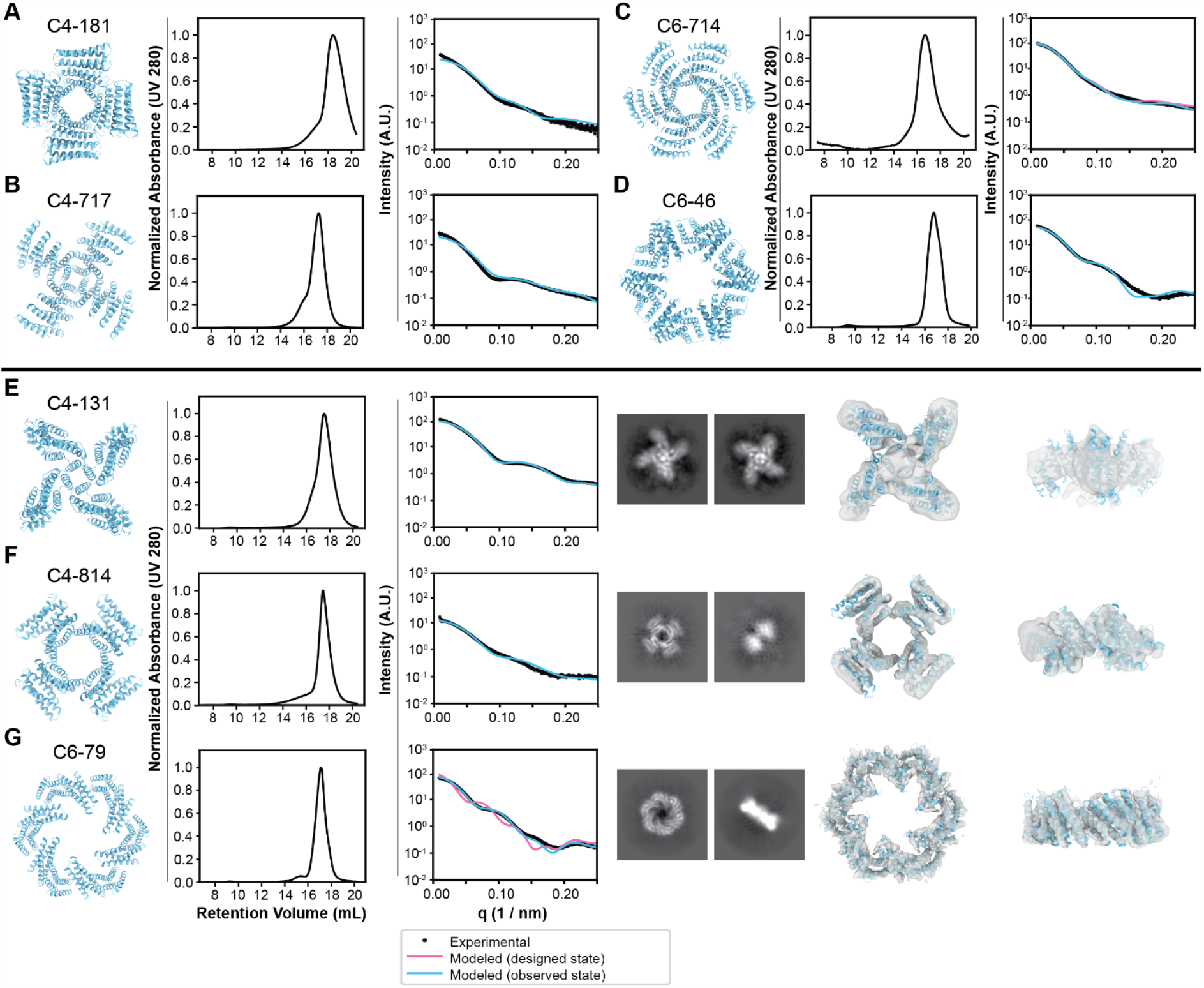
Biophysical characterization of designed protein oligomers. From left to right: design model, size-exclusion chromatogram, SAXS data comparison of model to experimental data (A) C4-181, (B) C4-717, (C) C6-714, (D) C6-46, (E) C4-131 design model, size-exclusion chromatogram, SAXS data analysis, *Right:* cryo-EM 2D class average, cryo-EM map overlay to design model (cyan) top and side view (F) C4-814 design model, size-exclusion chromatogram, SAXS data analysis, *Right:* cryo-EM 2D class average, cryo-EM map superimposed to design model (cyan) top and side view (G) C6-79 SEC characterization and SAXS fit using both the C8 design model and the C6 dock. *Right*: cryo-EM 2D class average, cryo-EM map superimposed to design model top and side view.

### Oligomer extension

An advantage of using modular repeat proteins as building blocks is that the length of the oligomer arms can be increased or decreased simply by inserting or deleting repeat units (**Figure 2A**).^9,10,17^ To explore the viability of this approach, three designs (C4-71, C6-71, C8-71) derived from DHR71 were selected for repeat extension. Two or four repeat units were added at the N-terminus, creating a 6-repeat variant and an 8-repeat variant of each design. The oligomeric state of each extended design was characterized by SEC-MALS, SAXS, and cryo-electron microscopy. Both 2D classes and 3D reconstructions from single-particle cryo-EM analysis of the extended oligomers show overall geometry in good agreement with design models, with sufficiently high resolution in some cases to confirm positions of individual helices. The C-terminal helix of C4-71 docked as designed against the mid-axis of the neighboring chain horizontally, yielding an inner cavity distance of 47.4 Å between opposing chain C-termini (**Figure 2B**). The interface was designed harboring 10 tryptophans allowing for pi-pi stacking interactions to stabilize the C4 symmetric complex.

**Figure 2:**
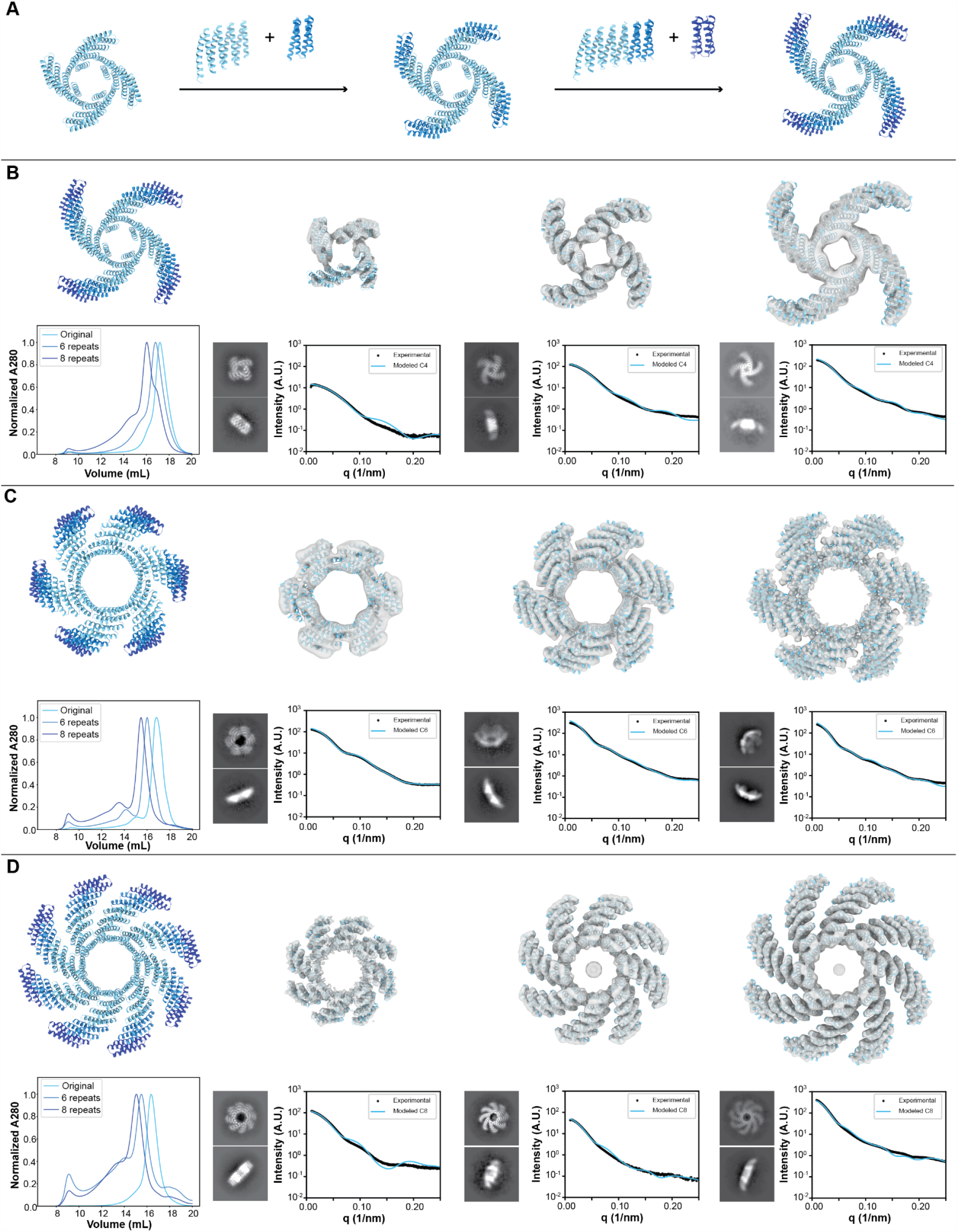
Repeat extensions of designed oligomers. (A) Depiction of DHR-based repeat extension for oligomers. Each extension unit consists of 2 repeats. (B) C4-71 4-repeat, 6-repeat and 8-repeat cryo-EM maps superimposed with design model, top and side-view class averages and SAXS characterization below the cryo-EM maps of the different repeat extension variants. *Bottom Left:* SEC overlay of the individual structures. (C) C6-71 4-repeat, 6-repeat and 8-repeat cryo-EM maps superimposed with design model, top and side-view class averages and SAXS characterization below the cryo-EM maps of the different repeat extension variants. *Bottom Left:* SEC overlay of the individual structures. (D) C8-71 4-repeat, 6-repeat and 8-repeat cryo-EM maps superimposed with design model, top and side-view class averages and SAXS characterization below the cryo-EM maps of the different repeat extension variants. *Bottom Left:* SEC overlay of the individual structures.

C6-71, in contrast, has an inner diameter of 72.0 Å between opposing chain C-termini and harbors a tilted chain-chain interaction, where the interfacial C-terminal helix is only in contact with the neighboring chain along half its length. The C6-71 8-repeat extension map in particular contains sufficient detail to hint at side chain orientation, remarkable given the low number of total particles used in constructing this map (**Figure 2C**). The octopus-like C8-71 structure has N-terminal extensible arms with C-terminal helices of the individual chains docked together along the full horizontal length of the structure. This arrangement yields an inner diameter of 55.1 Å and a maximal distance between opposing N-termini of 170.0 Å in the largest 8-repeat extension (**Figure 2D**).

All cryo-EM maps were in good agreement with the respective design models, with the exception of C6-79, which as noted above formed a hexamer instead of the designed octamer. None of the other designs showed any off-target oligomeric states in the 2D class averages (**Supplementary Figure 6-17**).

### Cryo-electron microscopy reconstructions of C6-79 and C8-71

Based on the resolution of the cryo-EM maps, we built models for C6-79 and C8-71 (**Supplementary Figure 18-20, Supplementary Table IV**). Both the C6-79 and C8-71 cryo-EM models align well with the corresponding design models, with pairwise root mean squared deviations (RMSDs) of 2.85 Å and 1.79 Å, respectively (**Figure 3**). In C8-71, the hydrophobic residues Trp152 and Leu198 on the adjacent chain are buried in the interface or the core of the structure respectively and are important for interface formation (**Supplementary Figure 21A**). Mutating these residues to hydrophilic residues (W152E and/or L198D) disrupts oligomer formation as shown by broadening of the SEC trace (**Supplementary Figure 21B**).

**Figure 3:**
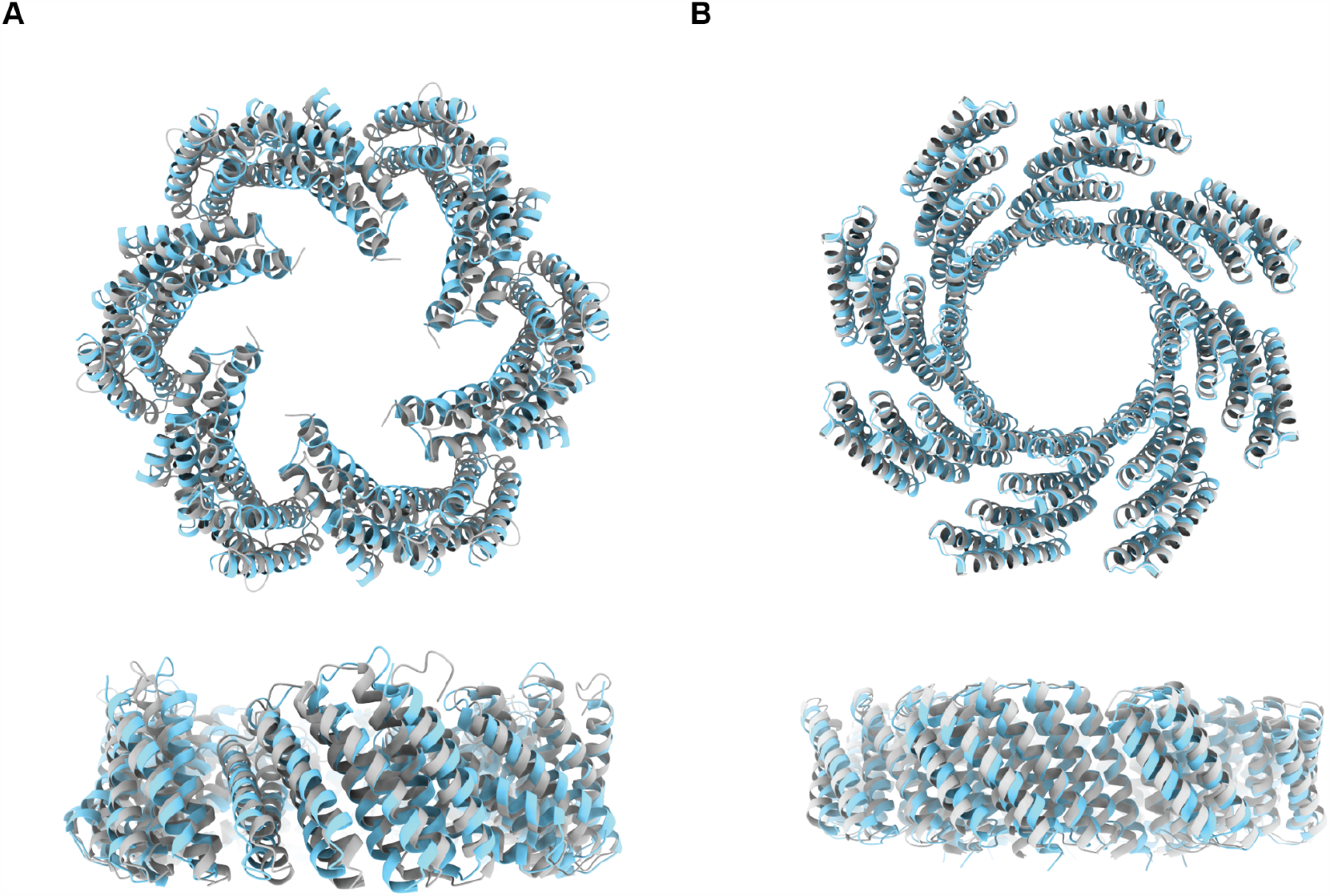
Cryo-EM structural analysis. (A) C6-79 alignment of design model (grey) with cryo-EM structure (cyan) in top and side view. Structures align well with an RMSD of 2.85 Å (B) C8-71 alignment of design model (grey) with cryo-EM structure (cyan) in top and side view. Structures are in good agreement with an RMSD of 1.79 Å.

### Design of FGFR agonists

We next investigated whether clustering receptor tyrosine kinases in higher-order geometries by presenting receptor binding domains on the designed oligomers could drive cross-phosphorylation of their intracellular kinase domains and induce downstream signalling.^30^ The multiple distinct valencies and geometries of our oligomeric ligands enable exploration of how the geometry and valency of tyrosine kinase receptor association influences signaling output and cell behavior (**Figure 4A, left**). We chose as a model system the FGF signaling pathway (**Figure 4A, right**), and fused a *de novo* designed minibinder (mb7) against the FGFR2 receptor at either the N- or C-termini of the designed cyclic oligomers with a short glycine-serine linker.^18^ Six oligomers were selected for fusion: C2-58, C4-71, C6-71, C6-79 and C8-71. Depending on the fusion terminus and the geometry of the oligomer, the binding domains are displayed at different spacings on adjacent subunits: for example, C6-79C_mb7 displays the minibinders 54 Å apart with mb7 on the C terminus of the oligomer, while C6-79N_mb7 displays the binders 18 Å apart with mb7 on the N terminus of the oligomer. The fusions eluted at the same volume as the oligomers determined by SEC, with the exception of C6-71C_mb7, which eluted significantly earlier than the base design. 2D EM class averages showed that C6-71C_mb7 particles were self-associating into dihedral structures, presumably via the hydrophobic interface of the minibinder domain being presented in a favorable conformation for this interaction. The other oligomeric fusions showed little to no self-association on EM or SEC (**Supplementary Figure 22**).

**Figure 4:**
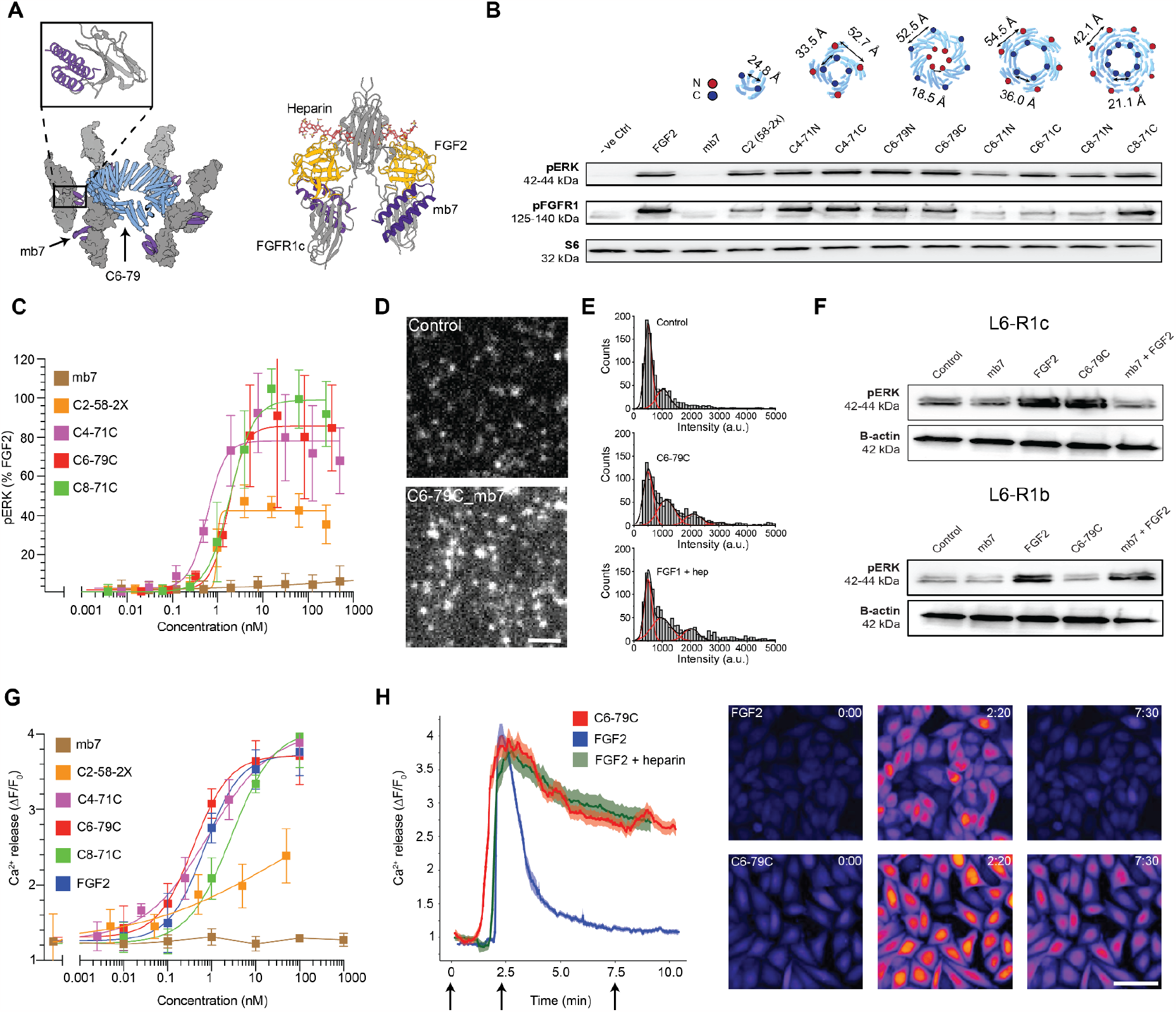
Modulation of FGFR signaling by designed agonists. (A) Cartoon model of C6-79C_mb7 oligomer (blue and purple) engaging six FGFR2 receptors (grey). Top left: Cartoon model of mb7 engaging FGFR4 domain 3 (pdb ID: 7N1J). Right: Natural geometry of signaling competent FGF2 (orange) with FGFR1c (grey) and heparin (red) (pdb ID: 1FQ9) together with superimposed mb7 (purple). (B) Signaling response to a library of oligomers presenting mb7 in CHO-R1c cells, analyzed through western blot. *Top:* Cartoons of oligomers presenting mb7 at their N- or C-termini; distances between neighboring chains are shown above their respective treatments. Total FGFR1 and ERK loading controls can be found in Supplementary Figure 23. (C) Dose-response curves of selected designs via phosphoflow for pERK1/2 stimulation. Error bars represent SEM from three independent biological repeats. (D) Single-particle tracking of FGFR1 receptors on the cell surface. (E) Intensity histograms of receptor clusters on the cell surface reveals receptor clustering induced via oligomerization. (F) Signaling response to FGF2, mb7, C6-79C_mb7 or mb7 + FGF2 in L6-R1c (*top*) or L6-R1b (*bottom*) cells (G) Dose-response curves of selected designs through intracellular calcium release. Error bars represent SEM from three independent biological repeats. (H) Comparison of a calcium intensity signaling trajectory after treatment with FGF2 (with or without heparin) or C6-79C_mb7 at 10 nM each. *Right*: Exemplary images comparing the calcium response exhibited in CHO-R1c cells following treatment with FGF2 or C6-79C_mb7 at 10 nM over three different timepoints (0:00, 2:20 and 7:30 min). Scale Bars: 2 µm (D), 66.3 µm (H).

### FGFR pathway activation

FGF-mediated FGFR signaling has several downstream effectors, including stimulation of the Ras signaling pathway leading to phosphorylation of extracellular signal-regulated kinase 1 and 2 (ERK1/2) and activation of phospholipase C-gamma (PLC-ɣ) leading to intracellular calcium release^31–33^. We evaluated the signaling activity of our designs by screening them in serum-starved CHO cells stably expressing hFGFR1c (CHO-R1c) at 10 nM each for 15 min at 37°C. Downstream activation through phosphorylation of ERK1/2 and the FGF receptor (Y653/654) was analyzed by western blot. Of the designs, we found that C6-79C_mb7, C6-79N_mb7, C4-71N_mb7, C4-71C_mb7 and C8-71C_mb7 broadly induce strong FGFR activation and ERK1/2 phosphorylation comparable to that achieved by native FGF2, while C2-58-2X_mb7, C6-71C_mb7, C6-71N_mb7, and C8-71N_mb7 displayed weaker activity (**Figure 4B, Supplementary Figure 23**).

To characterize their dose-dependent activity, we titrated a subset of these designs using phosphoflow^34^ and western blotting for ERK1/2 phosphorylation in CHO-R1c cells (**Figure 4C, Supplementary Figure 24**). C2-58-2X_mb7, C4-71C_mb7, C4-71N_mb7, C6-79C_mb7 and C8-71C_mb7 had similar EC_50_ values of 0.63 nM, 1.33 nM, 0.89 nM, 1.56 nM and 2.07 nM respectively, and similar maximal activation (Emax) values, while C2-58-2X_mb7 had a lower Emax. To investigate how the geometry of receptor association influences signaling, the rigid repeat arm length of C4-71N_mb7 was systematically varied, leading to distances between mb7 N-termini of 53 Å, 76 Å and 96 Å. Phosphoflow experiments showed that only the shortest separation distance (53 Å) was able to stimulate ERK phosphorylation (with an EC_50_ of 1.3 nM), whereas the larger separation distances of mb7 did not lead to pathway activation (**Supplementary Figure 25**).

### Receptor clustering on the cell surface

To investigate whether FGFR1c activation was due to induced receptor clustering, cells were examined by single-particle tracking with a HaloTag targeting FGFR1c^35^ to directly visualize their diffusion in the plasma membrane; receptors engaged in a signaling cluster should exhibit decreased diffusion manifesting in a decreased diffusion coefficient.^36^ Receptors on cells treated with C6-79C_mb7 showed slower diffusion than those treated with FGF1 and heparin (**Supplementary Figure 26**), indicating that C6-79C_mb7 induces an oligomeric state of FGFR1c at the membrane. To probe the presence of local receptor clusters on the cell surface after ligand treatment, intensity levels of single spots in HaloTagged CHO-R1c cells labeled with Alexa488 were evaluated.^37^ C6-79C_mb7 treated cells showed signals with an intensity distribution slightly shifted compared to FGF1 supplemented with heparin, with intensity peaks at 500, 1000, and 2000 a.u, suggesting that multiple receptors are clustered together by the designed mb7 presenting oligomers. The extent of signaling correlated with the ability of the designs to cluster receptors (**Figure 4D and E**).

### FGFR1c isoform specificity

FGFRs 1-3 have two alternatively spliced variants, the “c” and “b” isoforms, which have different third Ig-like domains and variable ligand affinities.^38^ Tissue-specific expression of these isoforms and their reciprocal signaling play roles in embryonic development, tissue repair, and cancer.^20^ Separating the functions of the FGFR b- and c-isoforms in differentiation has been hindered by a lack of ligands that can selectively bind one isoform or the other. The mb7 minibinder was designed to specifically bind the c-isoform of the FGF receptor, and it selectively inhibits signaling through this isoform.^19^ We evaluated the receptor isoform specificity of our synthetic agonists by treating serum-starved L6 rat myoblast cells stably expressing either the c- or b-isoform of hFGFR1 (L6-R1c or L6-R1b, respectively) with 10 nM of mb7, FGF2, or C6-79C_mb7 for 15 min at 37°C. Overexpression of each cell line’s respective FGFR splice variant was validated with RT-qPCR **(Supplementary Figure 27)**. While FGF2 does not discriminate between the two FGFR1 isoforms and activates signaling in both cell types, C6-79C_mb7 stimulates ERK1/2 phosphorylation in L6-R1c cells only, and is inactive in L6-R1b. We reasoned that it should be possible to specifically activate signaling through the b subunit by combining FGF with the monomeric mb7 (which blocks signaling through the c subunit); to test this we stimulated both L6 cell lines with a combination of mb7 and FGF2 at 10 nM each for 15 minutes. We found that this combination stimulates ERK1/2 phosphorylation in L6-R1b cells only; thus our designs enable selective activation of signaling through either isoform (**Figure 4F)**.

We investigated the ability of the designs to activate FGF signaling through the PLC-ɣ downstream branch of signaling by measuring the levels of intracellular calcium release following treatment of serum-starved CHO-R1c cells with varying concentrations of the designs. These results show a similar trend: C6-79C_mb7, C4-71C_mb7 and C8-71C_mb7 induce strong intracellular calcium release with EC_50_ values of 0.38 nM, 0.72 nM and 3.09 nM respectively, while C2-58-2X_mb7 displays lower activity with an EC_50_ of 26.02 nM. (**Figure 4G, Supplementary Figures 28-31**). While the peak magnitude of calcium release was similar between FGF2 at 10 nM and the synthetic agonist C6-79C_mb7 at 10 nM, there was a pronounced difference in the duration of the response: the higher valency synthetic ligand, C6-79C_mb7, generated longer duration calcium transients (**Figure 4H**), similar to a control condition in which we supplemented FGF2 together with heparin. This shows the strong, heparin-independent signaling effect (**Supplementary Figure 32**) of our designed agonist and likely reflects the slow off rates of the high avidity multivalent agonists (**Supplementary Figure 33**).

### Sculpting vascular differentiation with the designed agonists

FGF signaling plays an important role during early embryogenesis;^39^ the controlled spatio-temporal expression of FGF receptors and their ligands drives specification and development of many cell lineages. Vascular development is dependent on FGF signaling and sustained VEGF and FGF signaling are critical for the development and maintenance of mature endothelial cells.^40,41^ Additionally, FGF is a key driver for both endothelial and mesenchymal vascular cell fates, and the c-isoform of FGFRs 1-3 is highly expressed in endothelial cells over mesenchymal pericytes; though the bifurcation between endothelial and pericyte fate is poorly understood and it is not clear whether FGFR c-isoform activity plays a role in this event.^42,43^ We investigated the effect of the c-isoform specific FGFR minibinder oligomers on vascular development by generating iPSC derived endothelial cells and pericytes through a cardiogenic mesoderm intermediate.^44^ This method employs a differentiation media containing FGF2, which engages both b- and c-isoforms of FGFR. The specificity of the mb7-based designs allows us to selectively engage FGFR b- or c-isoforms and explore their effects on vascular tissue formation. We replaced the 1 nM FGF2 in the differentiation media at day 2 (when mesodermal intermediates first appear) in the protocol with either 1 nM

C6-79C_mb7, 100 nM C2-58-2X_mb7, 10 nM mb7, or 10 nM mb7 in combination with 1 nM FGF2 (to specifically activate signaling through the b-receptor isoform) and allowed the cells to differentiate for 28 days; samples were harvested for scRNAseq analysis at days 0, 5, 14 and 28 **(Supplementary Figure 34A)**. The sequencing datasets were analyzed using Monocle3^45^ and visualized using uniform manifold approximation and projection (UMAP), which revealed 5 clusters of cells that segregated predominantly by time point and cell type (**Figure 5A)**; cell types were annotated based on the expression of previously published canonical marker genes; **Supplementary Figure 34B**).

**Figure 5:**
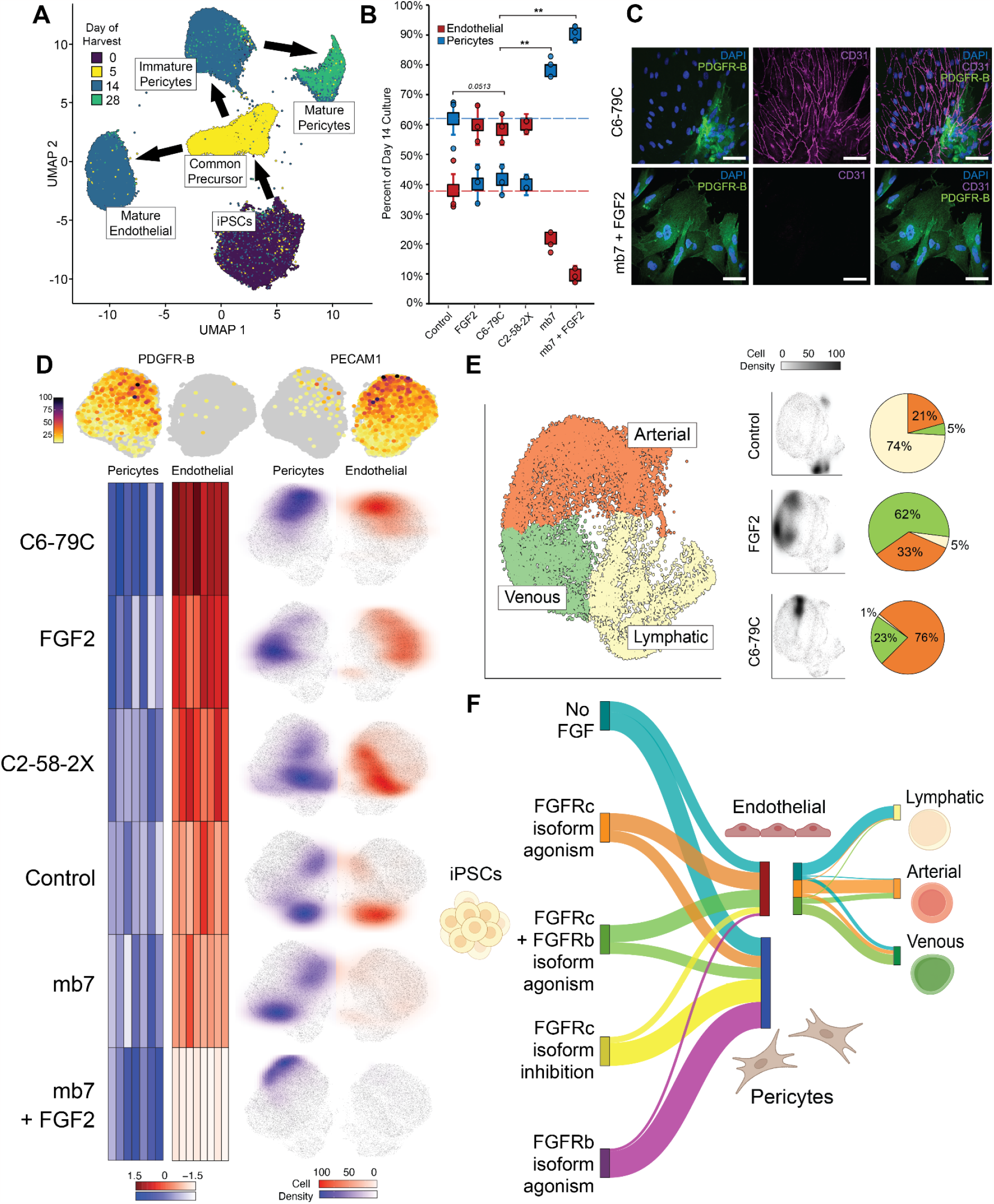
Control over vascular differentiation with designed agonists and inhibitors. (A) UMAP graph of all sequenced cells colored by day of harvest, along with given cluster annotations. (B) Proportion of endothelial and pericyte cells generated at day 14 following treatment with FGF2, C2-58-2X_mb7, C6-79C_mb7, mb7 alone, or mb7 in combination with FGF2. Error bars represent SEM from three independent biological repeats. (C) Representative immunofluorescence staining of differentiated cells treated with C6-79C_mb7 or mb7 in combination with FGF2, with anti-PDGFR-B and anti-CD31 to specifically mark pericytes and endothelial cells, respectively. Scale bar: 20 µm. (D) Clustered heatmap comparing the normalized average expression of selected endothelial and pericyte cell maturity markers across all analyzed treatment conditions. Marker gene names in the heatmap (left to right) can be found in Supplementary Table VII in the endothelial or pericyte rows (top to bottom) respectively. The cell density plots show specific cell populations enriched by the individual treatments. (E) Proportion of arterial, lymphatic or venous endothelial cells generated at day 14 following treatment with FGF2 or C6-79C_mb7. (F) Control over vascular differentiation. At the first bifurcation, the designs enable selective formation of endothelial cells or pericytes, and in subsequent endothelial cell differentiation, synthetic agonist treatments bias towards arterial fate.

All treatments (FGF2 and designed agonists) directed iPSCs at day 0 to differentiate and form a common endothelial-mesenchymal precursor at day 5. This common precursor population then bifurcated to form either mature endothelial cells at day 14, or mesenchymal vascular pericytes that continued to mature until day 28. The cellular differentiation trajectory was design-dependent and determined by day 14. Inclusion of FGF2, C6-79C_mb7, or C2-58-2X_mb7 generated roughly 60% mature endothelial cells in all three cases; the remaining population differentiated into pericytes. In contrast, the differentiation media without any FGF addition (control) resulted in a population that was only 38% endothelial (endothelial cell formation is weakly driven in the absence of any supplemented FGF2, presumably because of low levels of endogenously expressed FGFs; **Supplementary Figure 34C)**. On the other end of the spectrum, cells treated with mb7 showed a marked preference for pericyte formation, producing only 22% endothelial cells. Furthermore, cells treated with a combination of mb7 and FGF2 were almost exclusively mesenchymal, producing a population that was 90% pericytes (**Figure 5B)**. These results suggest that FGFR c-isoform activity is important for the development of mature endothelial cells, and overactivation of the b-isoform instead biases the cells towards pericyte fate. Immunostainings of differentiated iPSCs for endothelial (CD31) and pericyte (PDGFR-B) markers at day 14 confirmed the primary cell fate after treatment with C6-79C_mb7 (FGFRc - isoform signaling) or mb7 together with FGF2 (FGFRb - isoform specific signaling), which led to enrichment of endothelial cells or pericytes, respectively (**Figure 5C**).

We next investigated the maturity of the cell types generated across all the conditions tested. We compared the normalized average expression of a panel of known endothelial and pericyte maturity markers (**Supplementary Table VII**) across all treatments, and found that the endothelial cells generated by C6-79C_mb7 were the most mature, followed by FGF2, and C2-58-2X_mb7. Pericytes generated via b-isoform specific FGFR activation (mb7 in combination with FGF2) were also highly mature. Cells differentiated with the differentiation media without any FGF ligand addition were the least mature. Density plots generated for each treatment show a clear movement from the top to bottom of each cell type’s respective UMAP cluster as their maturity decreases (**Figure 5D**).

Sub-clustering and subsequent analysis of the day 14 endothelial expression data suggested that arterial, venous and lymphatic endothelial cells were generated in the differentiation experiments in different ratios with the different treatments.^46^ Endothelial cells generated without added FGF or C6-79C_mb7 agonist primarily adopted the lymphatic cell fate (74% lymphatic), while C6-79C_mb7 induced a strong bias towards an arterial-like endothelial cell fate (76% arterial-like), and FGF2 prompted the venous cell fate (66% venous-like) (**Figure 5E)**. These results highlight the potential use of designed proteins as tailored agonists for differentiation of cells into highly specific lineages.

## Discussion

The extensible star shaped oligomers designed in this work considerably expand the tools available for clustering cell surface receptors and other targets with different valencies and geometries. The designed scaffolds are highly expressed in *E. coli* and the spacing of attached binding domains can be systematically varied simply by adding or deleting the modular repeat units. C8-71 and its extensions are the first structures offering both a defined octameric symmetry and a stepwise variation in diameter through repeat units. The highest success rate was achieved with the DHR71 building block, perhaps because the design model (used in the docking protocol to avoid issues of missing terminal residues or imperfect repeat unit symmetry in the crystal structure) was closer to the crystal structure (0.67 Å RMSD)^9^ leading to greater accuracy of the oligomer computational models.

FGFR homodimerizes upon FGF binding, and hence attention has focused on activation of the FGFR pathway by receptor homodimerization and heparin-based oligomerization.^47^ The multivalent binders stimulate FGFR activation by dimerizing the FGFR or driving higher order assemblies. We observe considerable differences in pathway activation by the C2, C4, C6 and C8 FGFR engaging ligands, as well as dependence on the geometry of presentation. The C4 extension series revealed a strong distance dependence for activation: mb7 templated 53 Å apart showed strong pERK signaling, whereas the larger constructs with extension lengths of 76 Å and 96 Å did not signal, consistent with the FGF2-FGFR1 dimer complex structure (PDB ID: 1FQ9),^48^ in which the membrane proximal termini are 48 Å apart. Direct measurement of FGFR diffusion in the membrane (**Supplementary Figure 26**) and of the oligomerization state of the receptor in the membrane (**Figure 4D**,**E**), suggest that synthetic ligands drive FGFR clustering.

Currently available naturally occurring signaling molecules (such as FGF2) have pleiotropic effects and it can be difficult to use these to promote differentiation of highly specific cell subpopulations; small molecule treatments can have similar limitations. While our designed agonists broadly phenocopy FGF, there are a number of intriguing differences both in proximal signaling and in the promotion of vascular differentiation. Likely because of the slow offrate of mb7 for FGFR, and the avid binding of the multivalent constructs, the calcium transients have much longer duration for our synthetic agonists than for FGF. The specificity of mb7 for the c-isoform^19^ enables specific activation of signaling through the c-isoform receptor, while addition of mb7 to FGF enables activation of signaling exclusively through the b-isoform. These and perhaps other subtle differences in proximal signaling result in distinct outcomes at multiple developmental stages in vascular differentiation. Our designed scaffolds provide a means to control the prevalence of endothelial cells or pericytes by taking advantage of the capability to activate signaling through just the b or just the c receptor isoforms (**Figure 5F**). In subsequent endothelial cell differentiation, C6-79 promotes the arterial cell fate while FGF2 promotes the venous cell fate, over lymphatic fate.

Our designed proteins have the ability to promote either endothelial cell or pericyte fate, and can even specify subtypes of vascular endothelial cells, which should facilitate studies of blood vessel development from both theoretical and engineering standpoints. The designed oligomers described here provide a general means to drive receptor clustering and sculpt pathway activation for any signaling pathway of interest. Our approach enables multiple levels of control compared to the native signaling molecules: the binding domain can have higher receptor subtype specificity, the on and off rates for receptor subunits can be tuned, and the valency and geometry of receptor engagement can be systematically varied. We envision that such customized synthetic agonists will have broad applications in both ex vivo and in vivo control of cellular differentiation.

## Acknowledgments

We thank Michelle DeWitt for help with protein expression, George Ueda and Brent Herdlicka for advice on endotoxin removal methods, and Ryan Kibler for assistance for SAXS sample submission. We also want to thank Xinting Li, Mila Lamb and Paul Levine for mass spectrometry verification of asymmetric units of the designs. We want to thank Mohamad Abedi and Maggie Ahlrichs for advice on phosphoflow cytometry and cell culture. We thank Luki Goldschmidt and Patrick Vecchiato for computational support. We also want to thank Will Sheffler for computational support. We thank Nanditaa Balachander and Yen Lim for help in molecular biology, Dr. Yan Ting Zhao for help with HUVEC cells, Professors Benjamin Freedman, Ying Zheng and Julie Mathieu for advice in differentiation assays and Dale Hailey and Garvey microscopy core for help with the microscopy. We thank William Rice, Alice Paquette, and Bing Wang of the NYU Cryo-EM Core Facility for assistance with cryoEM grid screening. Collection of 300 kV data was enabled by a block allocation grant through the National Center for CryoEM Access and Training (NCCAT), and we thank Ed Eng, Carolina Hernandez, and Charlie Dubbeldam for their work in data collection, scheduling, sample handling, and grant administration. We thank all members of the Bhabha/Ekiert lab, especially Nicolas Coudray, for helpful discussions regarding computing and data processing, and the NYU high-performance computing (HPC) team. All Krios datasets were collected through NCCAT, part of the Simons Electron Microscopy Center located at the New York Structural Biology Center.

## Author Contributions

N.E. and D.B. conceived the project. A.P and H.R-B conceived analyzing the designed proteins in endothelial differentiation. N.E. and A.E. designed oligomeric constructs. W.Y., B.C., D.R.H. designed building blocks for oligomers. N.E., T.S., A.F., Z.L., L.C., M.M. characterized oligomers. F.P. developed script for oligomer extension. R.R., G.B., D.E., N.E. performed electron microscopy analysis of the designed constructs. N.B. performed cyclic docking of the oligomers. A.P., T.S., S.A., M.G., D.E. performed cell assays. L.C. and B.C. developed minibinder, P.H., A.M., S.G. purified endotoxin free protein, S.L. provided L6-R1c and L6-R1b cells, A.P., S.S. performed transcriptomics study. D.B., L.S., J.S., C.T., B.N., J.S. and H.R-B. supervised the study. N.E., T.S., A.P., R.R., H.R-B and D.B. wrote the manuscript with input from all authors.

## Data and materials availability

Structures are available under the following link: https://files.ipd.uw.edu/pub/2022_Edman_cyclic_oligomers/Edman_et_al_Structures.zip

Sequencing Raw data is available under following link: https://files.ipd.uw.edu/pub/2022_Edman_cyclic_oligomers/sequencing_data.zip

All other scripts and data is available upon request.

