## Supplementary_Information for "Modulation of FGF pathway signaling and vascular differentiation using designed oligomeric assemblies"

#### STAR Methods

##### Key resources table

| REAGENT or RESOURCE | SOURCE | IDENTIFIER |
|---|---|---|
| **Antibodies** | | |
| Monoclonal Rabbit anti-phospho-p44/42 MAPK (Erk 1/2) (Thr202/Tyr204) | Cell Signaling | Cat# 4370S; RRID: AB_2315112 |
| Polyclonal Rabbit anti-p44/42 MAPK (Erk 1/2) | Cell Signaling | Cat# 9102; RRID: AB_330744 |
| Polyclonal Rabbit anti-phospho-FGF Receptor (Tyr653/654) | Cell Signaling | Cat# 3471; RRID: AB_331072 |
| Monoclonal Rabbit anti-FGF Receptor 1 (D8E4) | Cell Signaling | Cat# 9740; RRID: AB_11178519 |
| Monoclonal Rabbit Anti-S6 Ribosomal Protein | Cell Signaling | Cat# 2217S; RRID: AB_331355 |
| Monoclonal Mouse anti-phospho-p44/42 MAPK (ERK 1/2) (pT202/Y204) - AlexaFluor488 | BD Biosciences | Cat# 612592 |
| Goat Anti-Rabbit IgG (H+L) - HRP Conjugate | Bio-Rad | Cat# 1706515 |
| Monoclonal Mouse anti-CD31 (PECAM-1) | Cell Signaling | Cat# 3528, RRID: AB_2160882 |
| Monoclonal Rabbit anti-PDGFRb (28E1) | Cell Signaling | Cat# 3169, RRID: AB_2162497 |
| Goat anti-Mouse IgG secondary antibody, Alexa Fluor 633 | Invitrogen | Cat# A-21050 |

| | | |
|---|---|---|
| Goat anti-Rabbit IgG secondary antibody, Alexa Fluor 488 | Invitrogen | Cat# A-11008 |
| **Bacterial and virus strains** | | |
| E.coli BL21 (DE3) | NEB | Cat# C2527H |
| **Chemicals, peptides and recombinant proteins** | | |
| Fetal Bovine Serum (FBS) | Biowest | Cat# S1620 |
| Penicillin-Streptomycin | Gibco | Cat# 1514012 |
| GlutaMAX Supplement | Gibco | Cat# 35050061 |
| Sodium Pyruvate (100mM) | Gibco | Cat# 11360070 |
| HEPES (1M) | Gibco | Cat# 15630130 |
| Heparin | Fisher BioReagents | Cat# 9041-08-1 |
| Amphotericin B | Gibco | Cat# 15290018 |
| Gelatin from porcine skin | Sigma-Aldrich | Cat# G1890-100 |
| Streptomycin sulfate | Sigma-Aldrich | Cat# S9137 |
| Puromycin dihydrochloride | Gibco | Cat# A11138-03 |
| Human FGF-basic (FGF-2/bFGF) | Gibco | Cat# 13256-029 |
| Tris-HCl | Sigma-Aldrich | Cat# 1185-53-1 |
| Glycerol | Sigma-Aldrich | Cat# G5516 |
| Triton X-100 | Sigma-Aldrich | Cat# 9002-93-1 |
| Sodium dodecyl sulfate (SDS) | Sigma-Aldrich | Cat# 151-21-3 |
| B-Glycerol phosphate | Sigma-Aldrich | Cat# 50020-100G |
| Sodium Fluoride (NaF) | Sigma-Aldrich | Cat# 7681-49-4 |
| Sodium Pyrophosphate | Sigma-Aldrich | Cat# 13472-36-1 |
| Sodium Orthovanadate | Sigma-Aldrich | Cat# 13721-39-6 |
| Phenylmethylsulfonyl fluoride (PMSF) | Roche Life Sciences | Cat# 329-98-6 |
| Benzonase Nuclease | EMD | Cat# 70664 |
| Protease Inhibitor Tablets | Thermo Fisher Scientific | Cat# A32963 |

| | | |
|---|---|---|
| Phosphatase inhibitor cocktail | Sigma-Aldrich | Cat# P5726 |
| 2-Mercaptoethanol | Sigma-Aldrich | Cat# M7522-100 |
| Calbryte 520 AM | AAT Bioquest | Cat# 20651 |
| Heparin Oligosaccharide dp10 | Iduron | Cat# H010 |
| Vectashield with DAPI | Vector laboratories | Cat# H-2000-2 |
| Phalloidin | Invitrogen | Cat# A12380 |
| GSK3-Inhibitor (CHIR99021) | Cayman Chemical | Cat# 13122 |
| Growth Factor Reduced Matrigel | Corning | Cat# 356231 |
| Activin A | Peprotech | Cat# 120-14P |
| Recombinant Human BMP-4 | R&D Systems | Cat# 314-BP-010 |
| B27 supplement, minus insulin | Thermo Fisher Scientific | Cat# A1895601 |
| Recombinant Human VEGF | R&D Systems | Cat# 293-VE-050 |
| L-Ascorbic Acid | Sigma-Aldrich | Cat# A8960 |
| 1-Thioglycerol | Sigma-Aldrich | Cat# M6145 |
| **Critical commercial assays** | | |
| BD Cytofix Buffer | BD Biosciences | Cat# 554655 |
| BD Phosflow PERM buffer | BD Biosciences | Cat# 558050 |
| BD BSA stain Buffer | BD Biosciences | Cat# 554657 |
| Illumina P3 100 cycle kit | Illumina | Cat#20040559 |
| Illumina P2 100 cycle kit | Illumina | Cat#20046811 |
| **Deposited data** | | |
| Cryo-EM Map of C4-131 | This paper | EMD-28958 |
| Cryo-EM Map of C4-71 | This paper | EMD-28974 |
| Cryo-EM Map of C4-71_6x | This paper | EMD-28966 |
| Cryo-EM Map of C4-71_8x | This paper | EMD-28967 |

| | | |
|---|---|---|
| Cryo-EM Map of C4-81 | This paper | EMD-28973 |
| Cryo-EM Map of C6-71 | This paper | EMD-28968 |
| Cryo-EM Map of C6-71_6x | This paper | EMD-28969 |
| Cryo-EM Map of C6-71_8x | This paper | EMD-28970 |
| Cryo-EM Map of C6-79 | This paper | EMD-28889 |
| Cryo-EM Map of C8-71 | This paper | EMD-28888 |
| Cryo-EM Map of C8-71_6x | This paper | EMD-28971 |
| Cryo-EM Map of C8-71_8x | This paper | EMD-28972 |
| Atomic Model of C6-79 | This paper | PDB 8F6R |
| Atomic Model of C8-71 | This paper | PDB 8F6Q |
| Sequencing Data | This paper | GEO: |
| **Experimental models: Cell lines** | | |
| WTC-11 human induced pluripotent stem cells | Coriell | Cat# GM25256 |
| HUVEC (pooled donor) endothelial cells | Lonza | Cat# CC-2519 |
| Chinese Hamster Ovary cells (CHO) (pgsD-677 cells) stably expressing FGFR1c (CHO-R1c) | ATCC<br>Modified cells were gift from Schlessinger Lab | Based on Cat# CRL-2244 |
| Rat Myoblast (L6) stably expressing FGFR1c (L6-R1c) | ATCC<br>Modified cells were gift from Sangwon Lee Lab[19] | Based on Cat# CRL-1458 |
| Rat Myoblast (L6) stably expressing FGFR1b (L6-R1b) | ATCC<br>Modified cells were gift from Sangwon Lee Lab[19] | Based on Cat# CRL-1458 |
| **Oligonucleotides** | | |
| q-RT-PCR Primer Sequences | Integrated DNA Technologies | Supplementary Table VIII |
| **Recombinant DNA** | | |
| pET-29b(+) | Integrated DNA Technologies | |
| **Software and algorithms** | | |

| | | |
|---|---|---|
| Rosetta | Koehler Leman et al.[49] | https://www.rosettacommons.org/software |
| Cyclic Docking Protocol | Fallas et al.[17] | |
| Relion 3.1 | Zivanov et al.[50] | https://relion.readthedocs.io/en/release-3.1/ |
| cryoSPARC v3 | Punjani et al. [51] | https://cryosparc.com/ |
| EMAN2 | Bell et al.[42] | https://blake.bcm.edu/emanwiki/EMAN2 |
| Sparx | Hohn et al.[52] | |
| FrameSlice | Dyer et al.[23] | |
| ScÅtter | Dyer et al.[23] | |
| FOXS server | Schneidman-Duhovny et al.[53] | https://modbase.compbio.ucsf.edu/foxs/ |
| Leginon | Suloway et al.[54] | www.leginon.org |
| MotionCor2 | Zheng et al.[55] | |
| Appion | Lander et al.[56] | www.appion.org |
| 3DFSC server | Tan et al.[57] | https://3dfsc.salk.edu/ |
| UCSF Chimera | Pettersen et al.[58] | https://www.cgl.ucsf.edu/chimera/ |
| Phenix v1.16 | Adams et al.[59] | www.phenix-online.org |
| COOT | Emsley et al.[60] | |
| MATLAB | Mathworks | https://www.mathworks.com/ |
| GaussStorm | Holden et al.[61] | |
| R Studio | Posit | www.posit.co |
| Origin Pro 9.1 | OriginLab | www.originlab.com |
| GraphPad Prism 9 | Dotmatics | www.graphpad.com |
| Fiji | Schindelin et al.[62] | www.imagej.net |
| Monocle3 | Trapnell et al.[63] | https://cole-trapnell-lab.github.io/monocle3/ |
| Adobe Illustrator | Adobe | https://www.adobe.com/prod |

| | | |
|---|---|---|
| | | ucts/illustrator.html |
| CellProfiler | Carpenter et. al.[69] | www.cellprofiler.org |
| **Other** | | |
| Formvar/carbon 400 mesh copper grids | Ted Pella | Cat# 01754-F |

**Scaffold selection and cyclic docking**

Subunit scaffolds consisted of a set of 18 monomeric designed repeat proteins with high-resolution crystal structures or SAXS data[9,64]. PDB IDs for designs with crystal structures are provided in Supplementary Table 1. Docking was performed as previously described.[17] Briefly, the protocol aligns subunits along the desired symmetry axis and scores these using a residue pair-motif database derived from PDB structures. This resulted in 829 outputs among all 5 symmetries attempted (C4, C5, C6, C7, and C8). Outputs were then sequence designed by Rosetta FastDesign to generate an oligomeric interface. These design outputs were filtered by $\Delta\Delta G$ (between -35 and -70), solvent accessible surface area (SASA > 700 $A^2$), shape complementarity (sc > 0.65), and fewer than 2 unsatisfied hydrogen bonds. This resulted in 150 outputs, which were then visually screened for geometric redundancy. Docking for alternative symmetries was performed as above without the interface design step.

**JHR generation**

Junior helical repeat protein (JHR) scaffolds are small curved repeat proteins. The backbones were designed by helical extension based on a library of short helical and loop fragments clustered to include only the most common and ideal fragments. These fragments were pieced together using a published helical extension method to create helix-turn-helix-turn modules that were repeated 4 times to generate 4 repeat DHR-like proteins (manuscript in preparation).[29]

**Repeat Extension Script**

DHR-based oligomers were extended using a custom PyRosetta[65] script that uses an align-and-replace approach. To extend an oligomer with accessible N-termini by two repeats, the second repeat of the 4-repeat parent DHR was aligned to the N-terminal repeat of the oligomer. Subsequently, the terminal repeat of the oligomer was replaced by repeats 2 to 4 of the parent DHR. For oligomers with accessible C-termini the third repeat of the 4-repeat parent DHR was aligned to the C-terminal repeat of the oligomer, and the C-terminal repeat of the oligomer was replaced by repeats 1 to 3 of the parent DHR. The process was repeated to achieve additional extensions.

**Expression and purification**

Sequences of the designed proteins were reverse translated with optimization for *Escherichia coli* expression, with a C-terminal glycine-serine linker followed by a 6x histidine tag. Sequences were ordered as synthetic genes from Integrated DNA Technologies within the pET29b+ vector between NdeI and XhoI cloning sites. This vector contains a kanamycin resistance marker and a T7 promoter. Plasmids were transformed into *E. coli* BL21 (DL3) competent cells and plated on LB with kanamycin at 50mg/L. Transformants were inoculated into 50mL of autoinduction expression media (for 1L: 12g tryptone, 24g yeast extract, 20 mL 50×M, 20 mL 50x5052, 2 mL 1M $MgSO_4$, 200 µL Studier Trace metals, 100 µg kanamycin, q.s. to 1L with filtered water) in a 250mL flask. Expression cultures were grown for 20 hours at 37˚C with 200rpm shaking. Cells were pelleted by centrifugation at 4000xg and resuspended in a lysis buffer consisting of 25mM Tris pH 8, 300mM NaCl, and 20mM imidazole with added protease inhibitor and DNase. Cells were lysed by sonication at 85% amplitude with 8 x 15 second pulses. Lysate was separated into soluble and insoluble fractions by centrifugation at 18,000xg. Immobilized metal affinity chromatography (IMAC) was used to purify designed protein. Nickel-nitrilotriacetic acid (Ni-NTA) resin was initially equilibrated with 5 column volumes (CV) lysis buffer. Supernatant was poured over the columns, followed by 20CV wash buffer (25mM Tris pH 8, 400mM NaCl, 30mM imidazole). Protein was eluted using 5CV elution buffer (25mM Tris pH 8, 300mM NaCl, 500mM imidazole). Eluate was purified by size exclusion chromatography (SEC) on an AKTA PURE FPLC system, using either a Superose 6 Increase 10/300 GL column or a Superdex 200 Increase 10/300 GL column, with Tris-buffered saline (TBS; 25mM Tris pH 8, 150mM NaCl) at a speed of 0.75mL/min. Fractions corresponding to the peak trace were collected and combined for further analysis.

**Low-endotoxin protein production**

Genes were expressed as described above. Cultures were resuspended and lysed in a phospho-buffered saline (PBS)-based lysis buffer with added protease inhibitor and DNase. Cells were sonicated and pelleted as described above. Supernatant was filtered through a 0.45µm filter prior to loading onto IMAC columns. IMAC columns were pre-washed with PBS + 1% Triton X-100 + 0.75% CHAPS to remove any residual endotoxin and equilibrated with PBS + 5 mM imidazole. Supernatant was poured onto the column and followed by washing with 5CV PBS + 30 mM imidazole. To remove endotoxin, 4 wash steps were performed using 5CV PBS + 1% Triton X-100 + 0.75% CHAPS, with 30 min 37 °C incubations on the first and third wash. This was followed by 2 washes with 10CV PBS, then elution with 5CV PBS + 400 mM imidazole. SEC was performed as described above on a dedicated AKTA PURE FPLC with lines, loops, and fraction dispenser pre-washed using 500mM NaOH + 0.75% CHAPS. Endotoxin levels were measured with the LAL endotoxin testing system (Charles River Laboratories).

Proteins expressed by the General Protein Production core were transformed as above then a pre-culture was inoculated into 50mL of LB media and grown at 37°C for 18 hours. 10mL of this pre-culture was used to inoculate 500mL of autoinduction expression media (recipe above) in a 2L flask. Cells were lysed using a Microfluidics M-110P microfluidizer. Soluble and insoluble portions of the lysate were separated at 17000g. Supernatant was flown over 3 mL of nickel resin and washed with PBS wash buffer (20 mM $NaPO_4$, 300 mM NaCl, 30 mM Imidazole, 0.75% CHAPS) for 6 washes of 10mL each (a total of 60 mL). SEC was performed as described above. Endotoxin levels were measured as above.

**Size exclusion chromatography with multi-angle light scattering**

Samples were run in TBS (50mM Tris-HCL, 150mM NaCl pH 8.0) at 1mL/min over a Superose 6 10/300 GL column using an Agilent 1260 HPLC. The HPLC is in line with a Heleos multi-angle static light scattering and Optilab T-rEX detector (Wyatt Technology Co.). Using ASTRA (Wyatt Technology Co.), a weighted average molecular weight (Mw) and number average molar mass (Mn) were calculated to determine monodispersity-by-polydispersity index (PDI), with PDI = Mw/Mn.

**Negative stain EM grid preparation, data collection, and data processing**

Proteins were diluted to 20 μg/ml in TBS, then immediately applied to freshly glow-discharged Formvar/carbon 400 mesh copper grids (Ted Pella catalog #01754-F). After incubation for 45s, excess protein solution was removed by blotting from the side with filter paper, then grids were inverted onto two successive drops of sample buffer followed by three to five successive drops of 2% uranyl formate, with excess solution removed by blotting after each application. The final stain applied was incubated for 15s before blotting. Air-dried grids were imaged using a FEI Talos L120C TEM equipped with a 4K × 4K Gatan OneView camera, at a nominal magnification of 73,000x and pixel size of 2.0 Å. Micrographs were imported to Relion 3.1[50] and/or cryoSPARC v2[51] and, after picking using automated protocols in each program, particles were subjected to 2D classification. Design model projections were generated using EMAN2[66] and Relion, and projections were aligned with experimental 2D class averages using Sparx[52].

**Small-angle X-ray scattering**

SEC-purified samples were prepared for small-angle X-ray scattering (SAXS) by concentrating (if needed) with a 10K molecular weight cut-off spin concentrator followed by filtration with a 0.22μm spin filter. Samples were sent in low (1mg/mL) and high (3-5mg/mL) concentrations in TBS, with flow-through from concentrators used as blanks for later buffer subtraction during data analysis. Scattering data were collected at the SIBYLS High Throughput SAXS Advanced Light Source in Berkeley, California[24] and analyzed with FrameSlice and ScÅtter software packages[23]. Experimental data were compared to design model predictions using the FOXS server (https://modbase.compbio.ucsf.edu/foxs/)[53]. For samples with clear deviation from the design model and indication of off-target symmetry by SEC-MALS and electron microscopy, data were additionally compared to a theoretical design model of the off-target symmetry (described above).

**Cryo-EM grid preparation and data collection**

All grids were plunge-frozen into liquid ethane using a Vitrobot Mark IV with a chamber maintained at 100% humidity and 22°C. Prior to plunge-freezing, 3.5 μL of each design at 0.1 - 1.0 mg/ml was applied to freshly glow-discharged grids of the following types: QUANTIFOIL® R 1.2/1.3 on Cu 400 mesh grids (C6-79), QUANTIFOIL® R 1.2/1.3 on Cu 400 mesh grids + graphene oxide (C8-71; Electron Microscopy Sciences cat. #GOQ400R1213Cu), and/or QUANTIFOIL® R 2/2 on Cu 300 mesh grids + 2 nm C (C4-71 and extensions, C4-81, C8-71 and extensions, C6-71 and extensions, and C4-131). All grids were first screened at NYU on a Talos Arctica microscope operated at 200 kV with a Gatan K3 camera. Larger datasets were acquired for C4-71, C4-81, C6-79, and C8-71 on a Titan Krios microscope operated at 300 kV with a Gatan K3 camera and BioQuantum energy filter ("Krios 6" operated by NCCAT at the New York Structural Biology Center). In both imaging setups, data acquisition was controlled via Leginon[54] and pre-processing (including motion correction and 2X binning) was performed with MotionCor2 as integrated in Appion[55,56]. Further data collection parameters are shown in Supplementary Table II.

Some designs exhibited preferred orientation that appeared to be correlated with ice thickness: in thicker (>30-40nm) ice, side views of the ring predominated, whereas top views (looking down the symmetry axis) could be seen only in the thinnest ice (15-20 nm, as measured by aperture-limited scattering). In such cases, and where grid quality allowed, data were collected in a range of ice thicknesses to minimize orientation bias.

**Processing of 200 kV cryo-EM screening datasets (C4-71 extensions, C6-71 and extensions, C8-71 extensions)**

Aligned, dose-weighted micrographs and STAR files for particles picked "on the fly" with Warp[67] were imported to cryoSPARC[51] v.3 for CTF estimation[68], particle picking, 2D classification, and 3D classification/refinement. 2D classification of particles imported from Warp was used to identify suitable starting classes for template-based auto-picking. In cases where Warp picking did not yield meaningful templates (or omitted certain particle views), unrepresented particle views were located using manual and/or blob picking and classified in 2D to generate additional templates for auto-picking. Initial maps were generated from 2D-curated particles by *ab initio* reconstruction in C1 followed by iterative rounds of heterogeneous and homogeneous refinement. Global resolution (using independent half-maps from refinement and FSC = 0.143 threshold) was estimated using the 3DFSC server (https://3dfsc.salk.edu/)[57]. Additional processing details for each dataset are shown in Supplementary Table III.

**Processing of 300 kV cryo-EM datasets (C4-71, C4-81, C6-79, C8-71)**

Detailed processing workflows are shown in Supplementary Figures S18 and S19. For all 300 kV datasets, aligned and dose-weighted micrographs were imported to cryoSPARC v.2/v.3 for CTF estimation, particle picking, 2D classification, and initial 3D curation and refinement. Templates for C6-79 auto-picking were generated using cryoSPARC's "blob picker". For C4-81, C4-71, and C8-71, 2D averages generated from cryo-EM pre-screening data were used for initial template-based auto-picking. Template-based auto-picking of the C8-71 ring was dominated by side views; to retain top views during auto-picking and curation, these views were picked separately using a single auto-picking template generated from 2D classification of manually-picked top views. For C4-81 and C8-71, curated particles from template picking were used as a training set for Topaz[69] picking within cryoSPARC. The final sets of curated particles from cryoSPARC were imported to Relion v.3[50] for further 2D/3D classification and 3D refinement, which improved map quality for C4-71 and C4-81. Final 3D refinements were performed with the highest expected symmetry imposed, as well as in C1 and with lower-order symmetries imposed. For all datasets, imposing the highest designed circular symmetry improved map quality without introducing substantial artifacts. Global resolution (using independent half-maps from refinement and FSC = 0.143 threshold) and sphericity were estimated using the 3DFSC server (https://3dfsc.salk.edu/)[57]. The FSC mask automatically tightened during the final round of homogenous refinement in cryoSPARC was used for resolution and sphericity calculations for C6-79 and C8-71. Additional processing details for each dataset are shown in Supplementary Table III.

**C6-79 and C8-71 model building and refinement**

*De novo* designed model coordinates for the C8-71 octamer were first docked into the cryo-EM map as a single rigid body using UCSF Chimera[58]. Initial fitting for C6-79 was performed in Chimera with six copies of the designed monomer manually placed into the map, fit as six individual rigid bodies, and merged into a single set of coordinates for the hexamer. In PHENIX v.1.16[59], docked coordinates were stripped of hydrogens using phenix.pdbtools and refined in real space using iterative rounds of phenix.real_space_refine and manual model adjustment in COOT[60,70]. For C8-71, a single instance of simulated annealing was performed at the beginning of automated refinement in PHENIX; non-crystallographic symmetry, secondary structure, Ramachandran, and rotamer restraints were enabled throughout. Additional density is present in the C8-71 cryo-EM maps near W113, at the interface between subunits of the octamer. In addition to the C8 map used for model refinement, this density is also visible at comparable thresholds in C1 and C4 maps refined from the same particles. As no obvious candidate molecule could be identified for this density, it was left unmodelled.

For C6-79, rigid-body refinement and a single instance of simulated annealing were used in early rounds of automated real-space refinement. Non-crystallographic symmetry, secondary structure, Ramachandran, and rotamer restraints were enabled throughout refinement. Additionally, the final round of phenix.real_space_refine included ADP refinement and reference model restraints (using the starting model as a reference to restrain residues 46-53 to manually-adjusted positions and strictly match rotamers). Additional model statistics are shown in Supplementary Table IV.

**Cell culture**

Human umbilical vein endothelial cells (HUVECs) were obtained from Lonza, Germany (#CC-2519). Cells were grown in EGM2 media (20% fetal bovine serum [BioWest, #S1620], 1% penicillin-streptomycin [Gibco, #1514012], 1% Glutamax [Gibco #35050061], 1% ECGS [endothelial cell growth factor], 1mM sodium pyruvate [Gibco, #11360070], 7.5mM HEPES [Gibco, #15630130], 0.08mg/mL heparin [Fisher BioReagents, #9041-08-1], 0.01% amphotericin B [Gibco, #15290018], a mixture of 1X RPMI 1640 +/- glucose [Gibco, #1187902] for a final concentration of 5.6mM glucose; filtered through 0.2-μm filter) on 0.1% gelatin-coated [Sigma, #G1890-100] 35mm cell culture dishes. Cells were cryopreserved at passage 4 for later thawing and use in Western blots.

ECGS was extracted from 25 mature whole bovine pituitary glands from Pel-Freeze biologicals [Lonza, #57133-2]. Pituitary glands were homogenized with ice-cold 0.15M NaCl [Fisher Chemical, #CAS7647-14-5] and adjusted to pH 4.5 with HCl [Sigma-Aldrich, 320331]. Following 1hr centrifugation at 4°C, the supernatant (wine colored) was collected and adjusted to pH 7.6, followed by addition of 0.5g/100 mL of streptomycin sulfate [Sigma, #S9137]. The following day, the supernatant was centrifuged at 4,000 RPM for 1hr at 4°C. The supernatant was sterile filtered using a 0.45-μm filter and stored at -20°C.

Parental heparan-deficient Chinese hamster ovary (CHO) cells [pgsD-677 cells; ATCC, #CRL-2244] stably expressing human FGFR1c were maintained in F-12K medium (ATCC, #30-2004) supplemented with 10% fetal bovine serum [BioWest, #S1620] (manuscript in preparation), 1% penicillin-streptomycin [Gibco, #1514012], and 10 μg/mL puromycin [Gibco, #A11138-03]. Rat myoblast (L6) cells [ATCC, #CRL-1458] stably expressing either human FGFR1c (L6-R1c) or FGFR1b (L6-R1b)[19] were maintained in DMEM medium [Gibco, #10566] supplemented with 10% fetal bovine serum [BioWest, #S1620], 1% penicillin-streptomycin [Gibco, #1514012], and 10 μg/mL puromycin [Gibco, #A11138-03].

**Treatment and protein isolation for Western blot**

For activation assays, cells were seeded onto 12-well plates and grown to ~80% confluence. Cells were serum-starved overnight in their respective media (F-12K for CHO cells, DMEM low glucose (1 g/L) [Gibco, 11885-084] for HUVEC, L6 cells). The following day, cells were stimulated with different concentrations of either recombinant FGF2 [Gibco, #13256-029] or designed scaffolds at 37 °C for 15 min. Concentration is reported as the concentration of the oligomeric particle, not the mb7 domain; therefore, 10 nM of C4-71C_mb7 corresponds to 10 nM of C4 oligomer and 40 nM of mb7. Following treatment, cells were washed once with 1X PBS before harvesting total protein for analysis.

Cells were lysed with 130µl of lysis buffer containing 20 mM Tris–HCl [Sigma-Aldrich, #1185-53-1] (pH 7.5), 150 mM NaCl, 15% Glycerol [Sigma-Aldrich, #G5516], 1% Triton [Sigma-Aldrich, #9002-93-1], 3% SDS [Sigma-Aldrich, #151-21-3], 25 mM b-Glycerophosphate [Sigma-Aldrich, #50020-100G], 50 mM NaF [Sigma-Aldrich, #7681-49-4], 10 mM Sodium Pyrophosphate [Sigma-Aldrich, #13472-36-1], 0.5% Sodium Orthovanadate [Sigma-Aldrich, #13721-39-6], 1% PMSF [Roche Life Sciences, #329-98-6], 25 U benzonase nuclease [EMD, #70664-10KUN], protease inhibitor cocktail [Pierce Protease Inhibitor Mini Tablets, Thermo Scientific, #A32963], and phosphatase inhibitor cocktail 2 [Sigma-Aldrich, #P5726] in a tube. 43.33µl of 4X Laemmli Sample Buffer [Bio-Rad, #1610747] containing 10% beta-mercaptoethanol [Sigma-Aldrich, #M7522-100] was added to the cell lysate and then heated at 95°C for 10 min. The boiled samples were either used immediately for Western blot analysis or stored at -80°C.

**Western blotting**

If frozen, protein samples were thawed and heated at 95 °C for 10 minutes. A 4-10% SDS-PAGE gel was loaded with 30uL of protein per well and separated for 30 min at 250V. Proteins were transferred onto a nitrocellulose membrane for 12 minutes using the semi-dry turbo transfer Western blot apparatus [Bio-Rad]; the membrane was then blocked in 5% bovine serum albumin for 1 hour. The membrane was incubated with the appropriate primary antibodies on a rocker at 4°C overnight. The antibodies used in this study were pERK1/2 p44/42 [Cell Signaling, #4370S] at 1:1,000 dilution, S6 [Cell Signaling, #2217S] at 1:1,000 dilution, ERK1/2 p44/42 [Cell Signaling, #9102] at 1:1,000 dilution, Phospho-FGF Receptor (Tyr 653/654) [Cell Signaling, #3471] at 1:1,000 dilution, and FGF Receptor 1 (D8E4) [Cell Signaling, #9740] at 1:1,000 dilution. The next day, membranes were washed with 1X TBS-T (3 times, 10min intervals) and incubated with the respective HRP-conjugated secondary antibody (1:10,000 dilution in 5% bovine serum albumin; Bio-Rad) at room temperature for 1 hour. All the membranes were washed with 1X TBS-T (3 times, 10 min intervals) after secondary antibody incubation, developed using Chemiluminescence developer, and imaged using Bio-Rad ChemiDoc Imager.

**Calcium release assay**

CHO-R1c cells were seeded on 96-well flat bottom microplates [Corning, #3603] and grown to ~70-80% confluence. Cells were starved in serum-free F12-K medium for 3 hours. Following starvation, the cells were incubated in serum-free media containing 5µM Calbryte 520 AM fluorescent intracellular calcium indicator [AAT Bioquest, #20651] for 30 min at 37°C. Cells were washed 3X with serum-free media and treated with various concentrations of recombinant FGF2 (with or without 40µg/mL heparin [Iduron, #H010]) or designed scaffolds. Confocal live imaging was done on a Leica TCS-SPE Confocal microscope using a 20X objective and Leica Software. Parameters for each live frame: Excitation/Emission filters for GFP fluorescence, Exposure time of 150ms, Acquisition rate of 5 sec/frame, and total recording time of 15 minutes (5 min baseline recording + 10 min ligand treatment time). Images were processed with Fiji software distribution of ImageJ v1.52i[62,71] and frame-by-frame cellular fluorescence intensity was tracked and quantified with CellProfiler[72,73]. Dose-specific average calcium release was calculated by tracking each individual cell's response during the recording time and computing the mean peak fluorescence achieved by all cells in the frame. An average of 50-100 cells were tracked per recording.

**Phosphoflow Assay**

CHO-R1c cells were grown in T75 flasks. One day before the experiment, cells were changed to starvation medium (F12K+P/S). On the day of the experiment, cells were washed, trypsinized, plated at 200k cells per well in a 96-well plate and incubated with ligands at corresponding concentrations for 15 min in starvation conditions at 37 °C in 100 µl. Afterwards cells were immediately fixed with 100 µl of prewarmed BD Cytofix buffer (BD Biosciences, #554655) and incubated at 37 °C for 10 min. Cells were spun down by centrifugation for 5 min at 300xg and supernatant was discarded by inverting the plate. The plate was gently vortexed, 100 µl of BD Phosflow PERM buffer (BD Biosciences, #558050) was added and cells were incubated for 30 min in the dark on ice. After the incubation, cells were washed twice with 200 µl of BSA stain buffer (BD Bioscience, #554657). After the washing steps, cells were resuspended in 100 µl of 1:10 diluted pERK-AlexaFluor488 (BD Biosciences, #612592) in BSA stain buffer and incubated for 30 min at RT in the dark. Cells were washed twice with 200 µl of BSA stain buffer and after final resuspension in 200 µl of BSA stain buffer immediately analyzed with the Attune flow cytometer. For analysis FSC, SSC and AlexaFluor488 laser settings were set to 1, 250 and 330. The plate autosampler was run at 100 µl/min and cells were gated to a single cell population and geometric mean of the population was calculated and plotted via Origin Pro 9.1. Data were fit using a Hill function in Origin. Data were normalized on 10 nM of FGF (ThermoFisher Scientific, #PHG0369) stimulation.

**Biolayer Interferometry (BLI) Assay**

BLI measurements were performed with the Sartorius Octet system. Streptavidin harboring tips were incubated in Octet Buffer for 30 min before the measurement. For the measurement, tips were equilibrated in Octet Buffer for 150 s, then biotinylated FGFR2 receptor (ectodomain residues 147-366, UniProt ID: P21802, previously expressed in mammalian cells using a IgK signal peptide (METDTLLLWVLLLWVPGSTG) at the N-terminus and a C-terminal TEV cleavage site, 6-His and Avitag (GSENLYFQGSHHHHHHGSGLNDIFEAQKIEWHE)) was loaded onto the tips at 30 nM for 300 s. After a brief equilibration in Octet Buffer for 300 s, tips were dipped into different concentrations of ligands for association for 1400-1800 s. Dissociation was performed for 1400 s in Octet Buffer. Data were analyzed and fit via the Octet Analysis Software.

**TIRF microscopy**

For single-molecule imaging experiments, pgsD-677 cells were plated on 35-mm glass-bottom dishes (MatTek Corporation,#P35G-1.5-14-C) to 75% confluence in phenol-red free DMEM (Gibco, #21063029) supplemented with 4.5 g/L glucose and 10% (vol/vol) FBS (FBS; Gibco, #16140071) and transfected with 0.25 g HaloTag-FGFR1c plasmid the next day using Lipofectamine 3000 reagent (Invitrogen, #L3000001), according to the manufacturer's instructions. The following day, cells were starved for 2-3 hours in serum-free media, labeled with 0.25 M cell-impermeant Alexa488 HaloTag ligand (Promega, #G1001) for 15 min at 37°C and 5% CO2, and then washed 3x with phenol-red free media. After labeling, cells were immediately imaged at 37°C and 5% CO2 in a cage incubator (OkoLab) housing a Nikon Eclipse Ti2 microscope (Nikon) equipped with a motorized Ti-LA-HTIRF module with a 15-mW LU-N4 488 laser, using a CFI Plan Apochromat Lambda 100x/1.45 Oil TIRF objective and a Prime95B cMOS camera (110-nm pixel size; Teledyne Photometrics). Images were acquired using a 100-ms exposure time at 10 Hz with the laser power set at 100%. The penetration depth of the evanescent field was ~118 nm.

**Single-particle tracking**

Particles were localized and tracked using the MATLAB software GaussStorm.[61,74] Briefly, particles were automatically detected by application of a bandpass filter to remove noise, followed by convolution with a Gaussian kernel, and then the selection of above-threshold pixels. Particles were then fitted with elliptical two-dimensional Gaussian functions, which yielded their intensities expressed as the volume under the curve, as well as their positions with subpixel accuracy. Particles were tracked frame to frame using a tracking algorithm with a tracking window of 7 pixels between consecutive frames. The distribution of the displacements of single particles was used to calculate mean diffusion coefficient in a field of view encompassing an entire cell.

**Transcriptomics on HUVEC endothelial cells**

HUVEC endothelial cells were seeded at a density of 80,000 cells/well in a 0.1% gelatin-coated 12-well tissue culture dish, and allowed to grow to 80% confluence. Cells were washed 3X with 1X PBS and serum-starved overnight in DMEM low glucose (1 g/L). Following starvation, cells were treated with either recombinant FGF2 or C6-79C_mb7 at 10 nM, or 100 nM in serum-free media for 6 hours. Concentration is reported as the concentration of the oligomeric particle, not the mb7 domain. After treatment, cells were enzymatically detached using Tryp-LE (Thermo, #12563011), pelleted at 500g for 5 minutes and washed once with cold PBS. Cells from each treatment were then counted and loaded at a concentration of 10,000 cells/lane on the 10x 3' gene expression platform (10x genomics, PN-1000121). After library preparation, libraries were sequenced on the Nextseq 550 with a 75 cycle high-output kit (Read1: 26bp, Index1:8bp, Read2:58). Processed reads were then mapped using the 10x cell-ranger pipeline and mapped to the hg38 reference genome. Transcriptomes from treated samples (recombinant FGF or C6-79C_mb7) were then compared to serum starved cells using the fit_models() function in the Monocle3 software suite. Cells treated with C6-79C_mb7 showed a similar transcription pattern in comparison to cells treated with FGF2 (**Supplementary Figure 35**)[6].

**Immunostaining of differentiated iPSCs**

For immunofluorescence imaging of differentiated iPSCs, cells were seeded on glass coverslips coated with 0.1% gelatin on Day 5, and cultured until confluency on Day 14 following the process described below. The cells were then fixed with 4% paraformaldehyde (PFA) for analysis. The fixed cells were washed three times for 5 min each in 1X PBS before blocking for 1 hr with 3% BSA (VWR, 0332-500G) and 0.1% Triton X-100 (Sigma, T9284-500ML) in 1X PBS while on nutation. Primary antibody incubation was carried out at a 1:100 dilution in blocking buffer overnight: CD31 (Cell Signaling, Catalog #3528), and

PDGFR-B (Cell Signaling, Catalog #3169). Following overnight incubation, the cells were washed three times for 5 min each in 1X PBS while on nutation. The cells were then incubated with secondary antibodies (Invitrogen, A21050 and Invitrogen, A11008; 1:100 each) and Phalloidin (1:100, Invitrogen, A12380) diluted in blocking buffer for 1.5 hrs at 37°C. Secondary antibodies were then removed, and cells were washed three times for 10 min each in 1X PBS on nutation. Coverslips were sealed using VECTASHIELD including DAPI (Vector laboratories, H-2000-2) upside-down on glass slides for analysis in confocal (Leica) microscopy.

***In vitro* differentiation of endothelial cells**

Briefly, hiPSCs (WTC-11 human induced pluripotent stem cells) [Coriell, #GM25256] were seeded on 24-well plates coated with growth factor-reduced Matrigel [Corning, #356231] and cultured in mTeSR1 stem cell medium [StemCell Technologies, #85850] until cells reach confluence with media changes daily. One day before differentiation (deemed Day (-1)), cells were pre-treated with mTeSR1 supplemented with 1μM of GSK3-Inhibitor (CHIR99021) [Cayman Chemicals, #13122]. On the first day of differentiation (D0), stem cell media was replaced with cardiogenic mesoderm media consisting of RPMI 1640 Medium [Thermo, #11875093] supplemented with B27(-) [Fisher Scientific, #A1895601], 100ng/mL Activin A [PeproTech, #120-14P] and Matrigel for 17hrs. The next day, media was replaced with RPMI supplemented with 1μM of GSK3-Inhibitor (CHIR99021), B27 (-), and 5ng/mL bone morphogenetic protein-4 (BMP-4) [R&D systems, #314-BP-010] for 24 hours. On Day 2 of differentiation, cells were washed with 1X PBS and media was replaced with vascular differentiation media consisting of StemPro [Thermo Fisher, #10639011] supplemented with 1X Glutamax, 1X penicillin-streptomycin, 300ng/mL vascular endothelial growth factor (VEGF) [R&D systems, #293-VE-050], 5ng/mL BMP-4, 5ng/mL FGF2, 50ug/mL Ascorbic Acid [Sigma-Aldrich, #A8960], and 40μM monothioglycerol (MTG) [Sigma-Aldrich, #M6145]. On Day 5, cells were dissociated with Accutase [Thermo, #A1110501] and replated on 12-well 0.1% gelatin-coated tissue culture dishes in endothelial growth media (EGM) consisting of EBM basal media [Lonza, #CC-3121] supplemented with 20ng/mL VEGF, 20ng/mL FGF2 and 1μM GSK3-Inhibitor (CHIR99021). EGM media was replaced every 48 hours until the final harvest at Day 28.

After harvest, samples from each day were exposed to an hypotonic lysis buffer (10mM Tris-HCl Ph7.4, 10 mM NaCl, 3 mM MgCl2, 0.05% IGEPAL), labeled with hash oligos (**Supplementary Table IX**), chemically fixed, and then stored at -80°C until cells from all experimental timepoints had been collected. Following collection, cells were processed using the sci-RNA-seq as described previously[75]. Following library preparation, libraries were sequenced on 2 Nextseq2000 100 cycle kit with standard sequencing chemistry: Read1: 34bp, Index1: 10bp, and Read2: 66bp. Reads were then demultiplexed, assigned to cells and mapped to the hg38 reference genome. Sample barcodes were matched to a corresponding experimental condition only if a sample barcode was significantly enriched (Chi-squared test; q-value < 0.05) and displayed a 4 fold enrichment ratio in that cell[75].

All low-quality reads were removed from the data by setting UMI cutoff to greater than 100 and removing all mitochondrial reads. To eliminate effects of cell-cycle heterogeneity, we used Seurat's[76] workflow for cell-cycle scoring and regression. Following Monocle3's workflow, the data were normalized by size factor, preprocessed using PCA, embedded in 2 dimensions with UMAP, and clustered. Top marker analysis was performed to identify genes that were specifically expressed in each cluster, and this information was used to annotate each cluster based on the relative expression of canonical marker genes. The two clusters obtained at day 14 were compared across conditions to determine the relative contribution of each treatment to either the endothelial or pericyte cluster.

The differentiation and single cell sequencing experiment was repeated, collecting only cells on day 14. After processing the data, as described above, the endothelial cell cluster was selected for further sub-clustering, and the analysis was repeated (as described above). This analysis indicated that marker genes specific to arterial, venous and lymphatic endothelial cells, spanned the embedding. Based on marker gene expression, the cells were annotated into arterial, venous and lymphatic endothelial cells, and the localization of FGF2 and C6-79C_mb7 treated cells was calculated to determine the relative contribution of each treatment.

### Supplementary Figures

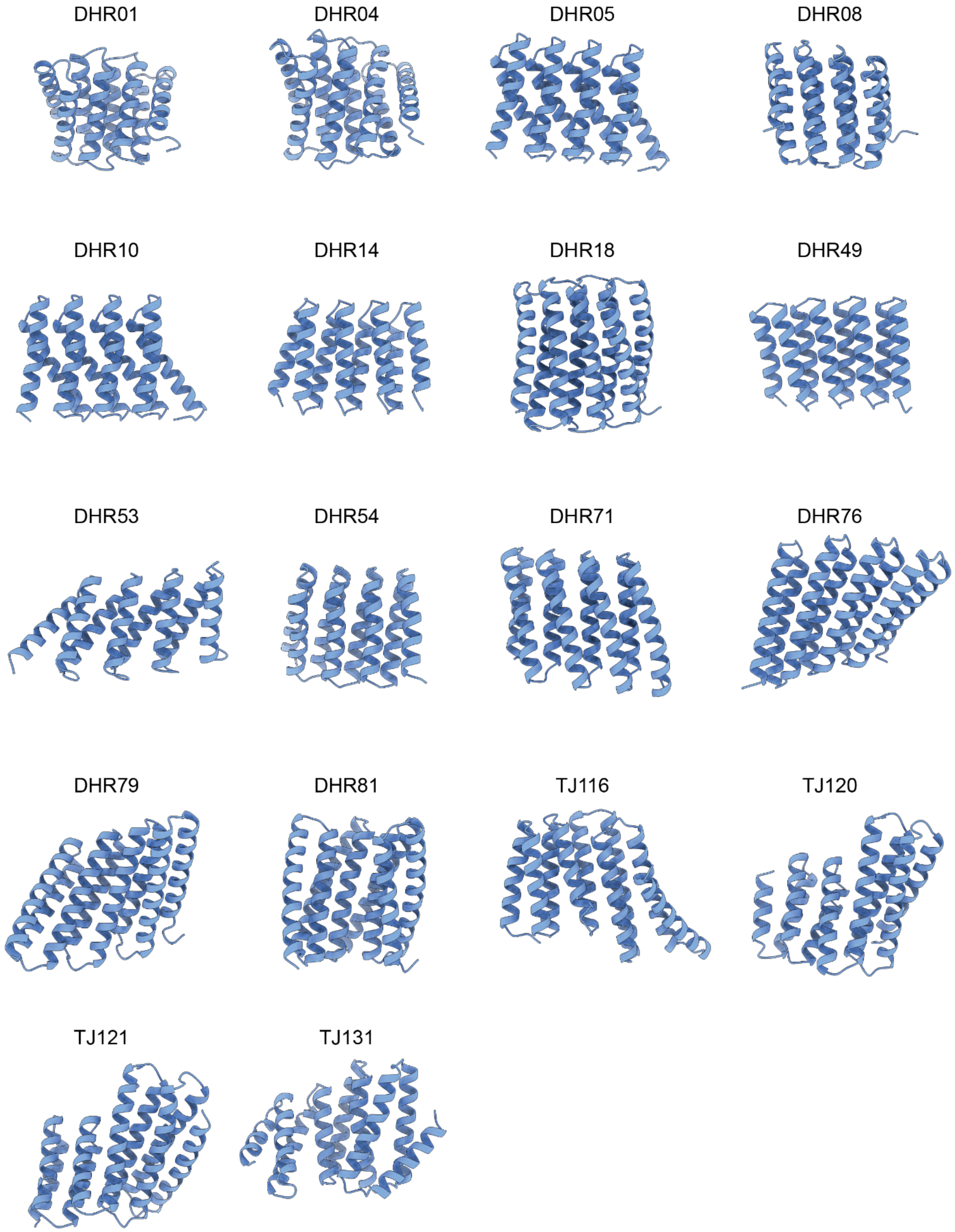


**Figure S1.** Library of building blocks used for docking Oligomers into various symmetries.

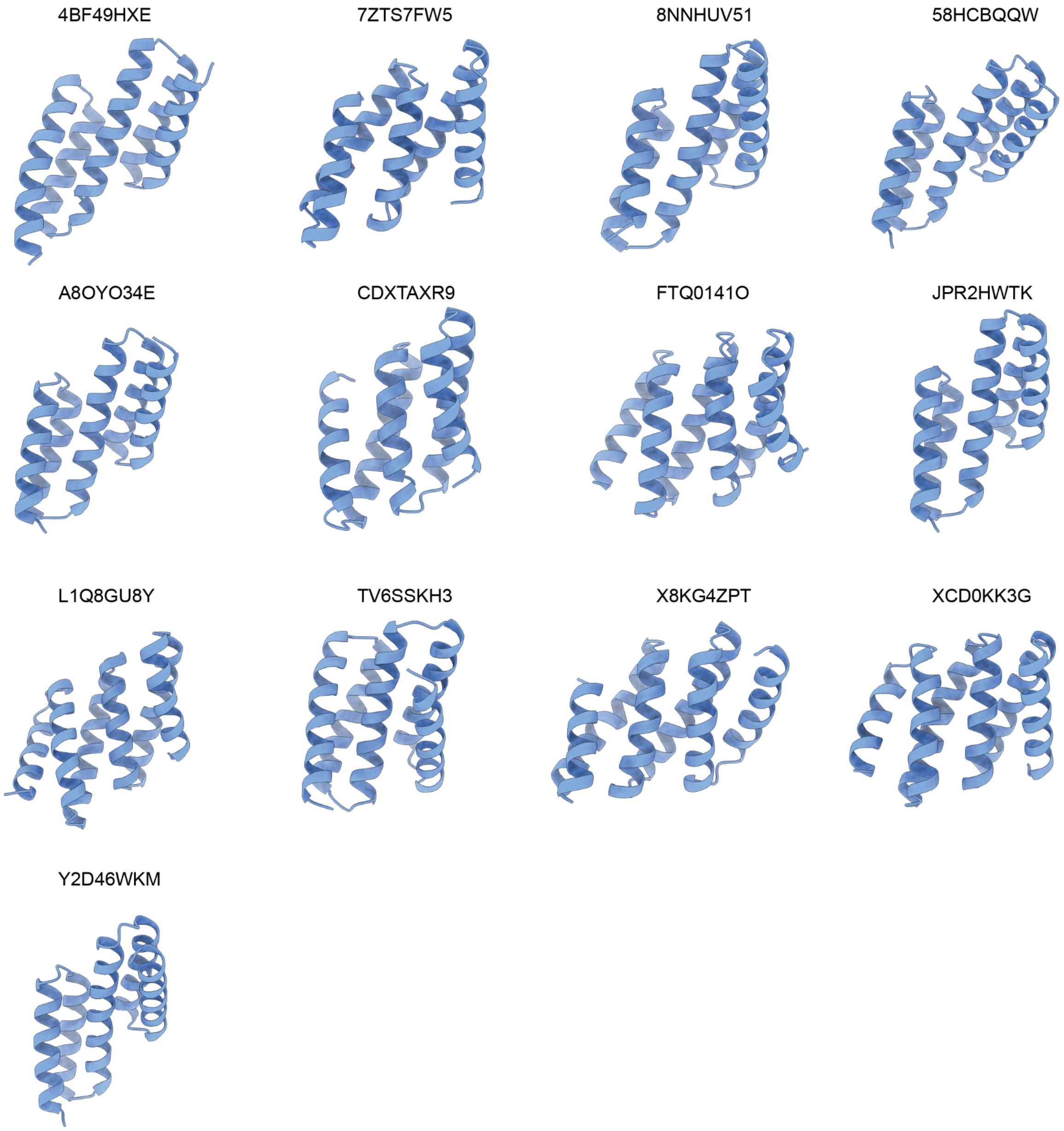


**Figure S2:** Library of building blocks used for docking Oligomers into C2 symmetries that advanced to experimental characterization.

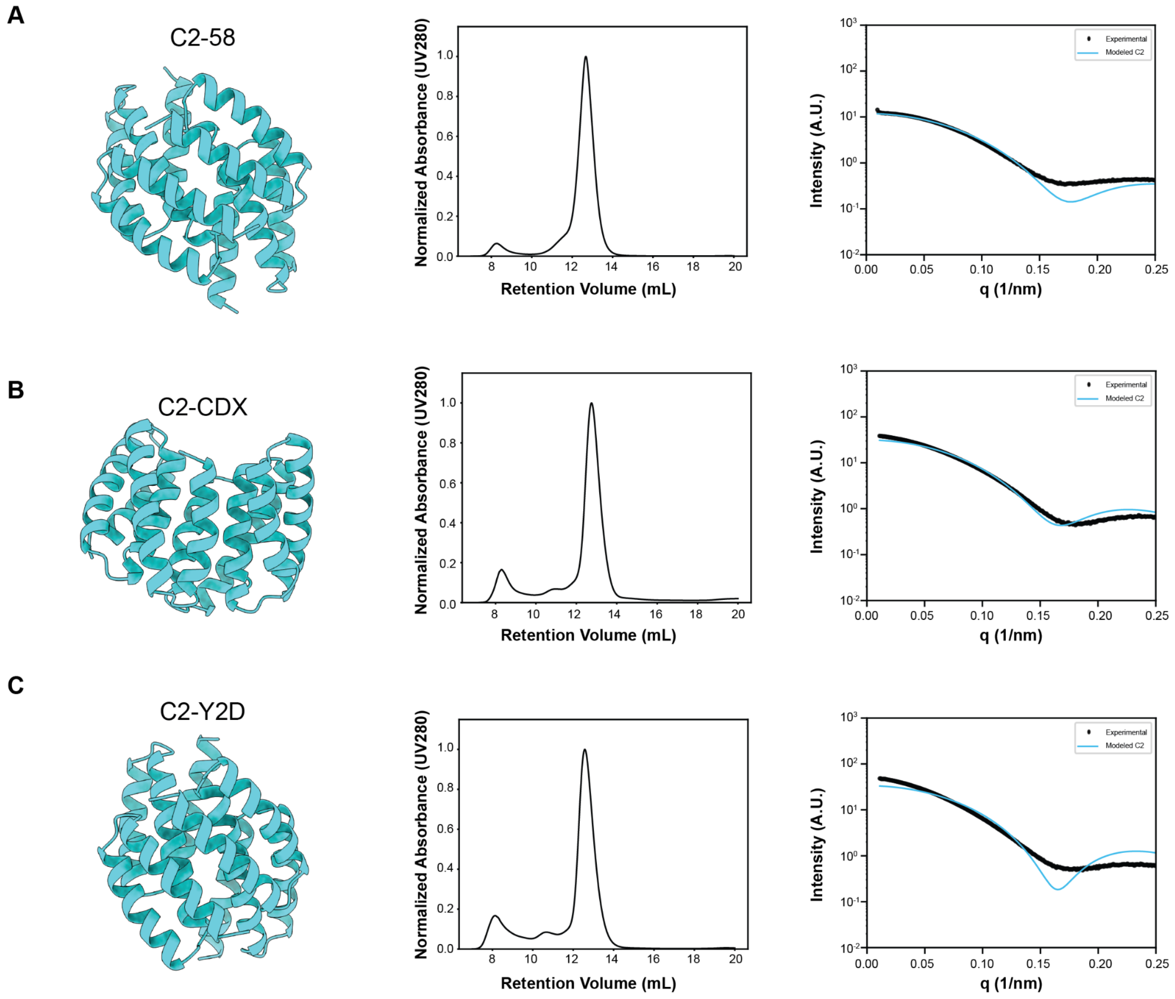


**Figure S3.** Biophysical characterization of C2-Oligomers. *Left*: Design model. *Middle*: SEC characterization. *Right*: SAXS trace comparison of data (black) versus design (cyan). (A) C2-58 (B) C2-CDX (C) C2-Y2D construct.

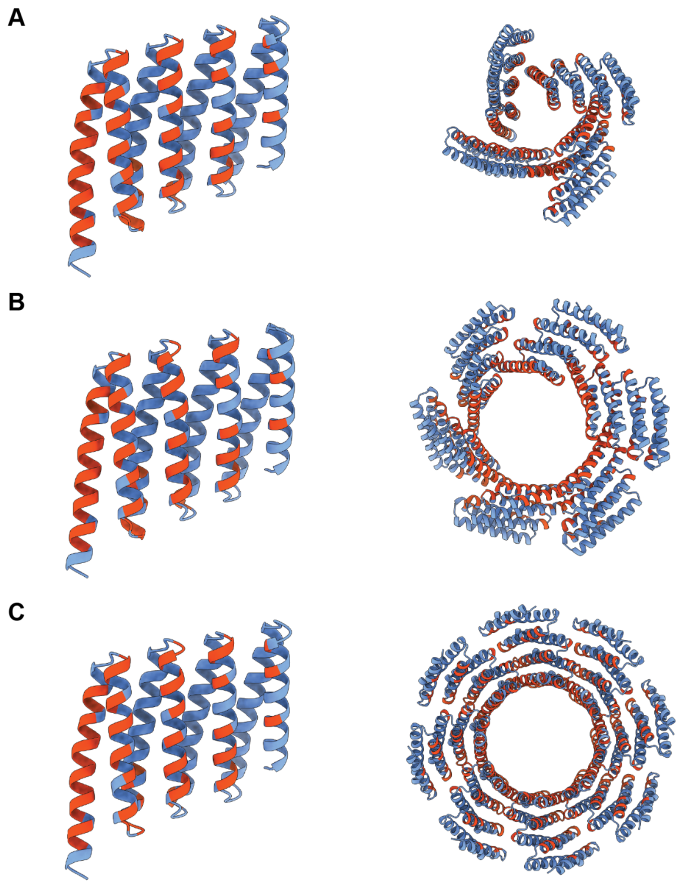

**Figure S4.** DHR-71 based oligomer formation. Residues that changed in comparison to the different oligomeric states are labeled in red. (A) C4-71 asymmetric unit (left) and oligomeric structure (right). (B) C6-71 asymmetric unit (left) and oligomeric structure (right). (C) C8-71 asymmetric unit (left) and oligomeric structure (right).

```
Symbols     ********:******:**.***********
Consensus   PEEILERAKESLERAREASERGDEEEFRKA
C4-71       PEEILERARESLERAREASERGDEEEFRKA  30
C6-71       PEEILERAKESLERAKEAFERGDEEEFRKA  30
C8-71       PEEILERAKESLERAREASERGDEEEFRKA  30

Symbols     ********************* :*. **.*
Consensus   AEKALELAKRLVEQAKKEGDPELVXEAARV
C4-71       AEKALELAKRLVEQAKKEGDPWMVMWAALV  60
C6-71       AEKALELAKRLVEQAKKEGDPELVFEAARV  60
C8-71       AEKALELAKRLVEQAKKEGDPELVLEAARV  60

Symbols     ***** **. .*******************
Consensus   ALWVAXLAAXNGDKEVFKKAAESALEVAKR
C4-71       ALWVALLALRNGDKEVFKKAAESALEVAKR  90
C6-71       ALWVAWLAAWFGDKEVFKKAAESALEVAKR  90
C8-71       ALWVAELAAKNGDKEVFKKAAESALEVAKR  90

Symbols     *****..****::*. ** *** ** :* :
Consensus   LVEVASKEGDPELVLXAAXVALXVAXLAXL
C4-71       LVEVASKEGDPEMVLLAAWVALFVAWLAWL  120
C6-71       LVEVAKEEGDPELVLKAAFVALLVAIMAVI  120
C8-71       LVEVASKEGDPDLVAWAALVALWVAFLAFL  120

Symbols      ******************.*:*:* :.*.
Consensus   XGDKEVFKKAAESALEVAKRLVEVAXKEGD
C4-71       FGDKEVFKKAAESALEVAKRLVEVASKEGD  150
C6-71       LGDKEVFKKAAESALEVAKRLVEIAAREGD  150
C8-71       NGDKEVFKKAAESALEVAKALMEVAMKVGA  150

Symbols     * *** *..**. *  **.. *****..:*
Consensus   PELVEEAAKVAEXVXXLAELXGDEEVREKA
C4-71       PELVEEAAKVAEEVEKLAEKQGDEEVREKA  180
C6-71       PELVEEAAKVAELVRELAKLMGDEEVYEKA  180
C8-71       PWLVELAIAVARAVWLLAELFGDEEVRRRA  180

Symbols      ..  .:.:  : *: . ***
Consensus   XETXXEVRLXXLXVRXWXGGG
C4-71       WETWMEVWLLWLEVRLRKGGG  201
C6-71       RETAREVRLFLLFVRIWEGGG  201
C8-71       EAFEIILRIAAIAVKAWLGGG  201
```

**Figure S5.** Multiple Sequence Alignment of DHR-71 based Oligomers.

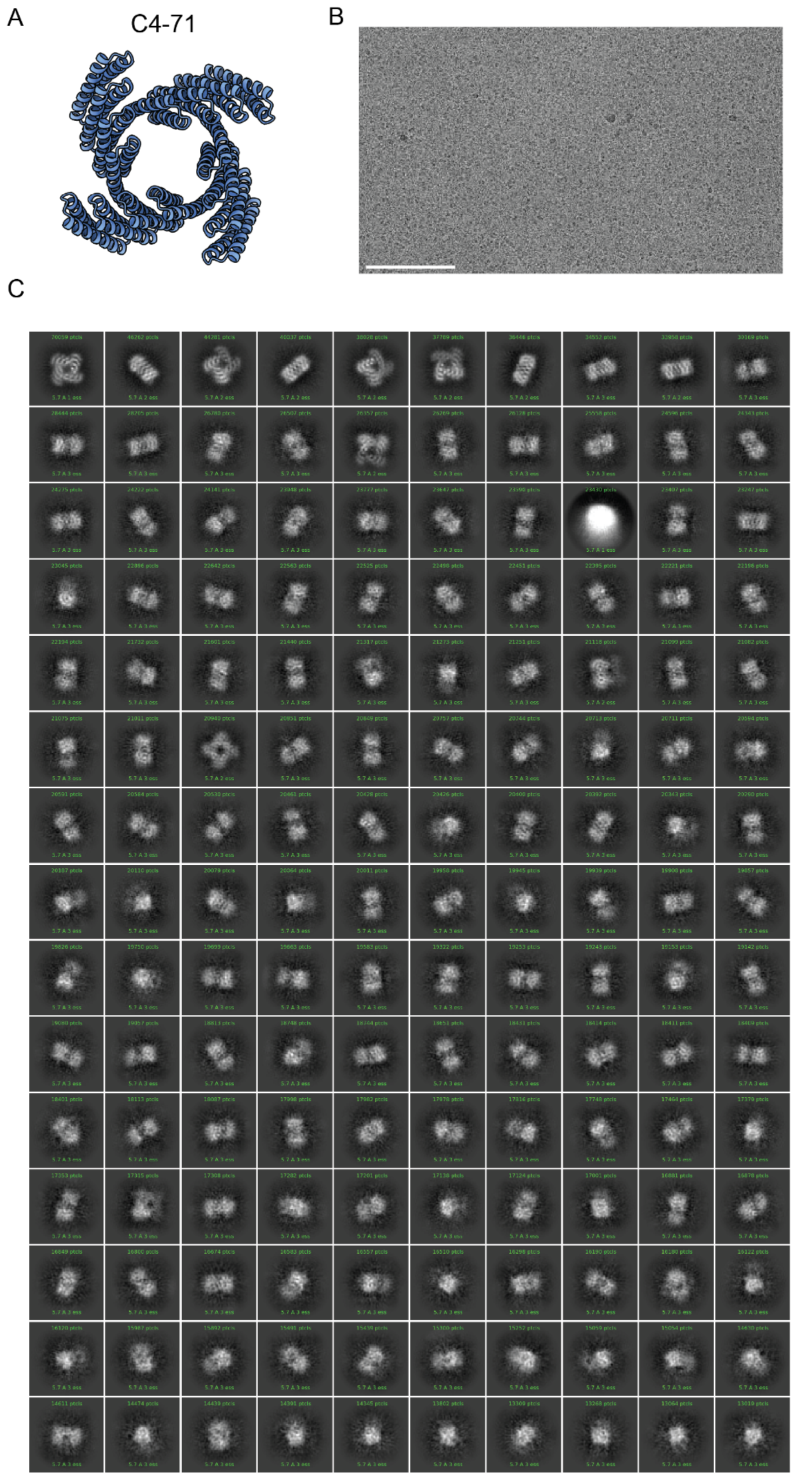


**Figure S6:** (A) C4-71 design model (B) cryo-EM grid image (C) class averages. Scale Bar: 100 nm (B)

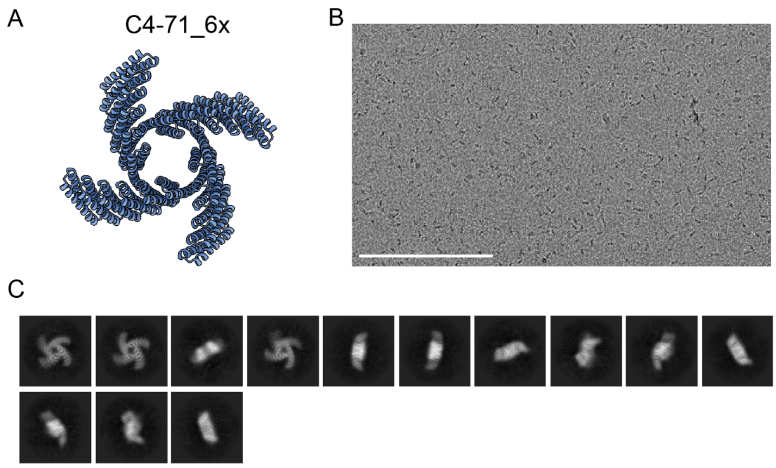


**Figure S7:** (A) C4-71_6x repeat extension design model (B) cryo-EM grid image (C) class averages. Scale Bar: 200 nm (B)

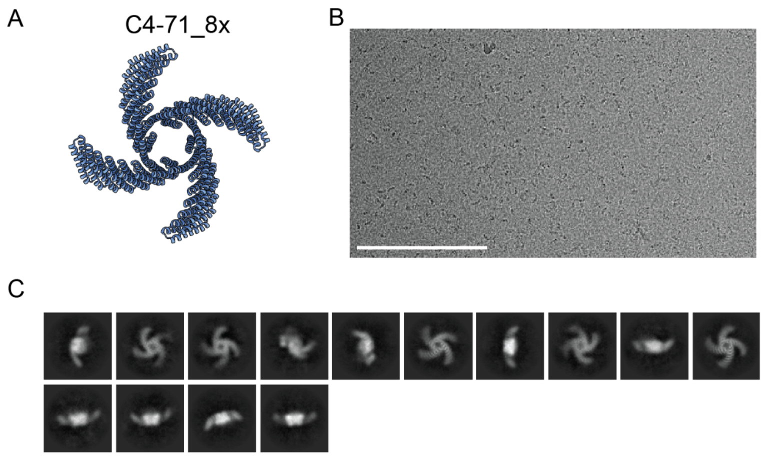


**Figure S8:** (A) C4-71_8x repeat extension design model (B) cryo-EM grid image (C) class averages. Scale Bar: 200 nm (B)

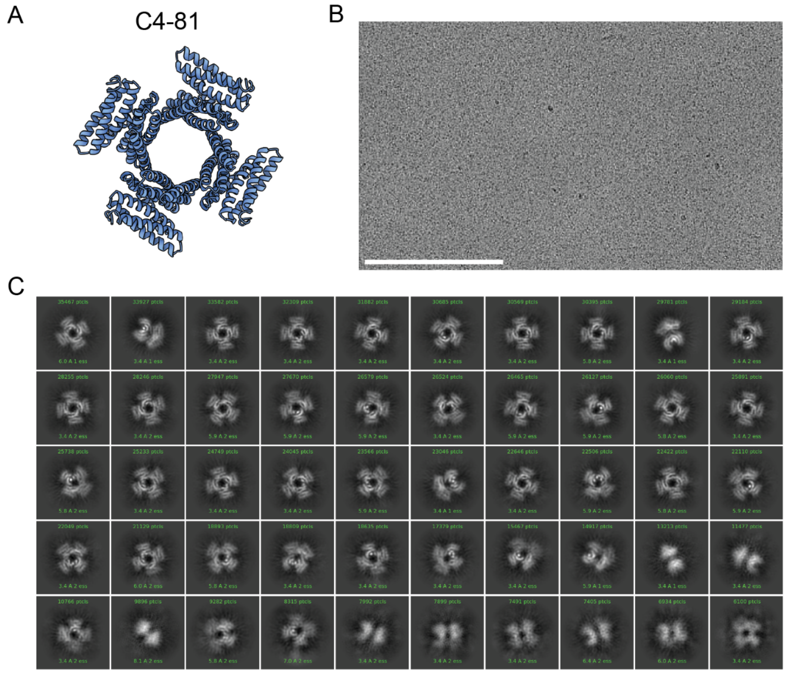


**Figure S9:** (A) C4-81 design model (B) cryo-EM grid image (C) class averages. Scale Bar: 200 nm (B)

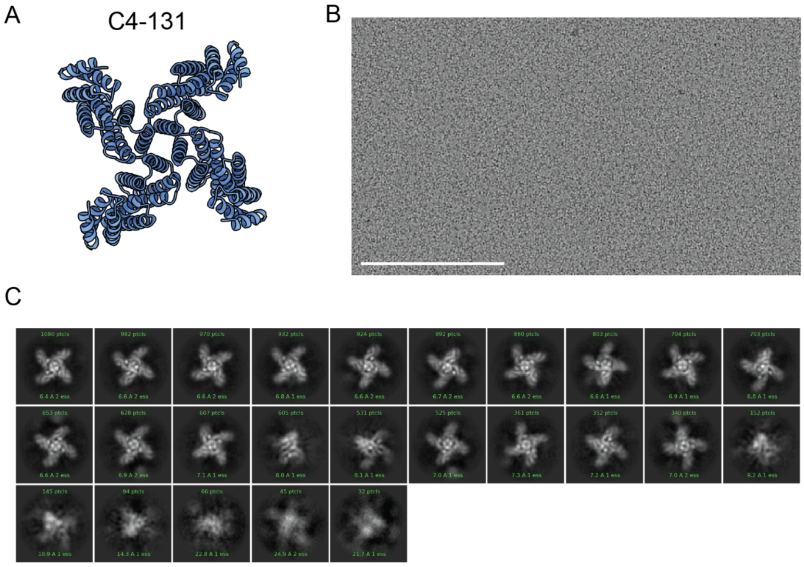


**Figure S10:** (A) C4-131 design model (B) cryo-EM grid image (C) class averages. Scale Bar: 200 nm (B)

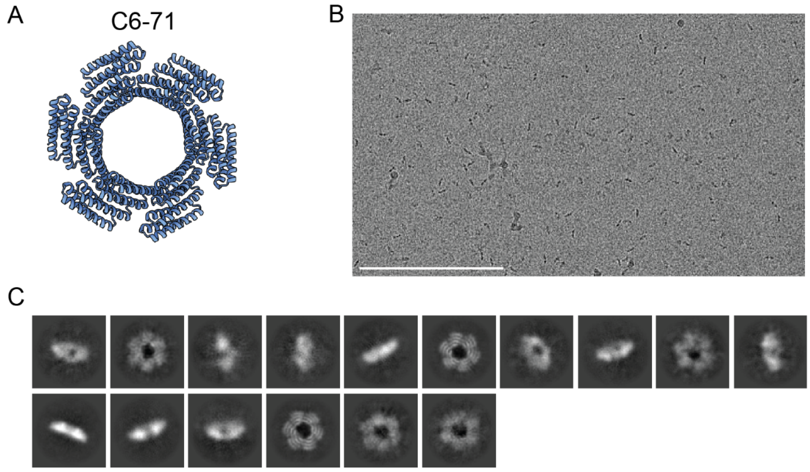


**Figure S11:** (A) C6-71 design model (B) cryo-EM grid image (C) class averages. Scale Bar: 200 nm (B)

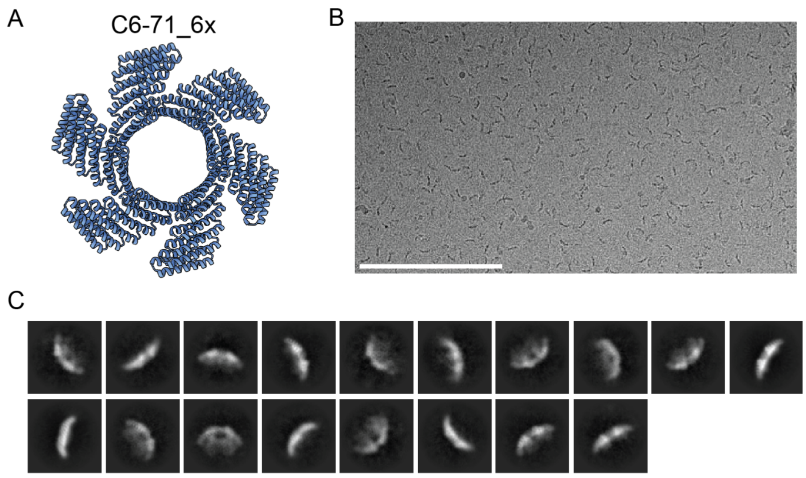


**Figure S12:** (A) C6-71_6x repeat extension design model (B) cryo-EM grid image (C) class averages. Scale Bar: 200 nm (B)

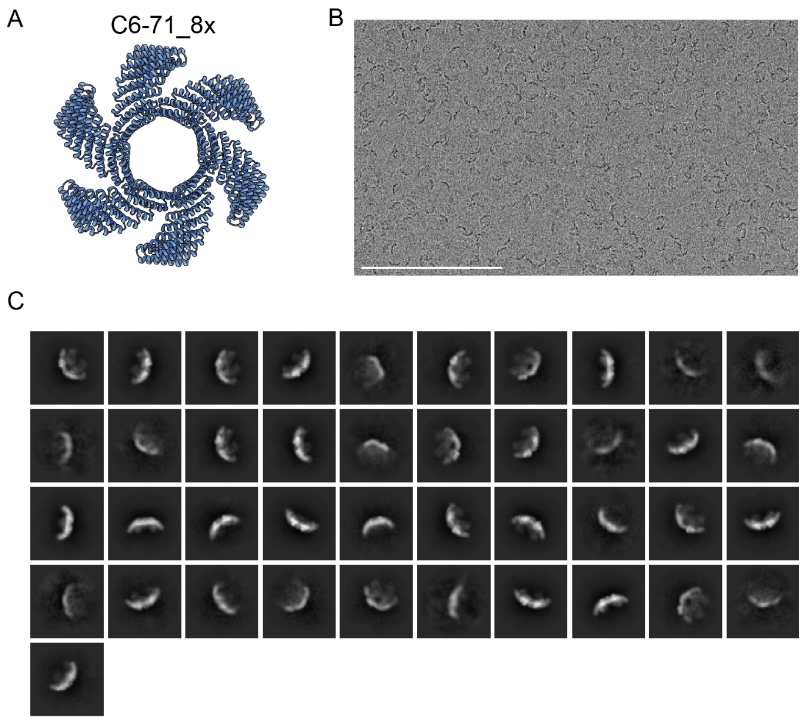


**Figure S13:** (A) C6-71_8x repeat extension design model (B) cryo-EM grid image (C) class averages. Scale Bar: 200 nm (B)

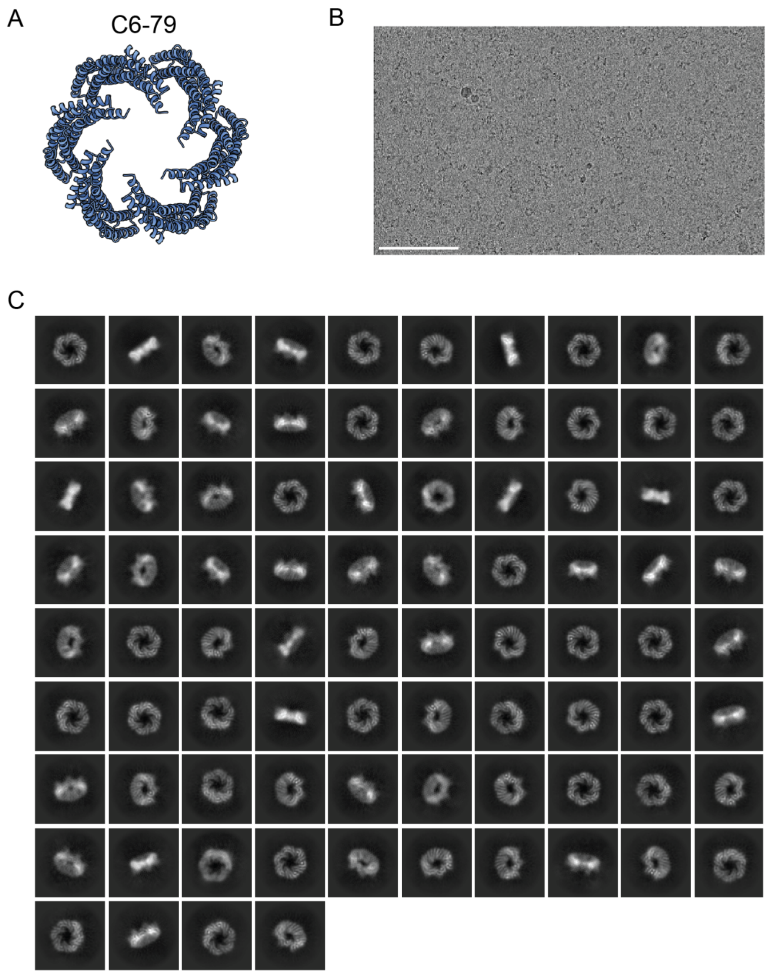


**Figure S14:** (A) C6-79 design model (B) cryo-EM grid image (C) class averages. Scale Bar: 100 nm (B)

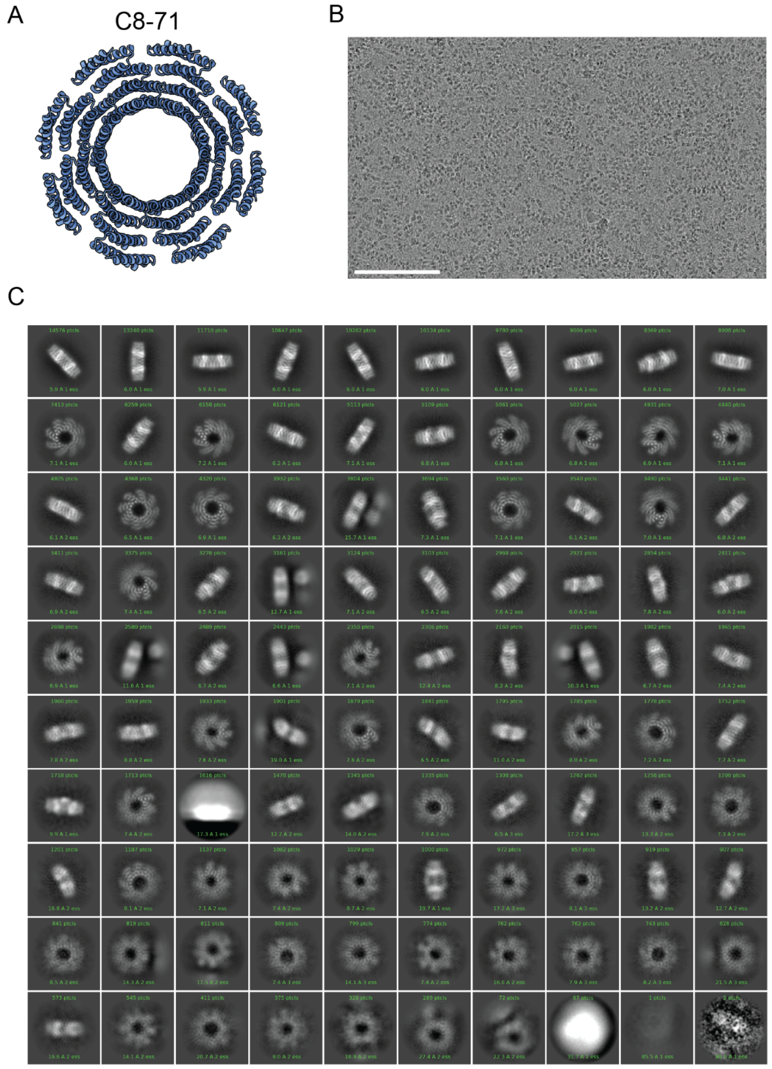


**Figure S15:** (A) C8-71 design model (B) cryo-EM grid image (C) class averages. Scale Bar: 100 nm (B)

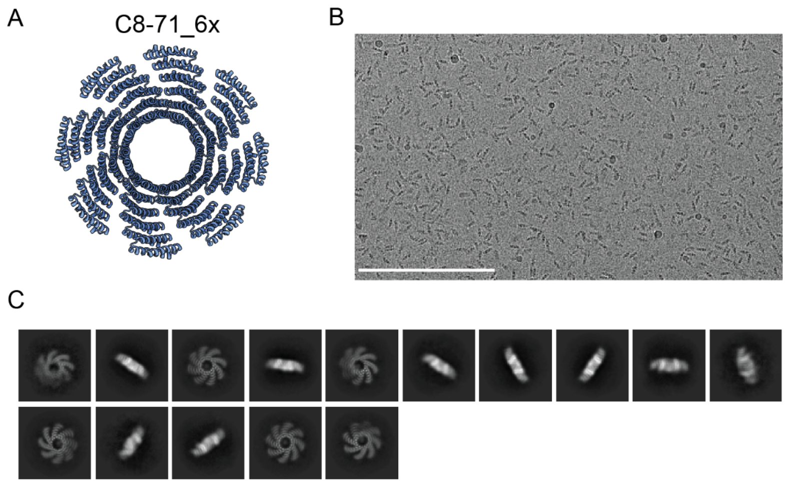


**Figure S16:** (A) C8-71_6 repeat extension design model (B) cryo-EM grid image (C) class averages. Scale Bar: 200 nm (B)

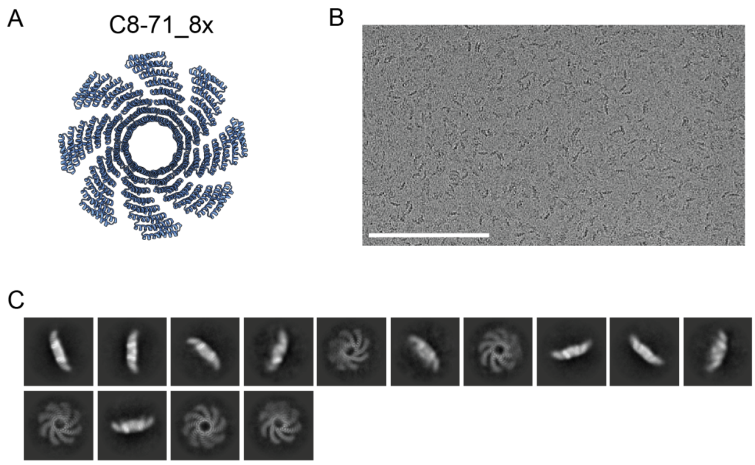


**Figure S17:** (A) C8-71_8 repeat extension design model (B) cryo-EM grid image (C) class averages. Scale Bar: 200 nm (B)

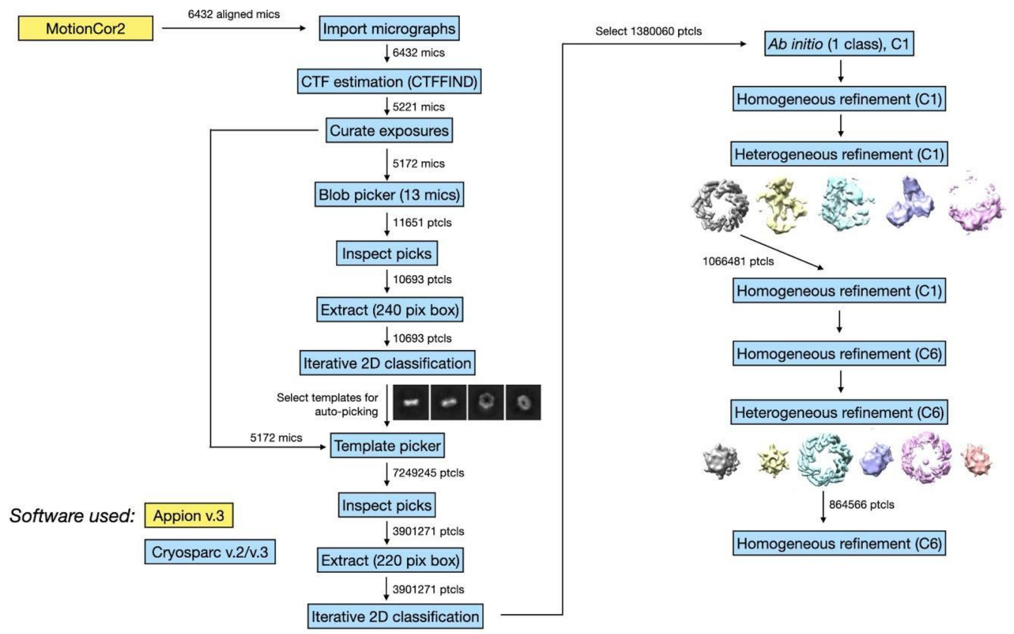


**Figure S18.** Cryo-EM processing workflow for C6-79

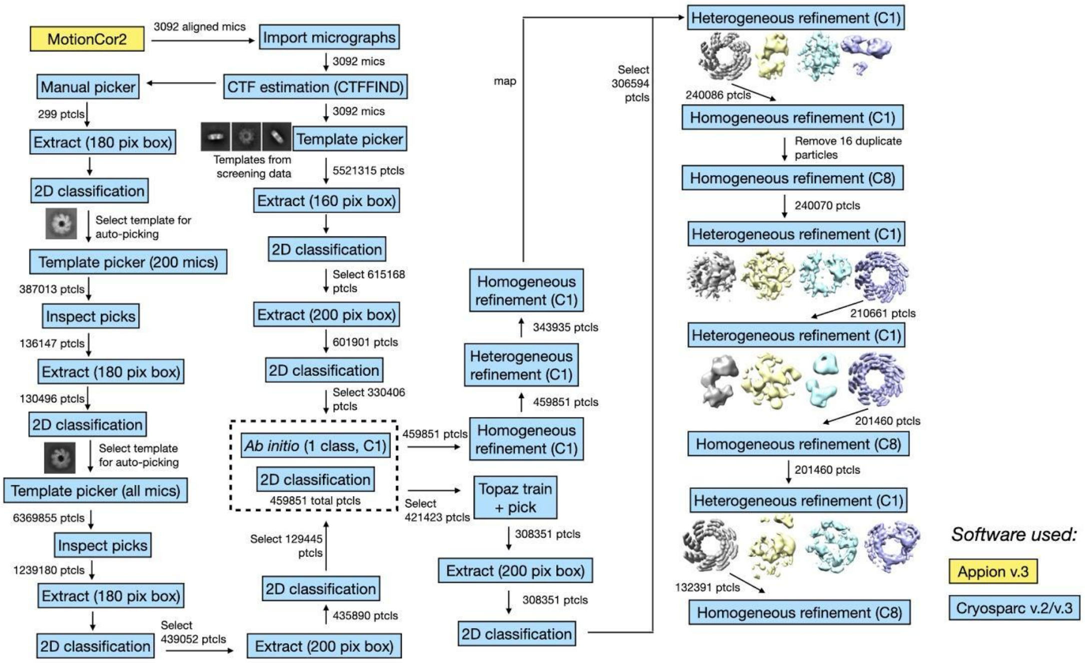


**Figure S19.** Cryo-EM processing workflow for C8-71

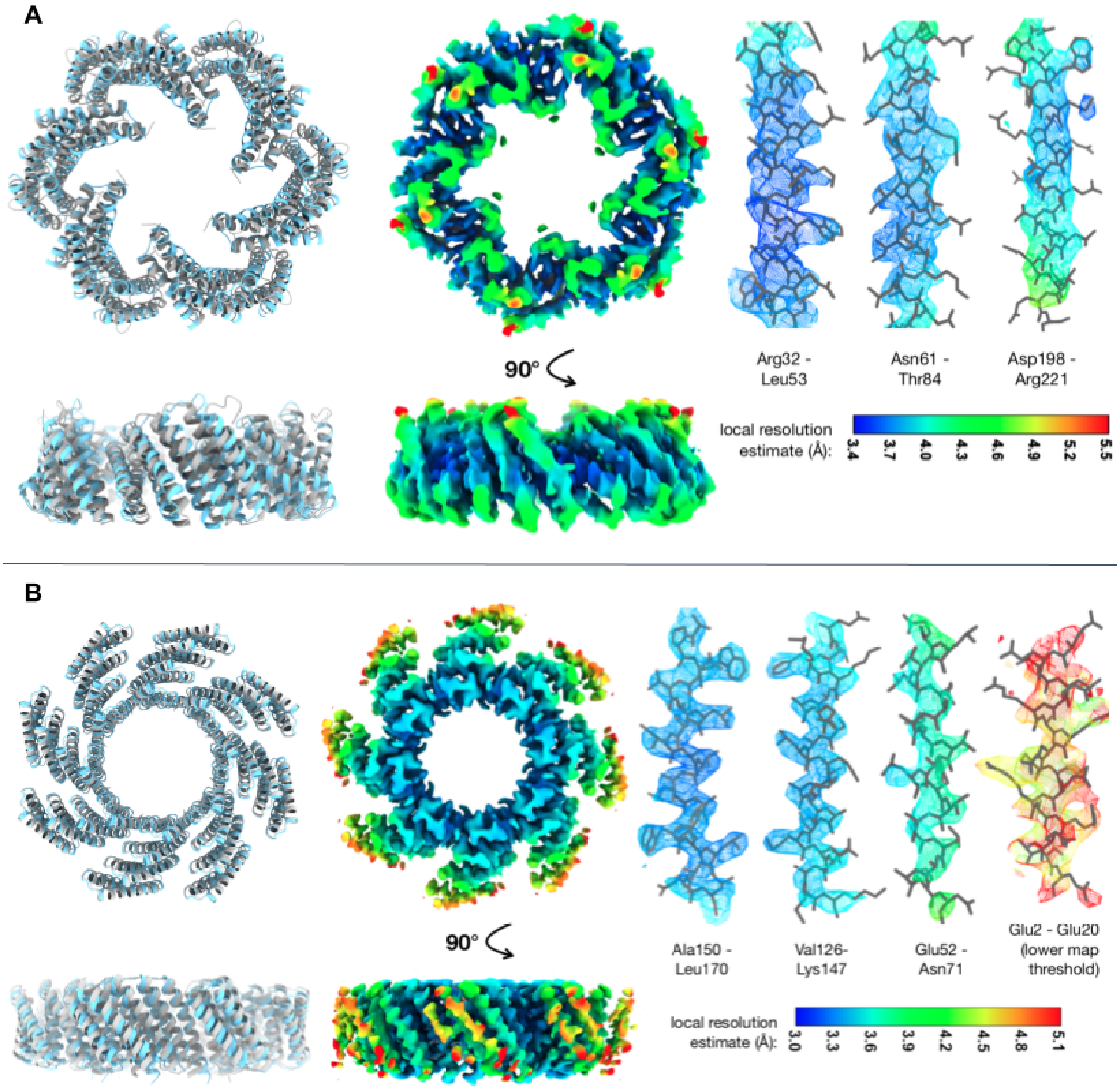


**Figure S20.** Overview of C6-79 (A) and C8-71 (B) cryo-EM models. *Left*: alignment of cryo-EM structures (cyan) with design models (grey). Middle: cryo-EM maps colored by local resolution. *Right*: helices from the cryo-EM models shown within the corresponding map regions, representing areas of the map with different local resolution. With the exception of the Glu2 - Glu20 helical region, the map segments for each design are shown at the same threshold.

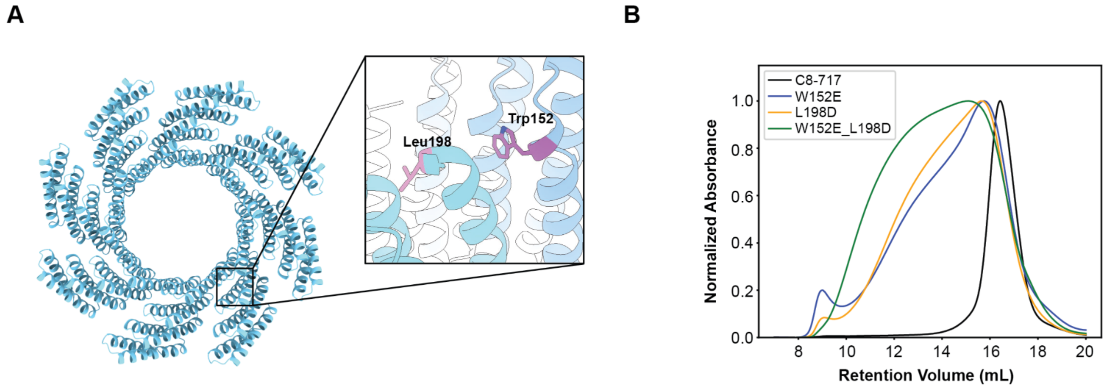


**Figure S21.** (A) C8-71 structure. Inset shows critical residues L198 and W152. (B) Mutations in interface residues W152 and L198 lead to aggregation of the C8-71 structure manifested by a broadening of the SEC peaks.

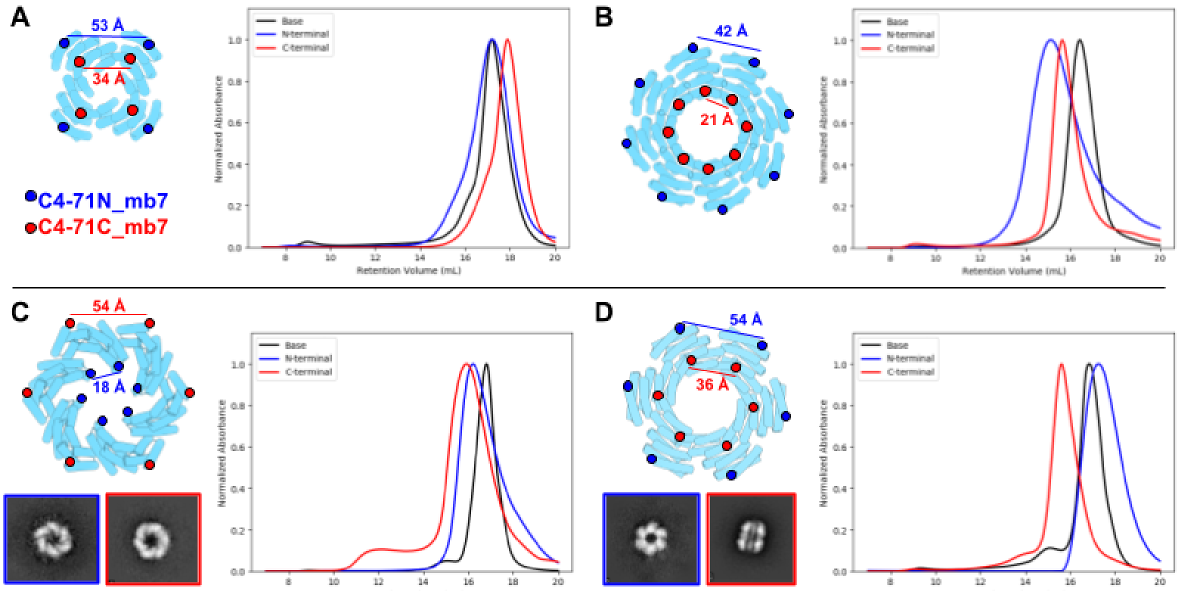


**Figure S22.** SEC Characterization of mb7 presenting oligomers. Red and blue circles indicate the N- or C-termini of the oligomers at which mb7 was flexibly fused to the structure. (A) C4-71 (B) C8-71 (C) C6-79 including top view class averages of N- (blue) and C-terminal (red) fusions. (D) C6-71 including top view classes of N- (blue) and C-terminal (red) fusions. C-terminal fusion leads to a macaron-like self-associating structure indicated in the class averages and shown by the large shift on SEC (red trace).

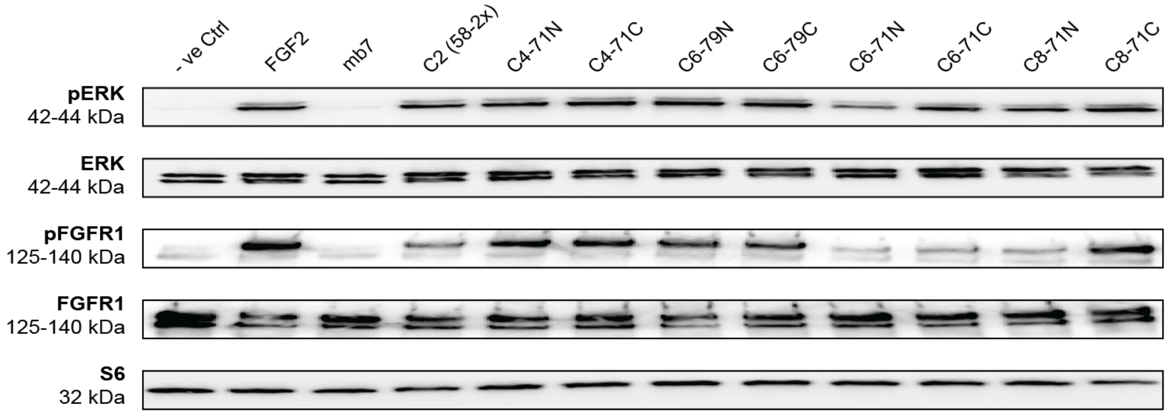


**Figure S23.** Western Blot analysis of pERK and pFGFR1 levels in CHO-R1c cells after treatment with 10 nM of agonists for 15 min and comparison with corresponding total protein levels. pERK, pFGFR1 and S6 data shown are the same as in Figure 4B and are shown here again for clarity.

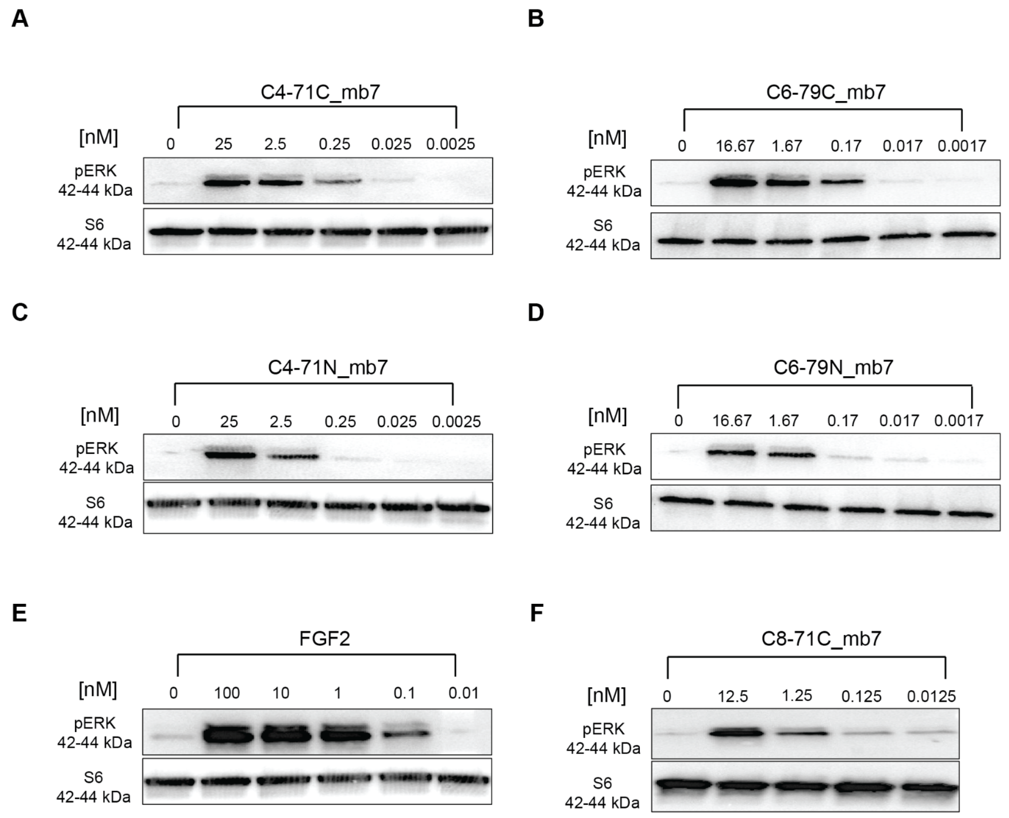


**Figure S24.** Western Blot titration analysis for selected designs. FGFR1c expressing CHO cells were treated with different concentrations of FGF2 or designed scaffolds for 15 min, followed by western blot analysis for phosphorylated ERK. (A) C4-71C_mb7, (B) C6-79C_mb7, (C) C4-71N_mb7, (D) C6-79N_mb7, (E) FGF2, (F) C8-71C_mb7.

A

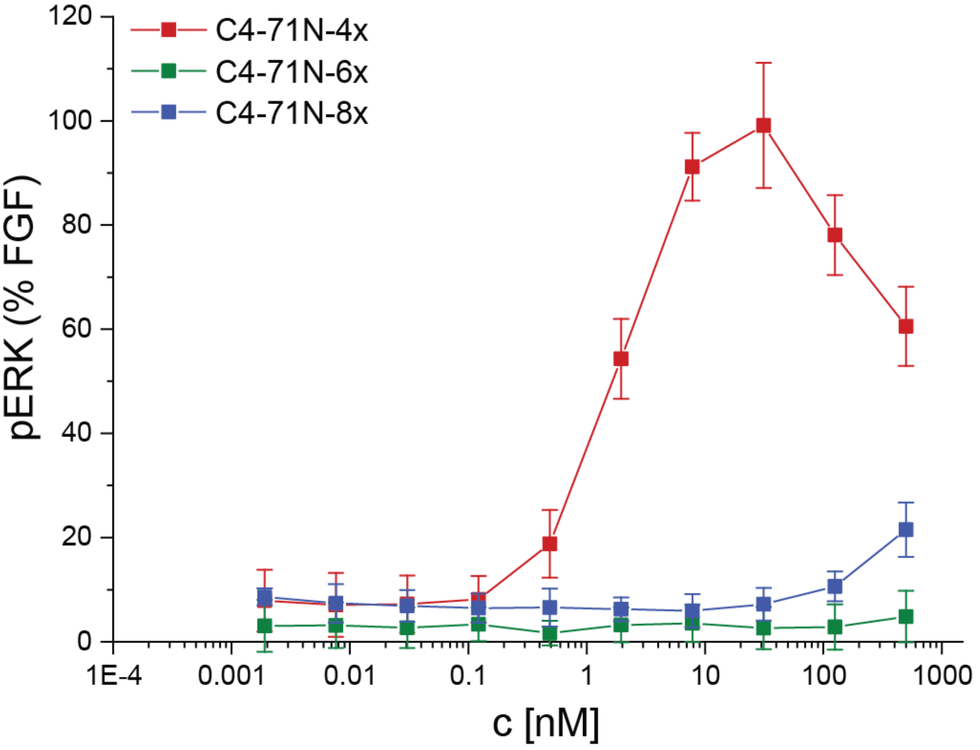


B

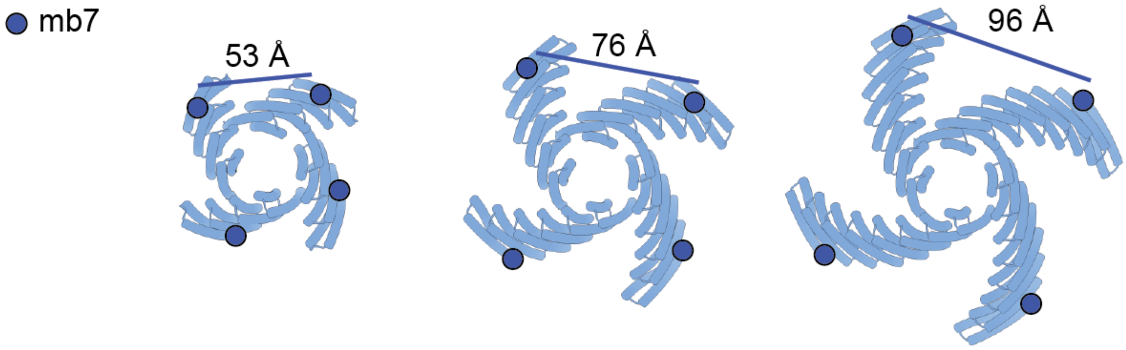


**Figure S25.** (A) Phosphoflow measurements of phosphorylated ERK for the C4-71N_mb7 extension series. (B) Cartoon models and distance measurements (N-terminus indicated with blue circle) of mb7 attachment points for the different extended constructs (C4-71N: 4-repeat, 6-repeat, 8-repeat).

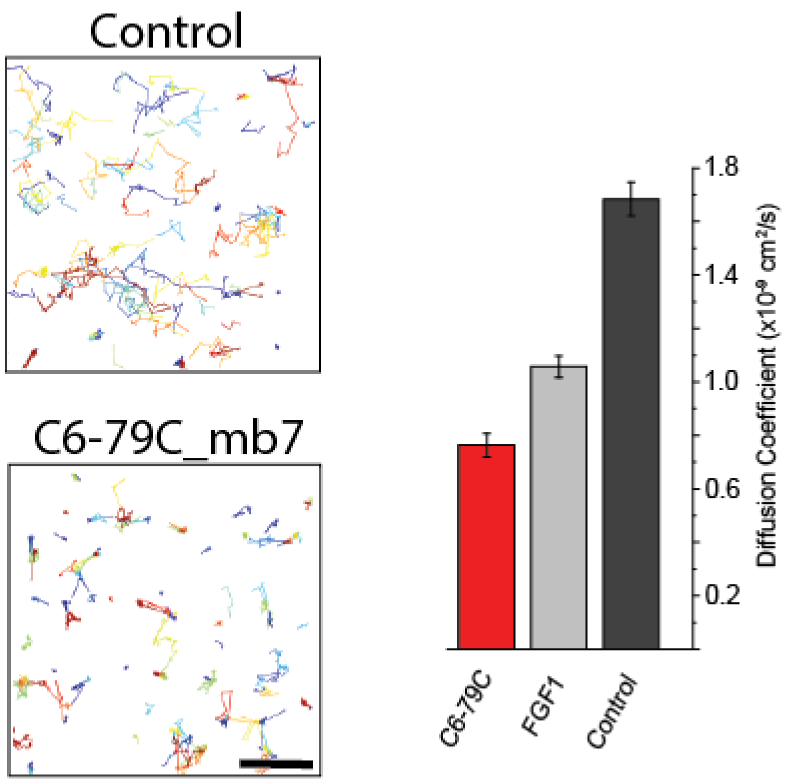


**Figure S26. Direct measurement of FGFR diffusion coefficients in the cell membrane.** Cells were treated with FGF1/heparin or C6-79C_mb7 and fluorescently tagged FGFR1c (HaloTag-R1c) revealed a strong decrease in the diffusion coefficient values in both conditions. The more pronounced effect of C6-79C_mb7 on the diffusion coefficient of FGFR1c may indicate a more robust FGFR1c clustering stimulated by the multivalent cyclic homo-oligomers. Diffusion coefficients were calculated from analyzing the labeled receptor tracks. Scale Bar: 2 μm

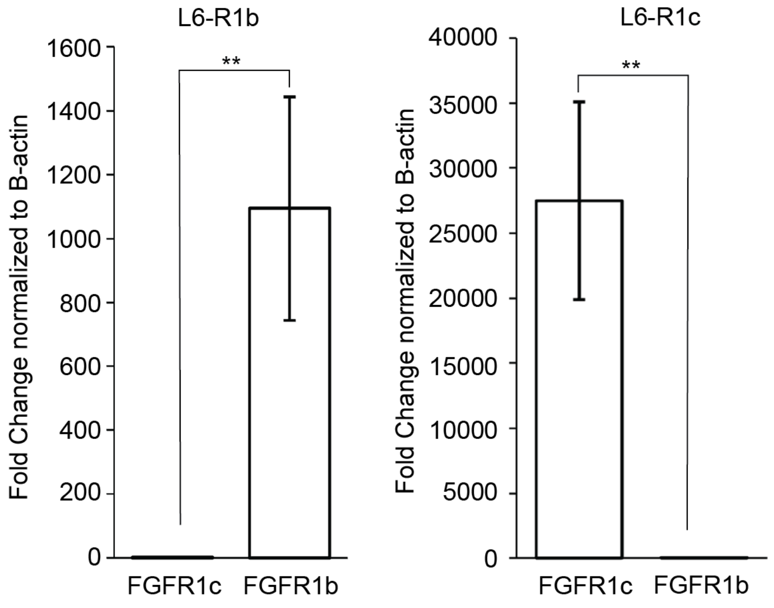


**Figure S27. qPCR validation of FGFR isoform overexpression cell lines.** Comparison of mRNA levels of FGFR isoforms (b or c) in L6 rat myoblast cells stably transfected with hFGFR1b (left) or hFGFR1c (right) via RT-qPCR. Error bars represent SEM from three independent biological repeats.

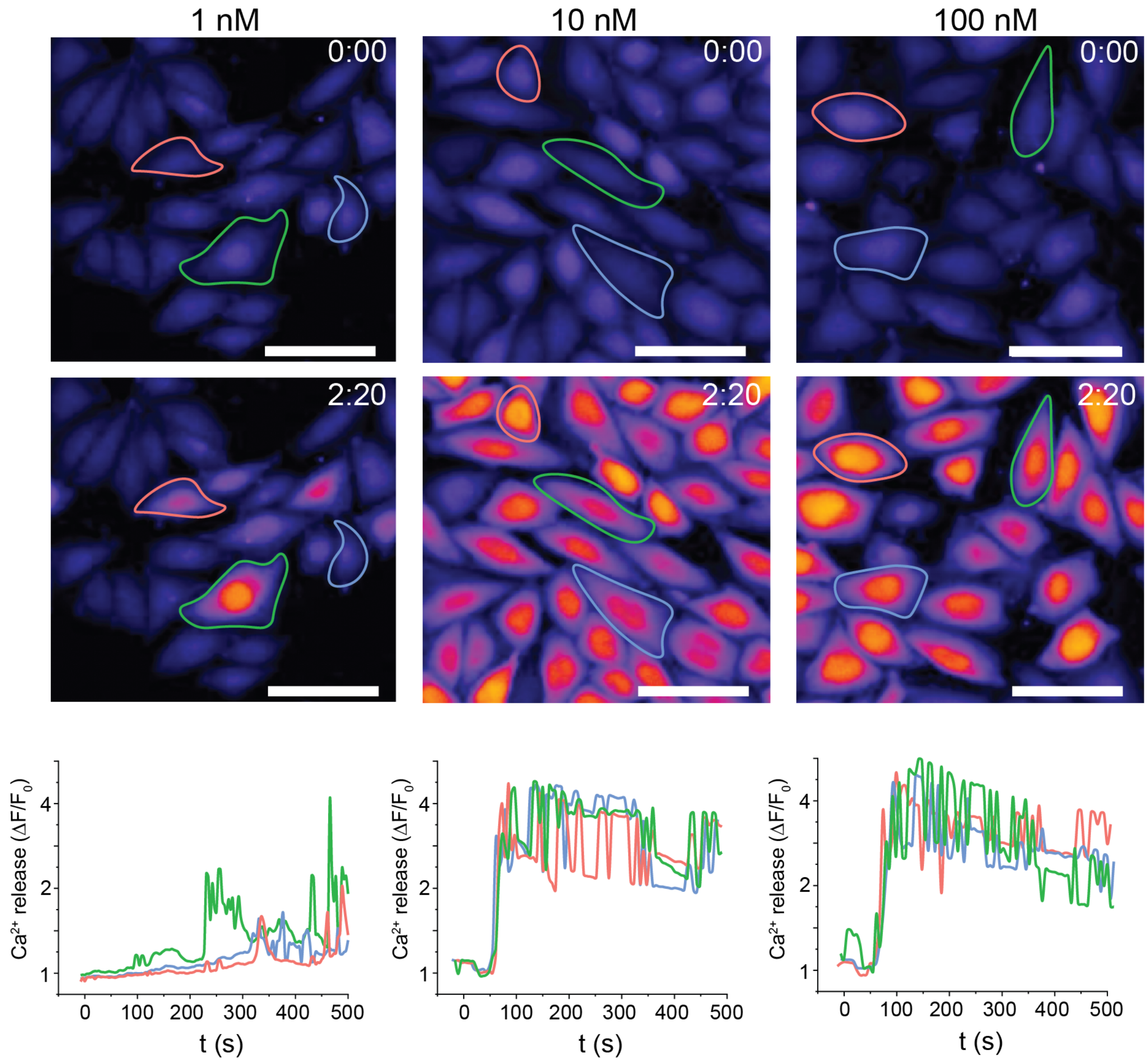


**Figure S28.** Normalized single cell fluorescence intensity measurements for three randomly selected cells tracking calcium release in CHO-R1c cells following treatment with different concentrations of C6-79C_mb7. Images show two different time points. Selected cells are outlined and their fluorescence intensity is tracked underneath. Scale Bar: 66.3 µm

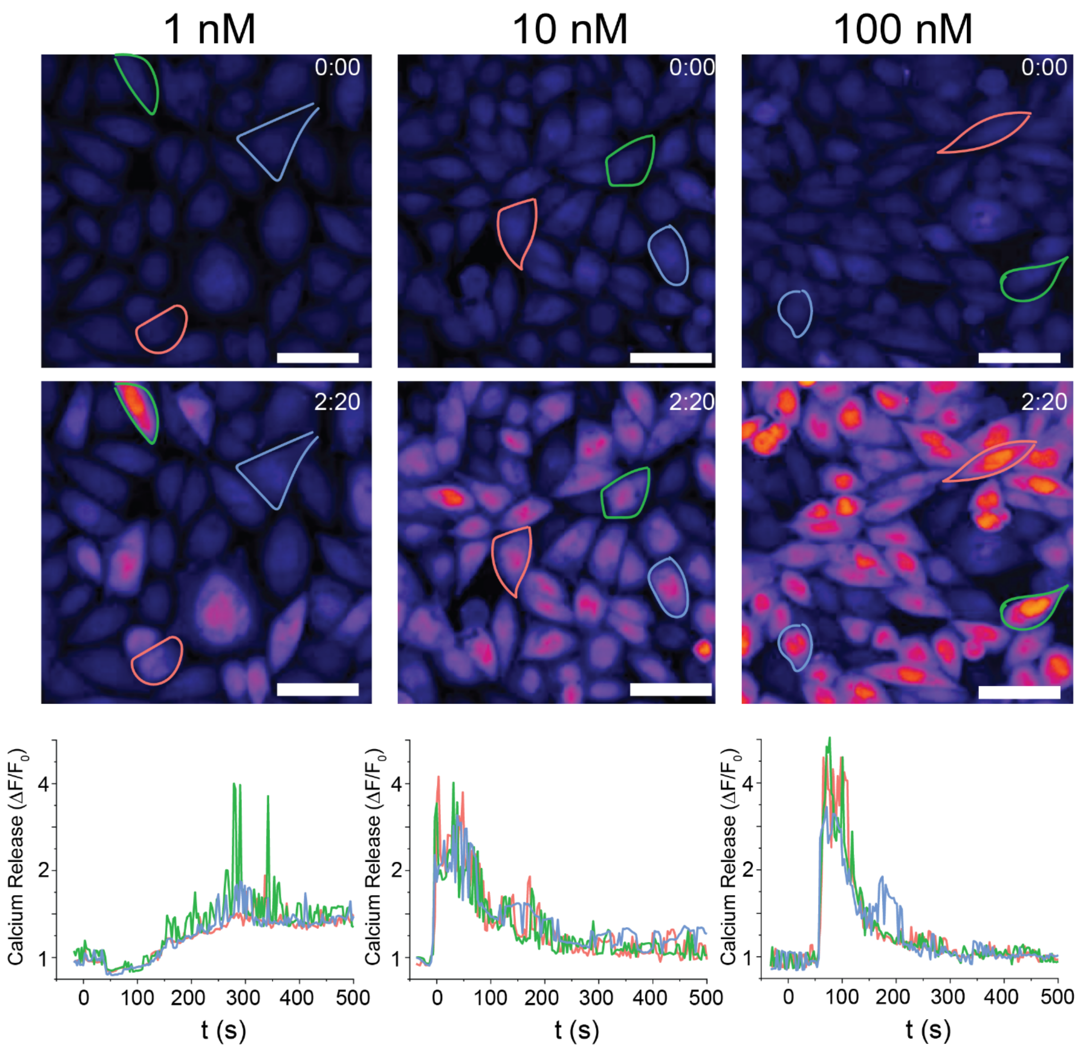


**Figure S29.** Normalized single cell fluorescence intensity measurements for three randomly selected cells tracking calcium release in CHO-R1c cells following treatment with different concentrations of native FGF2. Images show two different time points. Selected cells are outlined and their fluorescence intensity is tracked underneath. Scale Bar: 50 µm

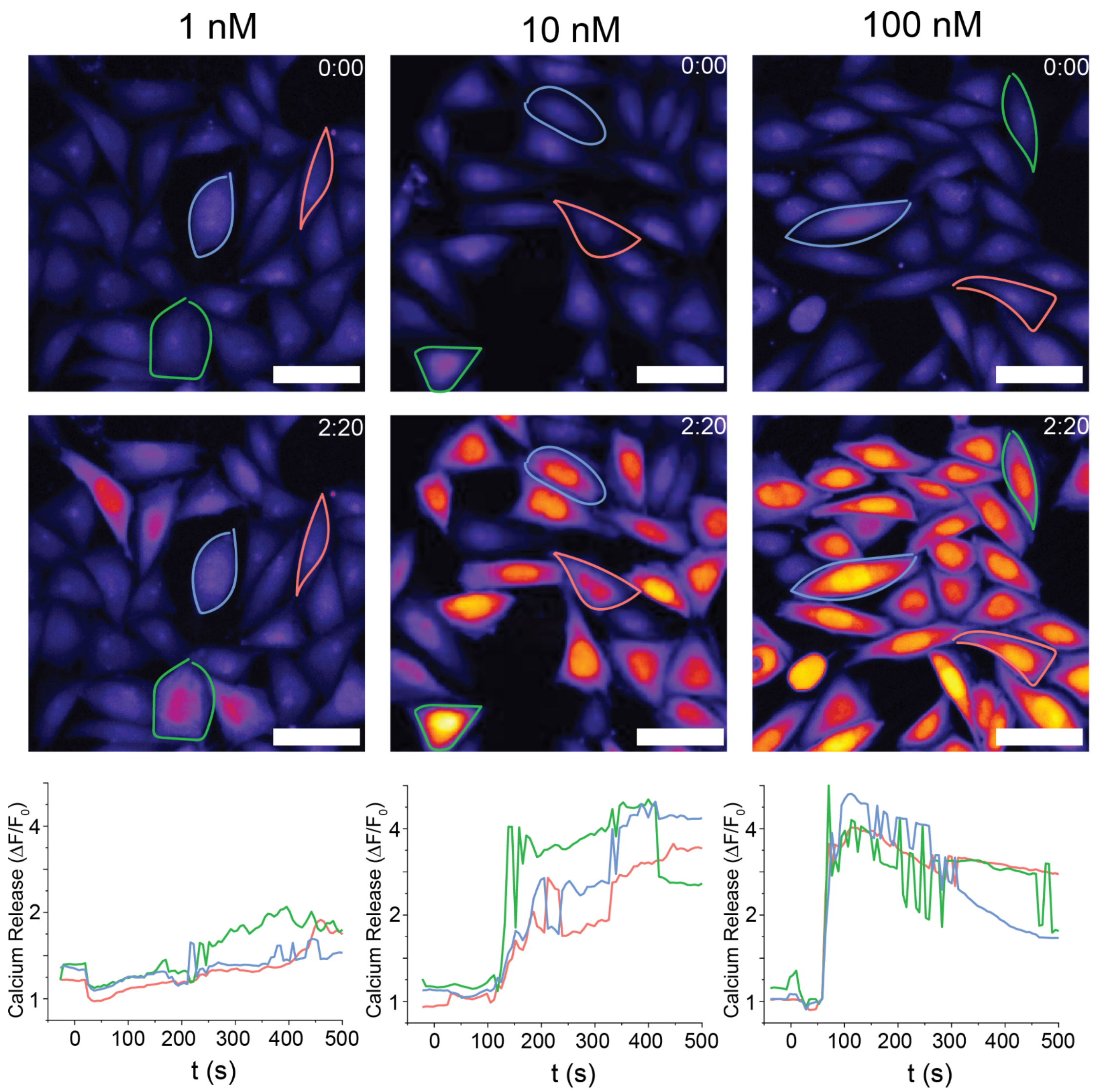


**Figure S30.** Normalized single cell fluorescence intensity measurements for three randomly selected cells tracking calcium release in CHO-R1c cells following treatment with different concentrations of native FGF2, along with 40 ug/mL of heparan sulfate proteoglycans. Images show two different time points. Selected cells are outlined and their fluorescence intensity is tracked underneath. Scale Bar: 66.3 μm

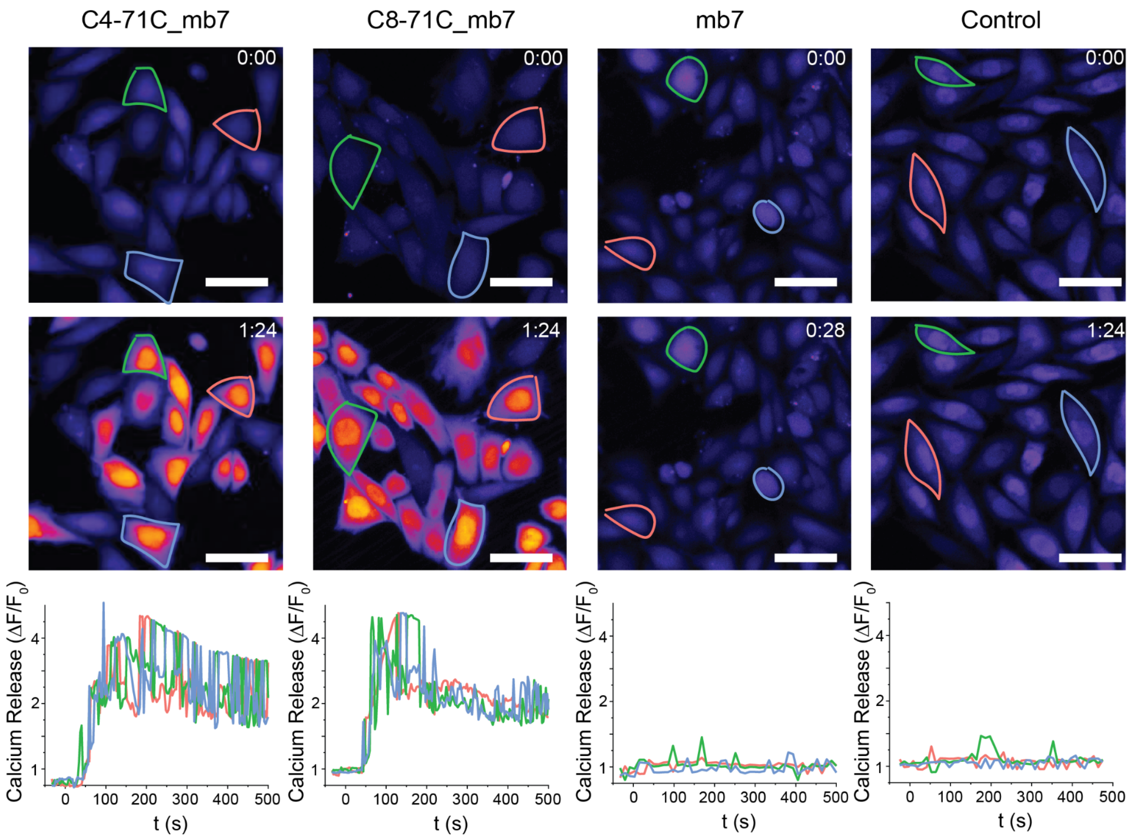


**Figure S31.** Normalized single cell fluorescence intensity measurements for three randomly selected cells tracking calcium release in CHO-R1c cells following treatment with 100nM of C4-71C_mb7, C8-71C_mb7 and mb7 alone. Images show two different time points. Selected cells are outlined and their fluorescence intensity is tracked underneath. Scale Bar: 50μm

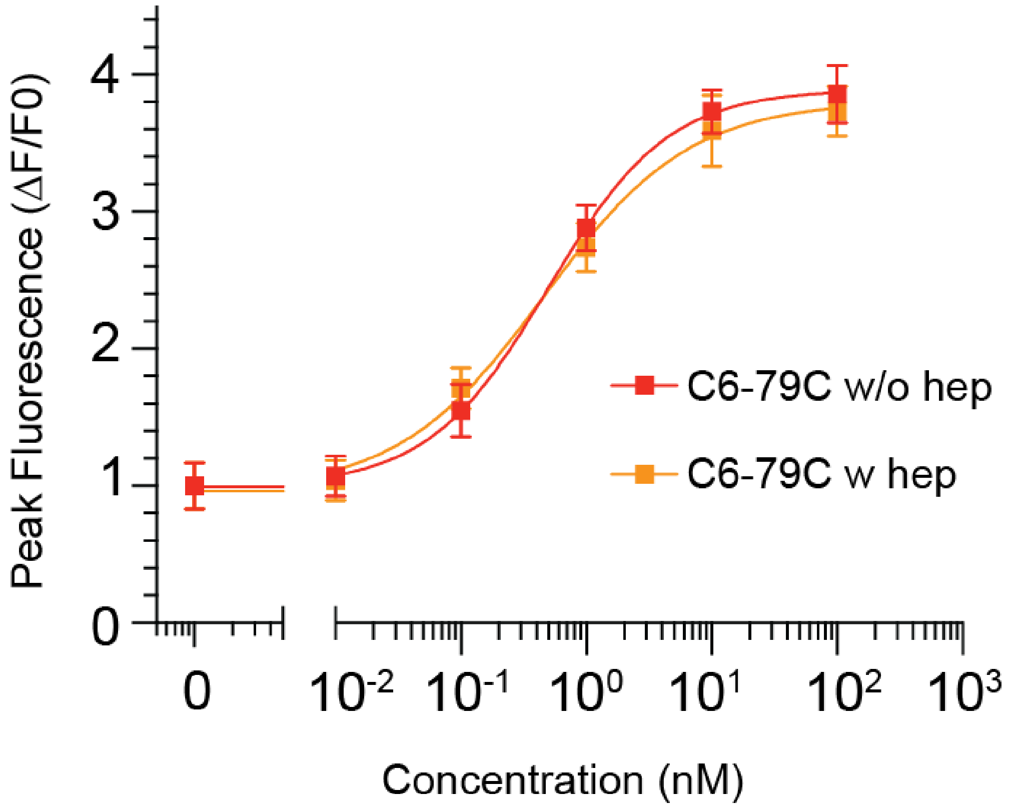


**Figure S32:** Titration curve for C6-79C_mb7 with and without heparin shows heparin independent $Ca^{2+}$ signaling of the designed agonist in CHO-R1c cells. 40µg/mL heparin was supplemented in each condition.

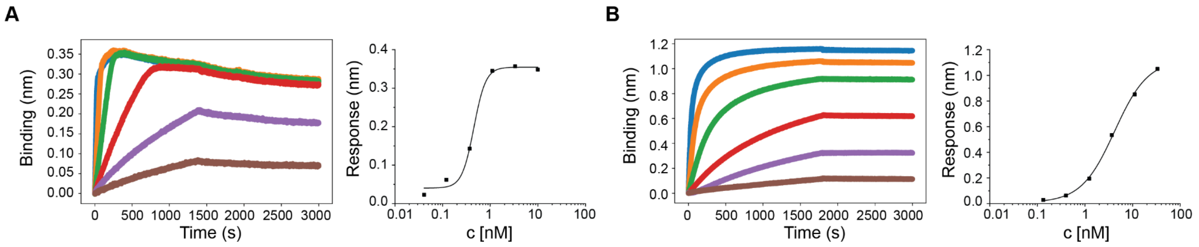


**Figure S33.** Biolayer-Interferometry (BLI) analysis of mb7 and C6-79C_mb7. (A) mb7 binding titration with highest concentration of 10 nM and subsequent 3 fold dilution steps (Blue: 10 nM, Orange: 3.33 nM, Green: 1.11 nM, Red: 0.37 nM, Purple: 0.12 nM, Brown: 0.04 nM). Right: Steady-state analysis yields a Kd of 453 pM. (B) C6-79C_mb7 binding titration with highest concentration of 30 nM and 3 fold dilution steps (Blue: 30 nM, Orange: 10 nM, Green: 3.33 nM, Red: 1.11 nM, Purple: 0.37 nM, Brown: 0.12 nM). Right: Steady-state analysis yields a Kd of 4.1 nM. Note that fusing the minibinder to the scaffold decreased binding affinity. We did not measure avidity effects with the low density of biotinylated receptors on the Octet tip.

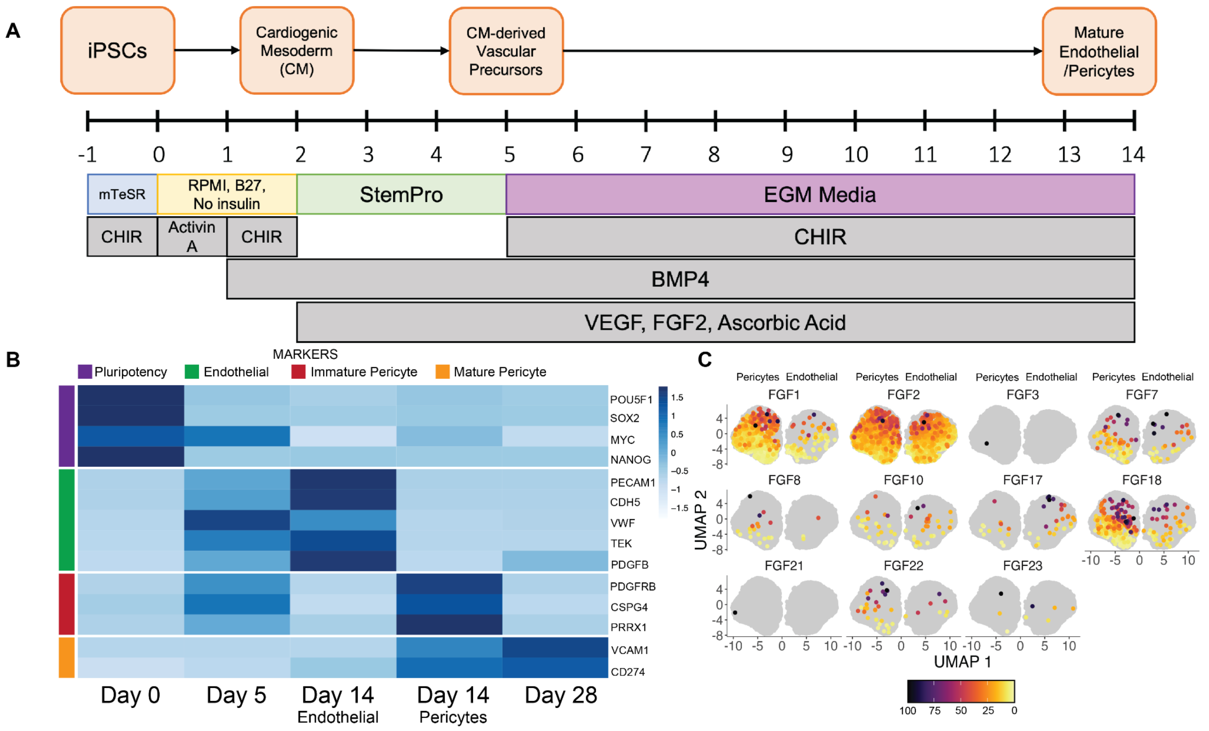


**Figure S34.** Vascular differentiation using designed agonists. (A) Schematic for endothelial cell differentiation from iPSCs through a cardiogenic mesoderm intermediate. (B) Clustered heatmap comparing the normalized expression of selected putative cell markers (iPSCs, endothelial and pericyte) across all obtained clusters along with given annotations. (C) Gene expression density plots of selected FGFs highlighting endogenous expression in both cell types obtained at Day 14.

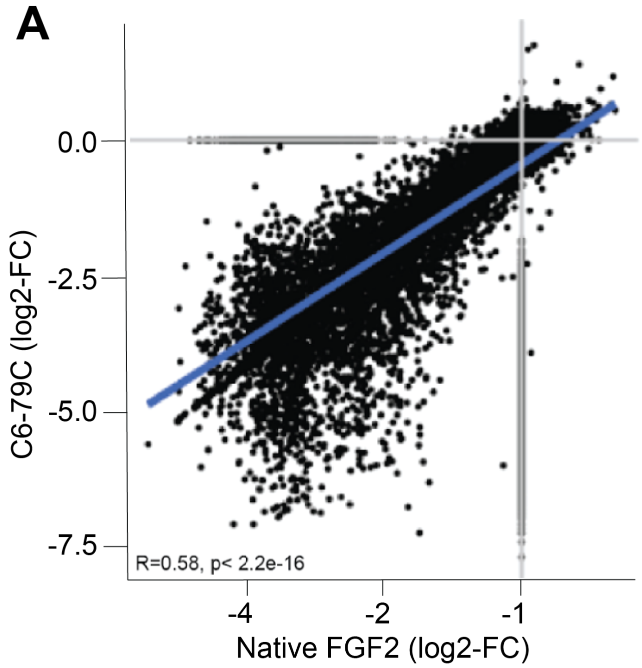


**Figure S35:** Bulk transcriptomics using designed agonists. (A) Correlation between differentially expressed genes (DEGs) in HUVEC cells treated with either native FGF2 or C6-79C_mb7.

### Tables

#### Supplementary Tables

**Supplementary Table I.** Designed monomeric repeat proteins used as building blocks and success rate.

| Name | PDB ID | Reference | Designs tested | Successful designs |
|---|---|---|---|---|
| DHR1 | | Brunette et al. 2015 | 2 | 0 |
| DHR4 | 5CWB | Brunette et al. 2015 | 5 | 1 |
| DHR5 | 5CWC | Brunette et al. 2015 | 5 | 0 |
| DHR8 | 5CWF | Brunette et al. 2015 | 5 | 0 |
| DHR10 | 5CWG | Brunette et al. 2015 | 8 | 0 |
| DHR14 | 5CWH | Brunette et al. 2015 | 6 | 0 |
| DHR18 | 5CWI | Brunette et al. 2015 | 2 | 1 |
| DHR49 | 5CWJ | Brunette et al. 2015 | 6 | 0 |
| DHR53 | 5CWK | Brunette et al. 2015 | 1 | 0 |
| DHR54 | 5CWL | Brunette et al. 2015 | 6 | 0 |
| DHR71 | 5CWN | Brunette et al. 2015 | 19 | 5 |
| DHR76 | 5CWO | Brunette et al. 2015 | 7 | 0 |
| DHR79 | 5CWP | Brunette et al. 2015 | 6 | 1 |
| DHR81 | 5CWQ | Brunette et al. 2015 | 9 | 1 |
| TJ116 | 6W2R | Brunette et al. 2020 | 1 | 0 |
| TJ120 | 6W2V | Brunette et al. 2020 | 4 | 0 |
| TJ121 | 6W2W | Brunette et al. 2020 | 1 | 0 |
| TJ131 | 6W2Q | Brunette et al. 2020 | 2 | 1 |

**Supplementary Table II.** Cryo-EM data acquisition

| | **C4-131** | **C4-71** | **C4-71_6x** | **C4-71_8x** | **C4-81** | **C6-71** | **C6-71_6x** | **C6-71_8x** | **C6-79** | **C8-71** | **C8-71_6x** | **C8-71_8x** |
|---|---|---|---|---|---|---|---|---|---|---|---|---|
| Microscope | Arctica (NYU) | Krios (NCCAT) | Arctica (NYU) | Arctica (NYU) | Krios (NCCAT) | Arctica (NYU) | Arctica (NYU) | Arctica (NYU) | Krios (NCCAT) | Krios (NCCAT) | Arctica (NYU) | Arctica (NYU) |
| Voltage (kV) | 200 | 300 | 200 | 200 | 300 | 200 | 200 | 200 | 300 | 300 | 200 | 200 |
| Exposure navigation | Stage position | Image shift | Stage position | Stage position | Image shift | Stage position | Stage position | Stage position | Image shift | Image shift | Stage position | Stage position |
| Nominal magnification | 36,000X | 105,000X | 36,000X | 36,000X | 105,000X | 36,000X | 36,000X | 36,000X | 81,000X | 81,000X | 36,000X | 36,000X |
| Detector | Gatan K3 | Gatan K3 | Gatan K3 | Gatan K3 | Gatan K3 | Gatan K3 | Gatan K3 | Gatan K3 | Gatan K3 | Gatan K3 | Gatan K3 | Gatan K3 |
| Detector mode | super-resolution | super-resolution | super-resolution | super-resolution | super-resolution | super-resolution | super-resolution | super-resolution | counting | counting | super-resolution | super-resolution |
| Unbinned pixel size (Å/pix) | 0.548 | 0.4124 | 0.548 | 0.548 | 0.4124 | 0.548 | 0.548 | 0.548 | 0.5346 | 0.5346 | 0.548 | 0.548 |
| No. of frames | 40 | 40 | 40 | 40 | 40 | 40 | 40 | 40 | 50 | 50 | 40 | 40 |
| Dose rate (e-/Å/s) | 20.33 | 29.23 | 20.37 | 20.37 | 29.23 | 20.37 | 20.37 | 20.28 | 24.52 | 24.52 | 21.89 | 21.89 |
| Exposure per frame (e-/Å$^2$) | 1.423 | 1.169 | 1.426 | 1.426 | 1.169 | 1.426 | 1.426 | 1.419 | 1.226 | 1.226 | 1.314 | 1.314 |
| Total dose (e-/Å$^2$) | 56.91 | 46.77 | 57.04 | 57.04 | 46.77 | 57.04 | 57.04 | 56.77 | 61.3 | 61.3 | 52.54 | 52.54 |
| Exposure time (s) | 2.8 | 1.6 | 2.8 | 2.8 | 1.6 | 2.8 | 2.8 | 2.8 | 2.5 | 2.5 | 2.4 | 2.4 |
| Defocus range (μm) | 2.1 - 2.9 | 0.3 - 1.3 | 2.1 - 2.9 | 2.1 - 2.9 | 0.8 - 2.5 | 2.1 - 2.9 | 2.1 - 2.9 | 1.1 - 2.9 | 0.8 - 2.5 | 0.8 - 2.5 | 2.0 - 3.0 | 2.0 - 3.0 |
| No. of micrographs | 26 | 5,221 | 121 | 127 | 18,412 | 81 | 87 | 223 | 6,434 | 3,092 | 263 | 96 |

**Supplementary Table III.** Cryo-EM data processing

| | **C4-131** | **C4-71** | **C4-71_6x** | **C4-71_8x** | **C4-81** | **C6-71** | **C6-71_6x** | **C6-71_8x** | **C6-79** | **C8-71** | **C8-71_6x** | **C8-71_8x** |
|---|---|---|---|---|---|---|---|---|---|---|---|---|
| Software used | Appion v.3, Cryosparc v.3 | Appion v.3, Cryosparc v.2/v.3, RELION v.3 | Appion v.3, Cryosparc v.3 | Appion v.3, Cryosparc v.3 | Appion v.3, Cryosparc v.2/v.3, RELION v.3 | Appion v.3, Cryosparc v.3 | Appion v.3, Cryosparc v.3 | Appion v.3, Cryosparc v.3 | Appion v.3, Cryosparc v.2/v.3 | Appion v.3, Cryosparc v.2/v.3 | Appion v.3, Cryosparc v.3 | Appion v.3, Cryosparc v.3 |
| No. of extracted particles | 60,555 | 3,355,984 | 9,276 | 7,265 | 1,878,998 | 16,793 | 15,332 | 93,046 | 3,901,271 | 308,351 | 28,236 | 62,332 |
| No. of final particles | 13,986 | 107,483 | 4,919 | 2,694 | 259,010 | 4,593 | 7,460 | 23,855 | 864,566 | 132,391 | 9,092 | 3,584 |
| Box size (pixels) | 150 | 212 | 200 | 220 | 256 | 200 | 220 | 300 | 220 | 200 | 256 | 256 |
| Symmetry Imposed | C1 | C4 | C4 | C4 | C4 | C6 | C6 | C6 | C6 | C8 | C8 | C8 |
| Global map resolution (FSC 0.143 unmasked): | > 9 Å | > 5 Å | > 9 Å | > 13 Å | > 5 Å | > 9 Å | > 8 Å | > 6 Å | > 4 Å | > 4 Å | > 7 Å | > 7 Å |
| Map sharpening B factor ($Å^2$) | N/A | N/A | N/A | N/A | N/A | N/A | N/A | -166.2 | -313.2 | -223.3 | N/A | N/A |
| EMDB ID: | EMD-28958 | EMD-28974 | EMD-28966 | EMD-28967 | EMD-28973 | EMD-28968 | EMD-28969 | EMD-28970 | EMD-28889 | EMD-28888 | EMD-28971 | EMD-28972 |

**Supplementary Table IV.** Model statistics for C6-79 and C8-71 cryo-EM structures

| | **C6-79** | **C8-71** |
|---|---|---|
| Global map resolution (Å, FSC 0.143 unmasked / masked) | 4.8 / 4.0 | 4.3 / 3.6 |
| Sphericity from 3DFSC (unmasked/masked): | 0.92 / 0.95 | 0.82 / 0.98 |
| Map CC (mask) | 0.814 | 0.824 |
| Map CC (volume) | 0.808 | 0.813 |
| Map CC (peaks) | 0.709 | 0.696 |
| R.m.s. deviations (bonds) | 0.005 | 0.006 |
| R.m.s. deviations (angles) | 0.728 | 0.711 |
| Ramachandran plot values (%) | | |
| outliers | 0.00 | 0.00 |
| allowed | 1.38 | 2.35 |
| favored | 98.62 | 97.65 |
| Rotamer outliers (%) | 0.00 | 0.00 |
| C-beta deviations (%) | 0.00 | 0.00 |
| CaBLAM outliers (%) | 0.93 | 1.03 |
| Overall score (Molprobity[77]) | 1.75 | 1.87 |
| Clashscore | 17.80 | 20.00 |
| PDB ID | 8F6R | 8F6Q |

**Supplementary Table V.** Sequences for proteins in this study. Sequences include N-terminal Met +/- Gly and select sequences contain a MEKKI expression tag (DNA sequence: atggagaaaaaatc). 6xHis tags are either C-terminal or N-terminal with Gly-Ser linker. All linkers and tags are underlined. Genes were expressed in pET29b+.

| **Name** | **Sequence** |
|---|---|
| FGF_mb7 | MGDRRKEMDKVYRTAYKRITSTPDKEKRKEVVKEATEQLRRIAKDEEEKKKAAYMISFLKTLG LEHHHHHH |
| FGF_mb7 _mCherry | MVSKGEEDNMAIIKEFMRFKVHMEGSVNGHEFEIEGEGEGRPYEGTQTAKLKVTKGGPLPFA WDILSPQFMYGSKAYVKHPADIPDYLKLSFPEGFKWERVMNFEDGGVVTVTQDSSLQDGEFI YKVKLRGTNFPSDGPVMQKKTMGWEASSERMYPEDGALKGEIKQRLKLKDGGHYDAEVKT TYKAKKPVQLPGAYNVNIKLDITSHNEDYTIVEQYERAEGRHSTGGMDELYKGGSGGSGDRR |

| | |
|---|---|
| | KEMDKVYRTAYKRITSTPDKEKRKEVVKEATEQLRRIAKDEEEKKKAAYMISFLKTLGLEHHHHHH |
| C2-58 | MDEELLRELLKLLLKLLEQMGDEEARRVVEELREELEKKGDPRALVLAFALVILVFLLRILRELGDEELVRRVEELWEELLKEGDPQAMMEVFKLVQELQERRLEHHHHHH |
| C2-CDX | MPKKQLMKLLFKVLEALFRGDEETLRELAREAVELAERLLKLGDPELLFLALAIAIIVAWAVGDEELLKRLAQIIKELLKRAEELGDPDLRRLIEELVEFVERLLEHHHHHH |
| C2-Y2D | MGEELLQEVARVLLKLAQELGDPDVERVVRELLERLERKGDPRIVIRILLLLVALLLLWIARELGDPEVVRELEELLKRLIKKGDPRLFAEILRIVLELEEEVGLEHHHHHH |
| C4-18 | MGSIEKLCKKAESEAREARSKAEELRQRHPDSQAARDAQKLASQAEEAVKLACELAQEHPNAWIARACIRAASEAAEAASKAAELAQRHPDSKAARDAIKLASQAAEAVKLACELAQEHPNADIAELCILAAWAAARAASLAAELAQRHPDLWAANLAIRLASQAAEAVKLACELAQEHPNAEIARECIWLAWEAALLAALAAEEAQRHPNDIRAMLLFIEAIRKAEEVKKRCERGSLEHHHHHH |
| C4-717 | MGPEEILERAKESLERAREASERGDEEEFRKAAEKALELAKRLVEQAKKEGDPELVLEAAKVALRVAELAAKNGDKEVFKKAAESALEVAKRLVEVASKEGDPELVLEAAKVALRVAELAAKNGDKEVFKKAARSALWVAFILVKVALKEGDPELVEEAAKVAIRVFELAWEQGDEDVLRLALLTMIVVLILLILVLLKKGGWGSLEHHHHHH |
| C4-71 | MGPEEILERARESLERAREASERGDEEEFRKAAEKALELAKRLVEQAKKEGDPWMVMWAALVALWVALLALRNGDKEVFKKAAESALEVAKRLVEVASKEGDPEMVLLAAWVALFVAWLAWLFGDKEVFKKAAESALEVAKRLVEVASKEGDPELVEEAAKVAEEVEKLAEKQGDEEVREKAWETWMEVWLLWLEVRLRKGGGGSLEHHHHHH |
| C4-81 | MGELERESREAEKRLKEARLFAWAARLLGDLKLLAKALIEEARAVQELARVACERGNRDEAWDAFEKALEVFEEAVKVSEEAREQGDDEVLALALIAIALAVLALAEVACCLGISELAELAWKMAEWVLEEARKVSEEAREQGDDEVLALALIAIALAVLALAEVACCRGNKEEAERAYEDARRVEEEARKVKESAEEQGDSEVKRLAEEAEQLAREARRHVQECRGGWLEHGSLEHHHHHH |
| C4-131 | MGLKELLKRAEELAKSPDPEDLKEAVRLAEEVVRERPGSEAAKKALEIIQEAAEKLKKSPDPEAIIAAARALLKIAATTGDNEAAKQAIEAASKAAQLAEQRGDDELVCEALALLIAAQVLLLKQQGVPMLEVAIHVAETILQILQRLKRKGASEEVRKECLKRILREIAEALQRSGVPEEEIALIMLLIILLLMMLGSLEHHHHHH |
| C6-4 | MGDECEKKAREVALRVLVLWAKGTSEDEIAEEVAREISEVIRTLKESGSSYEVICECVARIVAFIVEVLVLMGTSEDEIAEIVARVISEVIRTLKESGSSYEVICKCVAFIVAEIVEALKRAGTSEDEIAEIVARVISEVIRTLKESGSSEDIIWECIMLIMIFIAEALLRSGTSEDEIREILRRVRSEVERTLKESGSGSLEHHHHHH |
| C6-714 | MGPEEILERAKESLERAREASERGDEEEFRKAAEKALELAKRLVEQAKKEGDPELVLEAAKVALRVAELAAKNGDKEVFKKAAESALEVAKRLVEVASKEGDPELVLEAAKVALEVARLAAENGDKEVFKKAAESALEVAAKLVWVAMKEGDPRMVINALMVALWVLLLAFLQGDEEVFERARTLFELVRNFIEALEMREGGGGSLEHHHHHH |
| C6-71 | MGPEEILERAKESLERAKEAFERGDEEEFRKAAEKALELAKRLVEQAKKEGDPELVFEAARVALWVAWLAAWFGDKEVFKKAAESALEVAKRLVEVAKEEGDPELVLKAAFVALLVAIMAVILGDKEVFKKAAESALEVAKRLVEIAAREGDPELVEEAAKVAELVRELAKLMGDEEVYEKARETAREVRLFLLFVRIWEGGGGSLEHHHHHH |
| C6-79 | MGSSDEEEARELEERAREAAKRAIEAAKRTGDPRVRELAEELVKLAIWAAVEVWLDPSSSDVNEALKLIVEAIEAAVRALEAAERTGDPEVRELARELVRLAVEAAEEVQRNPSSSDVNEALKLIVIAIEAAVRALEAAERTGDPEVRELARELVRLAVEAAEEVQRNPSSEEVNEALRKIIKLILFAVMVLELAEEIGDPTWREMARRAVREAVELAEEVQRDPSGWLGHGSLEHHHHHH |

| | |
|---|---|
| C8-71 | MGPEEILERAKESLERAREASERGDEEEFRKAAEKALELAKRLVEQAKKEGDPELVLEAARVALWVAELAAKNGDKEVFKKAAESALEVAKRLVEVASKEGDPDLVAWAALVALWVAFLAFLNGDKEVFKKAAESALEVAKALMEVAMKVGAPWLVELAIAVARAVWLLAELFGDEEVRRRAEAFEIILRIAAIAVKAWLGGGGSLEHHHHHH |
| C4-71-6x | MGPEEILERAKESLERAREASERGDEEEFRKAAEKALELAKRLVEQAKKEGDPELVLEAAKVALRVAELAAKNGDKEVFKKAAESALEVAKRLVEVASKEGDPELVLEAARVALEVARLAAENGDKEVFKKAAESALEVAKRLVEVASKEGDPWMVMWAALVALWVALLALRNGDKEVFKKAAESALEVAKRLVEVASKEGDPEMVLLAAWVALFVAWLAWLFGDKEVFKKAAESALEVAKRLVEVASKEGDPELVEEAAKVAEEVEKLAEKQGDEEVREKAWETWMEVWLLWLEVRLRKGGGGSLEHHHHHH |
| C4-71-8x | MGPEEILERAKESLERAREASERGDEEEFRKAAEKALELAKRLVEQAKKEGDPELVLEAAKVALRVAELAAKNGDKEVFKKAAESALEVAKRLVEVASKEGDPELVLEAAKVALRVAELAAKNGDKEVFKKAAESALEVAKRLVEVASKEGDPELVLEAAKVALRVAELAAKNGDKEVFKKAAESALEVAKRLVEVASKEGDPELVLEAARVALEVARLAAENGDKEVFKKAAESALEVAKRLVEVASKEGDPWMVMWAALVALWVALLALRNGDKEVFKKAAESALEVAKRLVEVASKEGDPEMVLLAAWVALFVAWLAWLFGDKEVFKKAAESALEVAKRLVEVASKEGDPELVEEAAKVAEEVEKLAEKQGDEEVREKAWETWMEVWLLWLEVRLRKGGGGSLEHHHHHH |
| C6-71-6x | MGPEEILERAKESLERAREASERGDEEEFRKAAEKALELAKRLVEQAKKEGDPELVLEAAKVALRVAELAAKNGDKEVFKKAAESALEVAKRLVEVASKEGDPELVLEAAKVALEVAKLAFENGDKEVFKKAAESALEVAKRLVEVASKEGDPELVFEAARVALWVAWLAAWFGDKEVFKKAAESALEVAKRLVEVAKEEGDPELVLKAAFVALLVAIMAVILGDKEVFKKAAESALEVAKRLVEIAAREGDPELVEEAAKVAELVRELAKLMGDEEVYEKARETAREVRLFLLFVRIWEGGGGSLEHHHHHH |
| C6-71-8x | MGPEEILERAKESLERAREASERGDEEEFRKAAEKALELAKRLVEQAKKEGDPELVLEAAKVALRVAELAAKNGDKEVFKKAAESALEVAKRLVEVASKEGDPELVLEAAKVALRVAELAAKNGDKEVFKKAAESALEVAKRLVEVASKEGDPELVLEAAKVALEVAKLAFENGDKEVFKKAAESALEVAKRLVEVASKEGDPELVFEAARVALWVAWLAAWFGDKEVFKKAAESALEVAKRLVEVAKEEGDPELVLKAAFVALLVAIMAVILGDKEVFKKAAESALEVAKRLVEIAAREGDPELVEEAAKVAELVRELAKLMGDEEVYEKARETAREVRLFLLFVRIWEGGGGSLEHHHHHH |
| C8-71-6x | MGPEEILERAKESLERAREASERGDEEEFRKAAEKALELAKRLVEQAKKEGDPELVLEAAKVALRVAELAAKNGDKEVFKKAAESALEVAKRLVEVASKEGDPELVLEAAKVALEVARLAAENGDKEVFKKAAESALEVAKRLVEVASKEGDPELVLEAARVALWVAELAAKNGDKEVFKKAAESALEVAKRLVEVASKEGDPDLVAWAALVALWVAFLAFLNGDKEVFKKAAESALEVAKALMEVAMKVGAPWLVELAIAVARAVWLLAELFGDEEVRRRAEAFEIILRIAAIAVKAWLGGGGSLEHHHHHH |
| C8-71-8x | MGPEEILERAKESLERAREASERGDEEEFRKAAEKALELAKRLVEQAKKEGDPELVLEAAKVALRVAELAAKNGDKEVFKKAAESALEVAKRLVEVASKEGDPELVLEAAKVALRVAELAAKNGDKEVFKKAAESALEVAKRLVEVASKEGDPELVLEAAKVALEVARLAAENGDKEVFKKAAESALEVAKRLVEVASKEGDPELVLEAARVALWVAELAAKNGDKEVFKKAAESALEVAKRLVEVASKEGDPDLVAWAALVALWVAFLAFLNGDKEVFKKAAESALEVAKALMEVAMKVGAPWLVELAIAVARAVWLLAELFGDEEVRRRAEAFEIILRIAAIAVKAWLGGGGSLEHHHHHH |
| C2-58-2x_mb7 | MHHHHHHAENLYFQSGSDEELLRELLKLLLKLLEQMGDEEARRVVEELREELEKKGDPRALVLAFALVILVFLLRILRELGDEELVRRVEELWEELLKEGDPQAMMEVFKLVQELQERRGSGSDRRKEMDKVYRTAYKRITSTPDKEKRKEVVKEATEQLRRIAKDEEEKKKAAYMISFLKTLGS |
| C4-71C_mb7 | MGPEEILERARESLERAREASERGDEEEFRKAAEKALELAKRLVEQAKKEGDPWMVMWAALVALWVALLALRNGDKEVFKKAAESALEVAKRLVEVASKEGDPEMVLLAAWVALFVAWLAWLFGDKEVFKKAAESALEVAKRLVEVASKEGDPELVEEAAKVAEEVEKLAEKQGDEEVREKAWET |

| | |
|---|---|
| | WMEVWLLWLEVRLRKGGGGSDRRKEMDKVYRTAYKRITSTPDKEKRKEVVKEATEQLRRIAKDEEEKKKAAYMISFLKTLGSLEHHHHHH |
| C6-71C_mb7 | MGPEEILERAKESLERAKEAFERGDEEEFRKAAEKALELAKRLVEQAKKEGDPELVFEAARVALWVAWLAAWFGDKEVFKKAAESALEVAKRLVEVAKEEGDPELVLKAAFVALLVAIMAVILGDKEVFKKAAESALEVAKRLVEIAAREGDPELVEEAAKVAELVRELAKLMGDEEVYEKARETAREVRLFLLFVRIWEGGGGSDRRKEMDKVYRTAYKRITSTPDKEKRKEVVKEATEQLRRIAKDEEEKKKAAYMISFLKTLGSLEHHHHHH |
| C6-79C_mb7 | MGSSDEEEARELEERAREAAKRAIEAAKRTGDPRVRELAEELVKLAIWAAVEVWLDPSSSDVNEALKLIVEAIEAAVRALEAAERTGDPEVRELARELVRLAVEAAEEVQRNPSSSDVNEALKLIVIAIEAAVRALEAAERTGDPEVRELARELVRLAVEAAEEVQRNPSSEEVNEALRKIIKLILFAVMVLELAEEIGDPTWREMARRAVREAVELAEEVQRDPSGWLGHGSDRRKEMDKVYRTAYKRITSTPDKEKRKEVVKEATEQLRRIAKDEEEKKKAAYMISFLKTLGSLEHHHHHH |
| C8-71C_mb7 | MEKKIGPEEILERAKESLERAREASERGDEEEFRKAAEKALELAKRLVEQAKKEGDPELVLEAARVALWVAELAAKNGDKEVFKKAAESALEVAKRLVEVASKEGDPDLVAWAALVALWVAFLAFLNGDKEVFKKAAESALEVAKALMEVAMKVGAPWLVELAIAVARAVWLLAELFGDEEVRRRAEAFEIILRIAAIAVKAWLGGGGSDRRKEMDKVYRTAYKRITSTPDKEKRKEVVKEATEQLRRIAKDEEEKKKAAYMISFLKTLGSLEHHHHHH |
| C4-71N_mb7 | MGDRRKEMDKVYRTAYKRITSTPDKEKRKEVVKEATEQLRRIAKDEEEKKKAAYMISFLKTLGSPEEILERARESLERAREASERGDEEEFRKAAEKALELAKRLVEQAKKEGDPWMVMWAALVALWVALLALRNGDKEVFKKAAESALEVAKRLVEVASKEGDPEMVLLAAWVALFVAWLAWLFGDKEVFKKAAESALEVAKRLVEVASKEGDPELVEEAAKVAEEVEKLAEKQGDEEVREKAWETWMEVWLLWLEVRLRKGGGGSLEHHHHHH |
| C6-71N_mb7 | MGDRRKEMDKVYRTAYKRITSTPDKEKRKEVVKEATEQLRRIAKDEEEKKKAAYMISFLKTLGSPEEILERAKESLERAKEAFERGDEEEFRKAAEKALELAKRLVEQAKKEGDPELVFEAARVALWVAWLAAWFGDKEVFKKAAESALEVAKRLVEVAKEEGDPELVLKAAFVALLVAIMAVILGDKEVFKKAAESALEVAKRLVEIAAREGDPELVEEAAKVAELVRELAKLMGDEEVYEKARETAREVRLFLLFVRIWEGGGGSLEHHHHHH |
| C6-79N_mb7 | MGDRRKEMDKVYRTAYKRITSTPDKEKRKEVVKEATEQLRRIAKDEEEKKKAAYMISFLKTLGSSSDEEEARELEERAREAAKRAIEAAKRTGDPRVRELAEELVKLAIWAAVEVWLDPSSSDVNEALKLIVEAIEAAVRALEAAERTGDPEVRELARELVRLAVEAAEEVQRNPSSSDVNEALKLIVIAIEAAVRALEAAERTGDPEVRELARELVRLAVEAAEEVQRNPSSEEVNEALRKIIKLILFAVMVLELAEEIGDPTWREMARRAVREAVELAEEVQRDPSGWLGHGSLEHHHHHH |
| C8-71N_mb7 | MEKKIGDRRKEMDKVYRTAYKRITSTPDKEKRKEVVKEATEQLRRIAKDEEEKKKAAYMISFLKTLGSPEEILERAKESLERAREASERGDEEEFRKAAEKALELAKRLVEQAKKEGDPELVLEAARVALWVAELAAKNGDKEVFKKAAESALEVAKRLVEVASKEGDPDLVAWAALVALWVAFLAFLNGDKEVFKKAAESALEVAKALMEVAMKVGAPWLVELAIAVARAVWLLAELFGDEEVRRRAEAFEIILRIAAIAVKAWLGGGGSLEHHHHHH |

**Supplementary Table VI.** SEC-MALS data of oligomeric constructs

| Name | Expected MW (Da) | Measured MW (Da) |
|---|---|---|
| C4-18 | 106692 | 102200 |
| C4-717 | 94040 | 122500 |
| C6-714 | 141473 | 140000 |
| C6-46 | 140032 | 125900 |
| C4-131 | 90902 | 86880 |
| C4-81 | 108462 | 155800 |
| C6-79 | 161070 | 144600 |
| C4-71 | 96666 | 178400 |
| C4-71-6xRepeat | 138862 | 152800 |
| C4-71-8xRepeat | 181055 | 187700 |
| C6-71 | 143240 | 136900 |
| C6-71-6xRepeat | 206630 | 212400 |
| C6-71-8xRepeat | 269920 | 254400 |
| C8-71 | 187183 | 196300 |
| C8-71-6xRepeat | 271576 | 286400 |
| C8-71-8xRepeat | 355961 | 341800 |
| C2-58 | 26753 | 26360 |
| C2-CDX | 26166 | 33460 |
| C2-Y2D | 26258 | 24250 |

**Supplementary Table VII.** Putative markers for cell type identification from scRNA-seq data

| Cell Type | Markers |
|---|---|
| IPSCs | POU5F1 (OCT4) |
| | SOX2 |
| | MYC |
| | NANOG |

| | |
|---|---|
| Endothelial | PECAM1 (CD31) |
| | CDH5 (VE-Cadherin) |
| | TIE1 |
| | TEK (TIE2) |
| | VWF |
| | ACE |
| | KDR (VEGFR2) |
| | PDGF-B |
| Pericytes | PDGFR-B |
| | CSPG4 (NG2) |
| | ANGPT1 |
| | THY1 |
| | PRRX1 |
| | VCAM1 |
| | CD274 (PDL1) |
| Arterial Endothelial | NOTCH1 |
| | NOTCH4 |
| | NRP1 |
| | DLL4 |
| | HEY1/2 |
| | EFNB2 |
| Venous Endothelial | NRP2 |
| | EPHB4 |
| | NR2F2 (COUP-TFII) |
| Lymphatic Endothelial | SOX18 |
| | LYVE1 |
| | PDPN |

**Supplementary Table VIII.** Primer sequences used in RT-qPCR assays

| **Gene** | **Forward Sequence (5' → 3')** | **Reverse Sequence (5' → 3')** |
|---|---|---|
| B-actin | CACCATTGGCAATGAGCGGTTC | AGGTCTTTGCGGATGTCCACGT |
| FGFR1b | ACCAGTCTGCGTGGCTCACT | TGCCGGCCTCTCTTCCA |
| FGFR1c | AGGTGAACGGGAGTAAGATTGG | GTGCAGCACCTCCATCTCTTT |

**Supplementary Table IX.** Hash Oligonucleotides for single cell RNA-seq

| **Well** | **Name** | **Sequence** |
|---|---|---|
| A1 | sciPlex_1056 | GTCTCGTGGGCTCGGAGATGTGTATAAGAGACAGATGGTTACCGBAAAAAAAAAAAAAAAAAAAAAAAAAAAAAA |
| B1 | sciPlex_1057 | GTCTCGTGGGCTCGGAGATGTGTATAAGAGACAGCTTAGGTTATBAAAAAAAAAAAAAAAAAAAAAAAAAAAAAA |
| C1 | sciPlex_1058 | GTCTCGTGGGCTCGGAGATGTGTATAAGAGACAGCGCCTTAGCCBAAAAAAAAAAAAAAAAAAAAAAAAAAAAAA |
| D1 | sciPlex_1059 | GTCTCGTGGGCTCGGAGATGTGTATAAGAGACAGACGTAACGTABAAAAAAAAAAAAAAAAAAAAAAAAAAAAAA |
| E1 | sciPlex_1060 | GTCTCGTGGGCTCGGAGATGTGTATAAGAGACAGTTATTCTCTABAAAAAAAAAAAAAAAAAAAAAAAAAAAAAA |
| F1 | sciPlex_1061 | GTCTCGTGGGCTCGGAGATGTGTATAAGAGACAGGCATAGTATABAAAAAAAAAAAAAAAAAAAAAAAAAAAAAA |
| G1 | sciPlex_1062 | GTCTCGTGGGCTCGGAGATGTGTATAAGAGACAGCATGCGCATABAAAAAAAAAAAAAAAAAAAAAAAAAAAAAA |
| H1 | sciPlex_1063 | GTCTCGTGGGCTCGGAGATGTGTATAAGAGACAGGTACCAATTCBAAAAAAAAAAAAAAAAAAAAAAAAAAAAAA |
| A2 | sciPlex_1064 | GTCTCGTGGGCTCGGAGATGTGTATAAGAGACAGAACGAGATCABAAAAAAAAAAAAAAAAAAAAAAAAAAAAAA |
| B2 | sciPlex_1065 | GTCTCGTGGGCTCGGAGATGTGTATAAGAGACAGTCTCGATTAABAAAAAAAAAAAAAAAAAAAAAAAAAAAAAA |
| C2 | sciPlex_1066 | GTCTCGTGGGCTCGGAGATGTGTATAAGAGACAGCGTCGACCGGBAAAAAAAAAAAAAAAAAAAAAAAAAAAAAA |
| D2 | sciPlex_1067 | GTCTCGTGGGCTCGGAGATGTGTATAAGAGACAGGTTGATAGCTBAAAAAAAAAAAAAAAAAAAAAAAAAAAAAA |
| E2 | sciPlex_1068 | GTCTCGTGGGCTCGGAGATGTGTATAAGAGACAGCAGCGTTGGTBAAAAAAAAAAAAAAAAAAAAAAAAAAAAAA |
| F2 | sciPlex_1069 | GTCTCGTGGGCTCGGAGATGTGTATAAGAGACAGATCGGCATTGBAAAAAAAAAAAAAAAAAAAAAAAAAAAAAA |

| | | |
|---|---|---|
| G2 | sciPlex_1070 | GTCTCGTGGGCTCGGAGATGTGTATAAGAGACAGGGTAGTCCTABAAAAAAAAAAAAAAAAAAAAAAAAAAAAAA |
| H2 | sciPlex_1071 | GTCTCGTGGGCTCGGAGATGTGTATAAGAGACAGTAGCATCGCGBAAAAAAAAAAAAAAAAAAAAAAAAAAAAAA |
| A3 | sciPlex_1072 | GTCTCGTGGGCTCGGAGATGTGTATAAGAGACAGACTGGTAACCBAAAAAAAAAAAAAAAAAAAAAAAAAAAAAA |
| B3 | sciPlex_1073 | GTCTCGTGGGCTCGGAGATGTGTATAAGAGACAGTCTACTGACCBAAAAAAAAAAAAAAAAAAAAAAAAAAAAAA |
| C3 | sciPlex_1074 | GTCTCGTGGGCTCGGAGATGTGTATAAGAGACAGCAAGAGTTATBAAAAAAAAAAAAAAAAAAAAAAAAAAAAAA |
| D3 | sciPlex_1075 | GTCTCGTGGGCTCGGAGATGTGTATAAGAGACAGCAATTAGGAABAAAAAAAAAAAAAAAAAAAAAAAAAAAAAA |
| E3 | sciPlex_1076 | GTCTCGTGGGCTCGGAGATGTGTATAAGAGACAGTTGCGGAACCBAAAAAAAAAAAAAAAAAAAAAAAAAAAAAA |
| F3 | sciPlex_1077 | GTCTCGTGGGCTCGGAGATGTGTATAAGAGACAGAGCCGAAGTABAAAAAAAAAAAAAAAAAAAAAAAAAAAAAA |
| G3 | sciPlex_1078 | GTCTCGTGGGCTCGGAGATGTGTATAAGAGACAGGGTCCGTACGBAAAAAAAAAAAAAAAAAAAAAAAAAAAAAA |
| H3 | sciPlex_1079 | GTCTCGTGGGCTCGGAGATGTGTATAAGAGACAGAGCATGCAAGBAAAAAAAAAAAAAAAAAAAAAAAAAAAAAA |
| A4 | sciPlex_1080 | GTCTCGTGGGCTCGGAGATGTGTATAAGAGACAGTCTTCGTTGCBAAAAAAAAAAAAAAAAAAAAAAAAAAAAAA |
| B4 | sciPlex_1081 | GTCTCGTGGGCTCGGAGATGTGTATAAGAGACAGGTTGGAAGAABAAAAAAAAAAAAAAAAAAAAAAAAAAAAAA |
| C4 | sciPlex_1082 | GTCTCGTGGGCTCGGAGATGTGTATAAGAGACAGGGAATACGGABAAAAAAAAAAAAAAAAAAAAAAAAAAAAAA |
| D4 | sciPlex_1083 | GTCTCGTGGGCTCGGAGATGTGTATAAGAGACAGACCGACGGTABAAAAAAAAAAAAAAAAAAAAAAAAAAAAAA |
| E4 | sciPlex_1084 | GTCTCGTGGGCTCGGAGATGTGTATAAGAGACAGATACTTGATTBAAAAAAAAAAAAAAAAAAAAAAAAAAAAAA |
| F4 | sciPlex_1085 | GTCTCGTGGGCTCGGAGATGTGTATAAGAGACAGGTCGTTCGCCBAAAAAAAAAAAAAAAAAAAAAAAAAAAAAA |
| G4 | sciPlex_1086 | GTCTCGTGGGCTCGGAGATGTGTATAAGAGACAGAATGATTGCABAAAAAAAAAAAAAAAAAAAAAAAAAAAAAA |
| H4 | sciPlex_1087 | GTCTCGTGGGCTCGGAGATGTGTATAAGAGACAGCCATTAGCAGBAAAAAAAAAAAAAAAAAAAAAAAAAAAAAA |
| A5 | sciPlex_1088 | GTCTCGTGGGCTCGGAGATGTGTATAAGAGACAGCTGACTAATGBAAAAAAAAAAAAAAAAAAAAAAAAAAAAAA |
| B5 | sciPlex_1089 | GTCTCGTGGGCTCGGAGATGTGTATAAGAGACAGCCTTACGGACBAAAAAAAAAAAAAAAAAAAAAAAAAAAAAA |

| | | |
|---|---|---|
| C5 | sciPlex_1090 | GTCTCGTGGGCTCGGAGATGTGTATAAGAGACAGCGCATAACTABAAAAAAAAAAAAAAAAAAAAAAAAAAAAAAA |
| D5 | sciPlex_1091 | GTCTCGTGGGCTCGGAGATGTGTATAAGAGACAGAACCGGAGGABAAAAAAAAAAAAAAAAAAAAAAAAAAAAAAA |
| E5 | sciPlex_1092 | GTCTCGTGGGCTCGGAGATGTGTATAAGAGACAGAATGCAGCGGBAAAAAAAAAAAAAAAAAAAAAAAAAAAAAAA |
| F5 | sciPlex_1093 | GTCTCGTGGGCTCGGAGATGTGTATAAGAGACAGCAGTCGGCAABAAAAAAAAAAAAAAAAAAAAAAAAAAAAAAA |
| G5 | sciPlex_1094 | GTCTCGTGGGCTCGGAGATGTGTATAAGAGACAGATAGAATCCABAAAAAAAAAAAAAAAAAAAAAAAAAAAAAAA |
| H5 | sciPlex_1095 | GTCTCGTGGGCTCGGAGATGTGTATAAGAGACAGTCTCAGATATBAAAAAAAAAAAAAAAAAAAAAAAAAAAAAAA |
| A6 | sciPlex_1096 | GTCTCGTGGGCTCGGAGATGTGTATAAGAGACAGACGAAGTATTBAAAAAAAAAAAAAAAAAAAAAAAAAAAAAAA |
| B6 | sciPlex_1097 | GTCTCGTGGGCTCGGAGATGTGTATAAGAGACAGAGACTTATCABAAAAAAAAAAAAAAAAAAAAAAAAAAAAAAA |
| C6 | sciPlex_1098 | GTCTCGTGGGCTCGGAGATGTGTATAAGAGACAGTCGCGCCGTABAAAAAAAAAAAAAAAAAAAAAAAAAAAAAAA |
| D6 | sciPlex_1099 | GTCTCGTGGGCTCGGAGATGTGTATAAGAGACAGAGCTTGAAGABAAAAAAAAAAAAAAAAAAAAAAAAAAAAAAA |
| E6 | sciPlex_1100 | GTCTCGTGGGCTCGGAGATGTGTATAAGAGACAGCGGTAGCTACBAAAAAAAAAAAAAAAAAAAAAAAAAAAAAAA |
| F6 | sciPlex_1101 | GTCTCGTGGGCTCGGAGATGTGTATAAGAGACAGGCGGCATGCGBAAAAAAAAAAAAAAAAAAAAAAAAAAAAAAA |
| G6 | sciPlex_1102 | GTCTCGTGGGCTCGGAGATGTGTATAAGAGACAGAAGACTGGCTBAAAAAAAAAAAAAAAAAAAAAAAAAAAAAAA |
| H6 | sciPlex_1103 | GTCTCGTGGGCTCGGAGATGTGTATAAGAGACAGCGCATTCTTABAAAAAAAAAAAAAAAAAAAAAAAAAAAAAAA |
| A7 | sciPlex_1104 | GTCTCGTGGGCTCGGAGATGTGTATAAGAGACAGTACCGTCTCCBAAAAAAAAAAAAAAAAAAAAAAAAAAAAAAA |
| B7 | sciPlex_1105 | GTCTCGTGGGCTCGGAGATGTGTATAAGAGACAGAATATAGTATBAAAAAAAAAAAAAAAAAAAAAAAAAAAAAAA |
| C7 | sciPlex_1106 | GTCTCGTGGGCTCGGAGATGTGTATAAGAGACAGATCATAAGATBAAAAAAAAAAAAAAAAAAAAAAAAAAAAAAA |
| D7 | sciPlex_1107 | GTCTCGTGGGCTCGGAGATGTGTATAAGAGACAGATCAATATCCBAAAAAAAAAAAAAAAAAAAAAAAAAAAAAAA |
| E7 | sciPlex_1108 | GTCTCGTGGGCTCGGAGATGTGTATAAGAGACAGTATTACCAACBAAAAAAAAAAAAAAAAAAAAAAAAAAAAAAA |
| F7 | sciPlex_1109 | GTCTCGTGGGCTCGGAGATGTGTATAAGAGACAGGAGAAGACCABAAAAAAAAAAAAAAAAAAAAAAAAAAAAAAA |

| | | |
|---|---|---|
| G7 | sciPlex_1110 | GTCTCGTGGGCTCGGAGATGTGTATAAGAGACAGTGGCGCTCTCBAAAAAAAAAAAAAAAAAAAAAAAAAAAAAA |
| H7 | sciPlex_1111 | GTCTCGTGGGCTCGGAGATGTGTATAAGAGACAGGCGACGATAABAAAAAAAAAAAAAAAAAAAAAAAAAAAAAA |
| A8 | sciPlex_1112 | GTCTCGTGGGCTCGGAGATGTGTATAAGAGACAGAACGTCGCGABAAAAAAAAAAAAAAAAAAAAAAAAAAAAAA |
| B8 | sciPlex_1113 | GTCTCGTGGGCTCGGAGATGTGTATAAGAGACAGATGAAGCTTCBAAAAAAAAAAAAAAAAAAAAAAAAAAAAAA |
| C8 | sciPlex_1114 | GTCTCGTGGGCTCGGAGATGTGTATAAGAGACAGGCCATAGAGTBAAAAAAAAAAAAAAAAAAAAAAAAAAAAAA |
| D8 | sciPlex_1115 | GTCTCGTGGGCTCGGAGATGTGTATAAGAGACAGTGACTGAGAABAAAAAAAAAAAAAAAAAAAAAAAAAAAAAA |
| E8 | sciPlex_1116 | GTCTCGTGGGCTCGGAGATGTGTATAAGAGACAGCGGCTCCGAABAAAAAAAAAAAAAAAAAAAAAAAAAAAAAA |
| F8 | sciPlex_1117 | GTCTCGTGGGCTCGGAGATGTGTATAAGAGACAGCAGCCAATTGBAAAAAAAAAAAAAAAAAAAAAAAAAAAAAA |
| G8 | sciPlex_1118 | GTCTCGTGGGCTCGGAGATGTGTATAAGAGACAGGCGACCAGTTBAAAAAAAAAAAAAAAAAAAAAAAAAAAAAA |
| H8 | sciPlex_1119 | GTCTCGTGGGCTCGGAGATGTGTATAAGAGACAGCCTAACGACGBAAAAAAAAAAAAAAAAAAAAAAAAAAAAAA |
| A9 | sciPlex_1120 | GTCTCGTGGGCTCGGAGATGTGTATAAGAGACAGCTTCGCAATCBAAAAAAAAAAAAAAAAAAAAAAAAAAAAAA |
| B9 | sciPlex_1121 | GTCTCGTGGGCTCGGAGATGTGTATAAGAGACAGTGACTCCGTTBAAAAAAAAAAAAAAAAAAAAAAAAAAAAAA |
| C9 | sciPlex_1122 | GTCTCGTGGGCTCGGAGATGTGTATAAGAGACAGGATAGTCGCTBAAAAAAAAAAAAAAAAAAAAAAAAAAAAAA |
| D9 | sciPlex_1123 | GTCTCGTGGGCTCGGAGATGTGTATAAGAGACAGAAGGTACTAABAAAAAAAAAAAAAAAAAAAAAAAAAAAAAA |
| E9 | sciPlex_1124 | GTCTCGTGGGCTCGGAGATGTGTATAAGAGACAGTTGCATGAGGBAAAAAAAAAAAAAAAAAAAAAAAAAAAAAA |
| F9 | sciPlex_1125 | GTCTCGTGGGCTCGGAGATGTGTATAAGAGACAGGTATTATATABAAAAAAAAAAAAAAAAAAAAAAAAAAAAAA |
| G9 | sciPlex_1126 | GTCTCGTGGGCTCGGAGATGTGTATAAGAGACAGATTCTTGGCTBAAAAAAAAAAAAAAAAAAAAAAAAAAAAAA |
| H9 | sciPlex_1127 | GTCTCGTGGGCTCGGAGATGTGTATAAGAGACAGTGCATCTTGGBAAAAAAAAAAAAAAAAAAAAAAAAAAAAAA |
| A10 | sciPlex_1128 | GTCTCGTGGGCTCGGAGATGTGTATAAGAGACAGGTTGGCTCAABAAAAAAAAAAAAAAAAAAAAAAAAAAAAAA |
| B10 | sciPlex_1129 | GTCTCGTGGGCTCGGAGATGTGTATAAGAGACAGAATATCATTABAAAAAAAAAAAAAAAAAAAAAAAAAAAAAA |

| | | |
|---|---|---|
| C10 | sciPlex_1130 | GTCTCGTGGGCTCGGAGATGTGTATAAGAGACAGTAACTAAGTCBAAAAAAAAAAAAAAAAAAAAAAAAAAAAAA |
| D10 | sciPlex_1131 | GTCTCGTGGGCTCGGAGATGTGTATAAGAGACAGCAATAACCAABAAAAAAAAAAAAAAAAAAAAAAAAAAAAAA |
| E10 | sciPlex_1132 | GTCTCGTGGGCTCGGAGATGTGTATAAGAGACAGTTATACTGCABAAAAAAAAAAAAAAAAAAAAAAAAAAAAAA |
| F10 | sciPlex_1133 | GTCTCGTGGGCTCGGAGATGTGTATAAGAGACAGTGAGCAGAGCBAAAAAAAAAAAAAAAAAAAAAAAAAAAAAA |
| G10 | sciPlex_1134 | GTCTCGTGGGCTCGGAGATGTGTATAAGAGACAGTGCAAGCCAABAAAAAAAAAAAAAAAAAAAAAAAAAAAAAA |
| H10 | sciPlex_1135 | GTCTCGTGGGCTCGGAGATGTGTATAAGAGACAGTGGAGAACGABAAAAAAAAAAAAAAAAAAAAAAAAAAAAAA |
| A11 | sciPlex_1136 | GTCTCGTGGGCTCGGAGATGTGTATAAGAGACAGATCGGATTCABAAAAAAAAAAAAAAAAAAAAAAAAAAAAAA |
| B11 | sciPlex_1137 | GTCTCGTGGGCTCGGAGATGTGTATAAGAGACAGACTAGACCGABAAAAAAAAAAAAAAAAAAAAAAAAAAAAAA |
| C11 | sciPlex_1138 | GTCTCGTGGGCTCGGAGATGTGTATAAGAGACAGCGAGATGCTTBAAAAAAAAAAAAAAAAAAAAAAAAAAAAAA |
| D11 | sciPlex_1139 | GTCTCGTGGGCTCGGAGATGTGTATAAGAGACAGTTCTATTAATBAAAAAAAAAAAAAAAAAAAAAAAAAAAAAA |
| E11 | sciPlex_1140 | GTCTCGTGGGCTCGGAGATGTGTATAAGAGACAGAATTAGTCCABAAAAAAAAAAAAAAAAAAAAAAAAAAAAAA |
| F11 | sciPlex_1141 | GTCTCGTGGGCTCGGAGATGTGTATAAGAGACAGGCTCCAAGCCBAAAAAAAAAAAAAAAAAAAAAAAAAAAAAA |
| G11 | sciPlex_1142 | GTCTCGTGGGCTCGGAGATGTGTATAAGAGACAGCTCTTCCTAABAAAAAAAAAAAAAAAAAAAAAAAAAAAAAA |
| H11 | sciPlex_1143 | GTCTCGTGGGCTCGGAGATGTGTATAAGAGACAGCCGCGTTAACBAAAAAAAAAAAAAAAAAAAAAAAAAAAAAA |
| A12 | sciPlex_1144 | GTCTCGTGGGCTCGGAGATGTGTATAAGAGACAGGACGGAATAGBAAAAAAAAAAAAAAAAAAAAAAAAAAAAAA |
| B12 | sciPlex_1145 | GTCTCGTGGGCTCGGAGATGTGTATAAGAGACAGCTCAGAGTTGBAAAAAAAAAAAAAAAAAAAAAAAAAAAAAA |
| C12 | sciPlex_1146 | GTCTCGTGGGCTCGGAGATGTGTATAAGAGACAGGGACGTATGABAAAAAAAAAAAAAAAAAAAAAAAAAAAAAA |
| D12 | sciPlex_1147 | GTCTCGTGGGCTCGGAGATGTGTATAAGAGACAGAAGATGAGTCBAAAAAAAAAAAAAAAAAAAAAAAAAAAAAA |
| E12 | sciPlex_1148 | GTCTCGTGGGCTCGGAGATGTGTATAAGAGACAGATGCGCTACCBAAAAAAAAAAAAAAAAAAAAAAAAAAAAAA |
| F12 | sciPlex_1149 | GTCTCGTGGGCTCGGAGATGTGTATAAGAGACAGGGACGCCTAABAAAAAAAAAAAAAAAAAAAAAAAAAAAAAA |

| | | |
|---|---|---|
| G12 | sciPlex_1150 | GTCTCGTGGGCTCGGAGATGTGTATAAGAGACAGCAGTTAGACCBAAAAAAAAAAAAAAAAAAAAAAAAAAAAAA |
| H12 | sciPlex_1151 | GTCTCGTGGGCTCGGAGATGTGTATAAGAGACAGCCGTCTCAATBAAAAAAAAAAAAAAAAAAAAAAAAAAAAAA |
